# An expanded structural and biochemical map of GLI-SUFU recognition identifies binding-defective disease variants

**DOI:** 10.64898/2026.01.06.698042

**Authors:** A. Jane Bardwell, Umaima Arif, Alex Salgado, Fatima Muhammad, Celeste N. Salinas Auris, Lee Bardwell

## Abstract

The tumor suppressor SUFU is a key negative regulator of oncogenic GLI transcription factors. *SUFU* mutations predispose to medulloblastoma and can cause Gorlin and Joubert syndromes. Limited structural information is available on the GLI-SUFU complex, contributing to the fact that the vast majority of *SUFU* missense variants sequenced from affected patients are of unknown clinical significance. Using AlphaFold3 and complementary computational analysis, we predicted structures of the three human GLI-SUFU complexes and the orthologous *Drosophila* Ci-Sufu complex, revealing previously unrecognized contact interfaces. By site-directed mutagenesis and GLI-SUFU binding assays, we validated the importance of key interface residues, many of which are mutated in familial medulloblastoma and Gorlin syndrome. These findings provide new structure-function insights into a signaling complex mutated in cancer and rare congenital diseases, and offer an expanded framework for interpreting clinically observed *SUFU* variants and mechanisms that tune SUFU repression.

**Teaser:** AI-powered modeling plus experimental validation reveal new GLI–SUFU interfaces linked to cancer and congenital mutations.

## INTRODUCTION

In humans and model organisms, the Hedgehog (HH) signaling pathway drives the proliferation, migration, and differentiation of stem/progenitor cells during embryonic patterning and adult tissue homeostasis, regeneration and repair (*1–3*). Defective HH signaling results in human birth defects, most notably affecting the face, the limbs, and the nervous system (*4–6*). Mutations that constitutively activate the HH pathway cause basal cell carcinoma, the most frequent cancer in the U.S. and many other parts of the world, as well as the rarer yet more aggressive skin tumor basosquamous carcinoma (*7, 8*). Moreover, activating mutations in the HH pathway are responsible for ∼30% of medulloblastomas, the most frequent brain cancer in infants and young children (*9*). HH pathway mutations and HH signaling in the tumor microenvironment are also thought to contribute to tumor progression in many other cancers (*10*).

Physiological HH signaling typically proceeds via a paracrine mechanism: HH ligand secreted by a nearby cell diffuses through the extracellular space before binding to a cell surface receptor on the target cell. Ligand-receptor binding activates a complicated signaling cascade that involves phosphorylation /dephosphorylation, dynamic protein-protein interactions, proteolytic processing, and trafficking into/out of the primary cilium. Despite tremendous progress, these events are still incompletely understood (*11–14*).

In mammals and most other vertebrates, three zinc-finger-type DNA-binding transcription factors – GLI1, GLI2 and GLI3 – are the transcriptional effectors of the HH pathway. GLI1 and GLI2 function mainly as transactivators while GLI3 functions mainly as a repressor. Targets for GLI activators in different cell types include pro-proliferative and anti-apoptotic genes, developmental regulators, and genes involved in the epithelial-to-mesenchyme transition (*10, 15–17*).

A 54 kilodalton protein named SUFU is a key regulator of the GLI proteins in mammalian cells. SUFU physically interacts with the three GLI transcription factors with low nanomolar affinity. In unstimulated cells, SUFU promotes the processing of GLI2 and GLI3 into truncated repressor forms, and restrains full-length GLI molecules from entering the nucleus, while also protecting them from degradation. Upon stimulation with Hedgehog ligand, GLI-SUFU complexes travel to the tip of the primary cilium, where the GLI proteins become fully activated by a series of phosphorylation events that are still being worked out, leading to full or partial dissociation from SUFU. Subsequently, active GLI1 and GLI2 enter the nucleus and bind to DNA, perhaps also accompanied by SUFU, or reengaging with it there (*12, 18–21*). SUFU also binds to and regulates the GLIS3 transcription factor (*22*). GLIS3 is not involved in Hedgehog signaling; it is a pioneer factor in the development of the pancreas, kidney and thyroid. Mutations in GLIS3 or in its binding motif in the insulin promoter are associated with neonatal diabetes (*23–25*).

Consistent with its role as a crucial negative regulator of HH signaling, SUFU is recognized as a tumor suppressor. Germline and somatic loss-of-function mutations in *SUFU* predispose to medulloblastoma, a highly malignant pediatric brain tumor that is both the most lethal and most common childhood tumor of the central nervous system (*26–32*). *SUFU* mutations are also found in patients with the rare congenital disorders Gorlin syndrome and Joubert syndrome (*33–37*). Adolescent and adult patients with inherited *SUFU* mutations have an increased risk of developing other brain tumors such as meningioma, as well as skin neoplasms such as basal cell carcinoma (*38–44*). Somatic *SUFU* mutations have also been found in various cancers, including sporadic basal cell carcinomas, lung tumors, and others (*19, 45–48*). Moreover, *GLI1* or *GLI2* gene amplifications, fusions or other forms of overexpression are often seen in human cancers and rodent preclinical models, often in the same spectrum of tumors in which *SUFU* mutations are known to play a role (*17, 29, 32, 49–59*). *GLI* amplifications or *SUFU* mutations can also promote resistance to clinical HH-pathway inhibitors such as vismodegib and sonidegib (*60–62*). These observations highlight the central role of an aberrant GLI-SUFU interaction in the pathology of multiple diseases.

Although the first animal *GLI* gene was cloned from a human glioma (*63*), many other components of the Hedgehog pathway were first discovered and characterized in classic experiments performed in the fruit fly *Drosophila melanogaster* (reviewed in (*64*)), and studies in this organism continue to drive the field (*65*). Of note for this study, *Drosophila* has a single GLI ortholog named Ci, and a SUFU ortholog, referred to herein as dSufu for clarity.

X-ray crystal structures of near-full length fly and human SUFU have been solved (*66, 67*), but much more limited information is available on the structure of the GLI proteins or of the GLI-SUFU complex. In 2003, a short, high-affinity binding site for SUFU, ^120^SYGHLS^125^, was identified in the N-terminal portion of GLI1 (*68*). In 2013, two independent groups published structures of SUFU bound to short GLI1 or GLI3-derived peptides containing this motif. In these structures, the peptide forms a β-strand that binds in a narrow channel between the N-terminal and C-terminal lobes of the bi-lobed SUFU structure (*66, 67*).

AlphaFold3 (AF3) is a groundbreaking AI model that can predict the 3D structures of biomolecular complexes (e.g., protein–protein, protein–DNA/RNA, and protein–ligand). It uses a diffusion approach, gradually refining an initially randomized 3D configuration into a final structure of the complex, guided by patterns learned from training data. AlphaFold3 was trained on a mixture of experimentally-determined macromolecular structures from the Protein Data Bank (PDB) and on large “distillation” sets, where earlier AlphaFold models generated structures for diverse protein sequences to broaden the training signal beyond what is available experimentally. These included sets designed to better handle disordered protein regions. During prediction, AF3 also leverages evolutionary covariation information from multiple sequence alignments built by searching large sequence databases (*69, 70*).

Here we used AlphaFold3 and other newly available tools to predict and analyze the structures of the human GLI1-SUFU, GLI2-SUFU, and GLI3-SUFU complexes, obtaining high-confidence predictions. We also predicted the structure of the orthologous fruit fly Ci-dSufu complex and the homologous human GLIS3-SUFU complex. The predicted structures revealed three novel interfaces between GLI2/GLI3 and SUFU, two of which are also present in the predicted GLI1-SUFU and Ci-dSufu structures. We then performed a series of biochemical experiments to interrogate important features of the predicted structures. First, we experimentally validated both the ability of the novel ‘SIC domain’ in human GLI proteins to bind to SUFU and the importance of key interface residues in this domain and in the N-terminus of SUFU. Next, by targeted mutagenesis of GLI1, we validated the importance of key interface residues in the novel ‘RSSL motif’ for SUFU binding. Finally, focusing on predicted contact residues that are mutated in medulloblastoma and Gorlin syndrome families and sporadic tumors, we tested the effect of an extensive panel of additional SUFU residues on GLI-SUFU binding, revealing their crucial role in GLI-SUFU complex formation.

## RESULTS

### AlphaFold3 prediction workflow

To begin this study, we used the AlphaFold3 server to investigate the predicted structure of the complex between full-length human GLI1, GLI2 or GLI3 proteins (1106, 1586 and 1580 residues in length, respectively) and full-length human SUFU (484 residues). Five zinc ions were included in each run to coordinate with the cysteine and histidine residues in the 5 zinc fingers of the GLI proteins. A 20-base pair oligonucleotide containing the GLI DNA-binding motif was also added to some runs. For comparison, we also ran predictions in the absence of various components, including GLI or SUFU.

AlphaFold3’s diffusion model involves randomness, so each server run is associated with a random seed that controls the initial noisy starting configuration and the stochastic choices made during iterative refinement. Thus, for each protein pair analyzed, we ran at least five replicate (“cloned”) jobs, each containing the same input sequences but initiated with a different random seed. For each job, AF3 produces five predicted structures by sampling the diffusion process five times; hence our pipeline resulted in (at least) 25 models for each protein pair (5 seeds x 5 diffusion samples). The job results were sorted by model confidence, and the best model from each job was chosen for further analysis, yielding a set of ‘top five’ Computed Structure Models (CSMs) – each resulting from a different job initiated with a distinct random seed – for every protein pair. This allowed us to identify the elements of the predicted structures that were reproducibly produced with different random seeds. All results discussed in this study were reproducibly observed across the top models for each interaction.

### Overview of novel GLI-SUFU interfaces predicted by AlphaFold3

Using AlphaFold3 to co-fold full-length human GLI1 with full-length human SUFU resulted in the formation of two distinct GLI-SUFU interfaces, which were also found in AF3-predicted GLI2-SUFU and GLI3-SUFU structures, as well as in the predicted structure of the complex of *D. melanogaster* SUFU protein (dSufu) with the fly GLI ortholog Ci (Figure 1, Figure 2, Supplemental Movies 1-4). A third interface was seen in the GLI2- and GLI3-SUFU structures only. These novel interfaces are briefly described in the next several paragraphs, and then in more detail subsequently.

**Figure 1.**
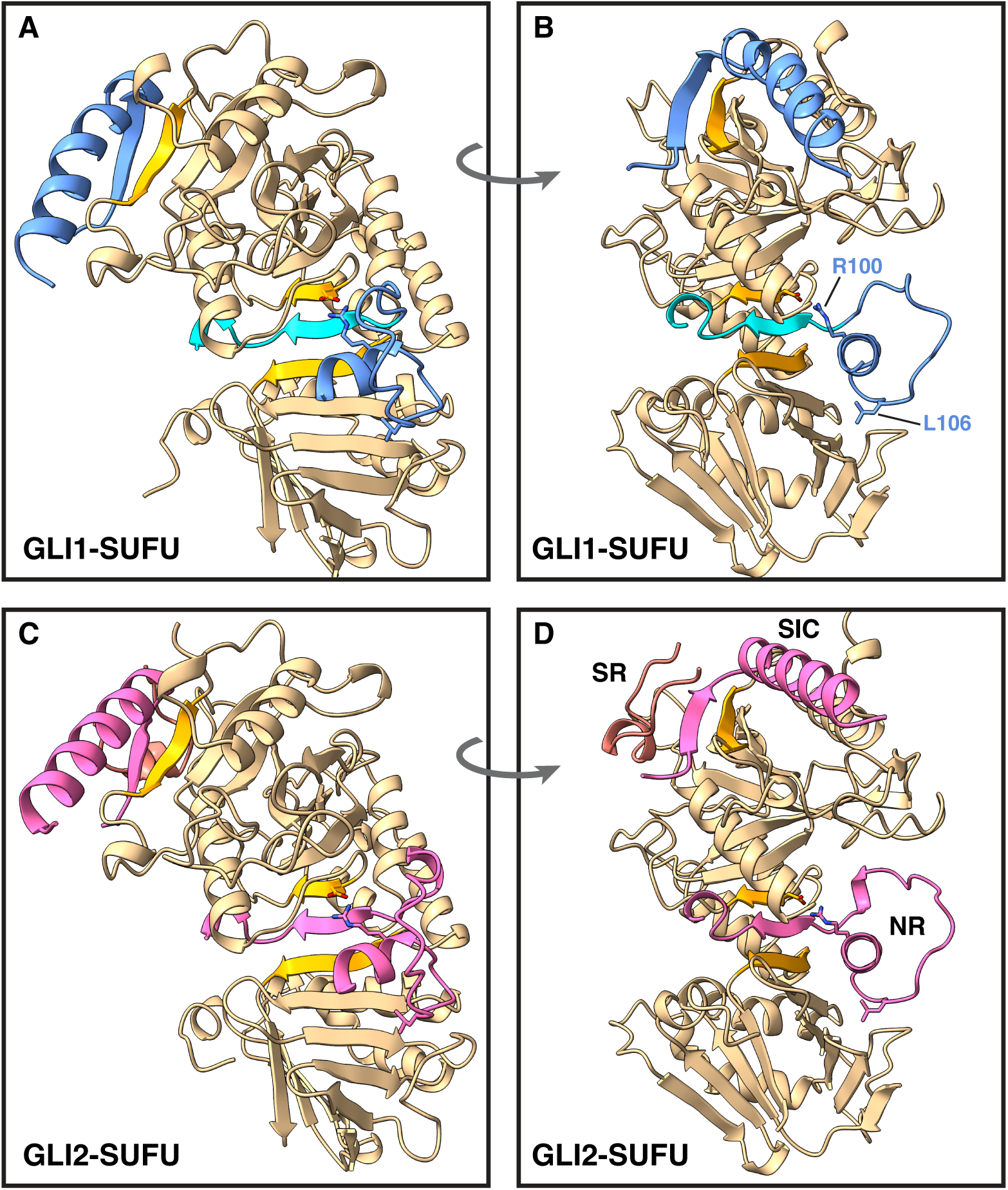
Predicted structures of human GLI1-SUFU and GLI2-SUFU complexes. **A** Predicted GLI1-SUFU complex, “front” view. SUFU (colored tan with β strands 1, 5 and 9 colored orange) is shown with the N-terminal domain at the top of each picture and the C-terminal lobe at the bottom; SUFU IDR1 and IDR2 are not shown; see text for details. The side chain of SUFU Asp159 is shown. The SUFU-binding domains of GLI1 (NR domain C-terminal region, residues 95-128; SIC domain, residues 1081-1106) are shown, with side chains visible for GLI1 Arg100 and Leu106. GLI1 is colored blue, with the SYGHLS motif (residues 119-128), whose structure was previously determined by x-ray crystallography, colored cyan. Model is CSM 261#, GLI1 5z SUFU. **B** GLI1-SUFU complex, “side” view. As in **A**, but the structure is rotated approximately 50 degrees (ccw) around the y axis and -15 around the x axis. **C** Predicted GLI2-SUFU complex, front view. The GLI2 NR domain C-terminal region (residues 245-279) and SIC domain (1561-1586) are colored pink; the SR motif (residues 61-75) is colored rose. Side chains are visible for SUFU Asp159 and GLI2 Arg250 and Leu256. Model is CSM 92#, GLI2 5z SUFU. **D** GLI2-SUFU complex, side view. The SR, SIC and NR domains are labeled.

**Figure 2.**
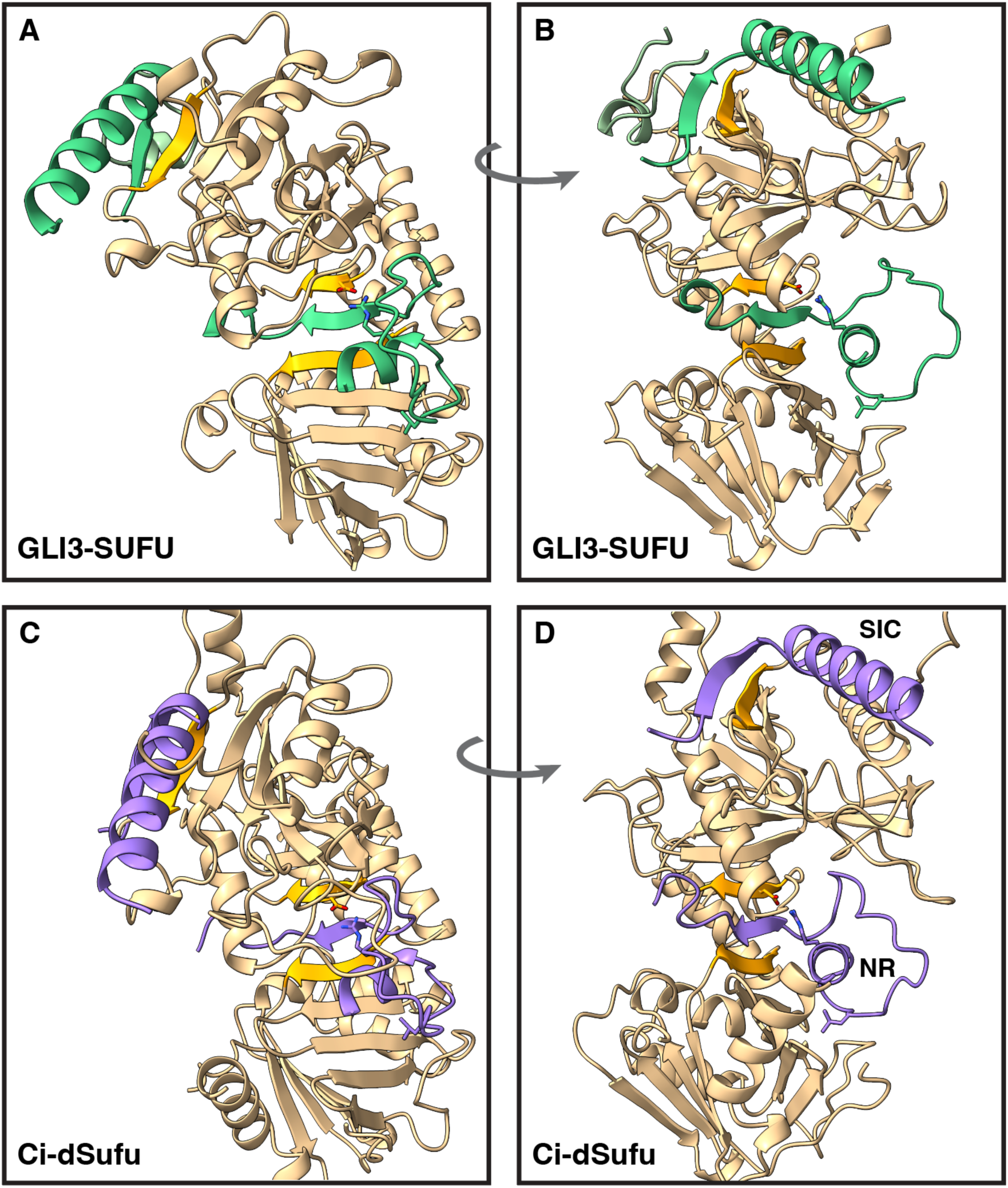
Predicted structures of human GLI3-SUFU and fly Ci-dSufu complexes. **A** Predicted GLI3-SUFU complex, front view. SUFU is colored as in Figure 1, with the side chain of Asp159 shown. The SUFU-binding domains of GLI3 (SR motif, residues 134-148; NR domain C-terminal region, residues 306-340; SIC domain, residues 1554-1580) are shown in green, with side chains visible for GLI3 Arg311 and Leu317. Model is CSM 248#. **B** GLI3-SUFU complex, side view. **C** Predicted *D. melanogaster* Ci-dSufu complex, front view. The side chain of dSufu Asp154 is shown. The dSufu-binding domains of Ci (NR domain C-terminal region, residues 226-263; SIC domain, residues 1371-1397) are shown in purple, with side chains visible for Ci Arg233 and Leu239. Model is CSM 228#. **D** Ci-dSufu complex, side view. The SIC and NR domains are labeled.

The first interface, herein designated the ‘RSSL motif’ after its most conserved residues, consists of a stretch of approximately 13 amino acids, which are LQTVI<u>R</u>T<u>S</u>PS<u>SL</u>V in GLI1. The first half of this motif forms a short α-helix from which protruding side chains contact SUFU; the second half forms an extended structure that docks into a complementary groove and hydrophobic pocket in SUFU. The RSSL motif makes extensive contacts with the C-terminal lobe of SUFU, particularly with residues in SUFU beta strands β13, β14, β16 and connecting loops. The RSSL motif also sits over the SYGHLS motif and shields it from solvent. The SYGHLS motif, for its part, resides in virtually the same position in all four AF3-predicted structures as it does in the published co-crystal structures of SUFU with GLI1 or GLI3 SYGHLS peptides: sandwiched between SUFU strands β5 and β9. The RSSL motif is connected to the SYGHLS motif by an 11-13 residue unstructured loop that does not make any contacts with SUFU. The RSSL motif and the SYGHLS motif form a combined, conditionally-folded unit hereafter referred to as the NR (N-terminal Regulatory) domain in accord with previous literature (*71*). The structure of the NR domain and its interface with SUFU are described in greater detail further below (Figures 4 and 5); in addition, the importance of many novel GLI-SUFU contact residues is experimentally tested and verified (Figures 7–10).

The second SUFU-binding interface seen in all three human GLI paralogs and in Ci – hereafter referred to as the SUFU-interacting C-terminal (SIC) domain in accord with previous studies (*72*) – consists of the C-terminal ∼27 residues of the GLI/Ci proteins. This region folds into a β-strand and α-helix and uses both elements to bind to the N-terminal domain of SUFU. In particular, the SIC β-strand binds to SUFU β1 by strand addition, while the amphipathic α-helix inserts hydrophobic residues into a hydrophobic docking groove in SUFU. The structure of the SIC domain is described in greater detail further below; we also experimentally demonstrate that the SIC domain is sufficient to bind SUFU *in vitro* and validate the importance of predicted GLI/SUFU interface residues for this interaction (Figure 6).

A third GLI-SUFU interface is present in the GLI2-SUFU and GLI3-SUFU structures, but not in the GLI1-SUFU or Ci-dSufu structures. This interface involves a stretch of 15 amino acids in the repressor domains of GLI2 and GLI3, which interacts both with the folded SIC domain and with residues in the N-terminus of SUFU. In this work, we have provisionally named this motif the SR (SUFU-binding, repressor domain) motif.

### Most of the GLI protein sequence is predicted to be intrinsically disordered

The structure of the 5-zinc finger DNA-binding domain (5ZF domain) of GLI1 bound to a 20 base pair DNA oligonucleotide was solved in 1993 (*73*). There is excellent structural superposition of the predicted 5ZF domain in AF3-generated GLI-SUFU structures with the 1993 x-ray structure, particularly when the DNA 20-mer is included in the AF3 job (Figure S1). The spatial position of the 5ZF domain relative to SUFU varied in different GLI-SUFU cofolding jobs, however, indicating that AF3 is unsure of their relative positioning (Figure S2). Furthermore, there are no contacts predicted between the 5ZF domain and SUFU, or between the 5ZF domain and the NR, SIC or SR domains in GLI. This is true both in the presence and absence of the GLI DNA-binding motif oligonucleotide. Hence, the 5ZF domain and the five bound zinc residues are hidden from view in Figures 1 and 2.

Other than the 5ZF domain and the SUFU-binding domains described above, the rest of the GLI proteins, comprising 74-80% of the GLI chain, remain unstructured when co-folded with SUFU, and form a disordered cloud around the folded elements; these residues are also hidden from view in Figure 1. Likewise, SUFU residues 1-25 and 279-362 are disordered in all three human structures and are hidden from view in Figures 1 and 2. Evidence suggests that these residues are also intrinsically disordered in native SUFU (*66, 67, 74*), and they have been named intrinsically disordered region 1 (IDR1) and IDR2, respectively (*74*).

Figure 3 shows chord diagrams (a.k.a. Circos plots) of the GLI1-SUFU and GLI2-SUFU interactions (*75*). In this representation, the primary structures of the interacting proteins are displayed radially around the circle, colored by AlphaFold confidence score (pLDDT). The very low confidence regions shown in orange are unstructured and likely intrinsically disordered (*76*). Contacts between the interacting proteins are drawn as arcs connecting them. Chord diagrams for GLI3-SUFU and Ci-dSufu are shown in Figure S3.

**Figure 3.**
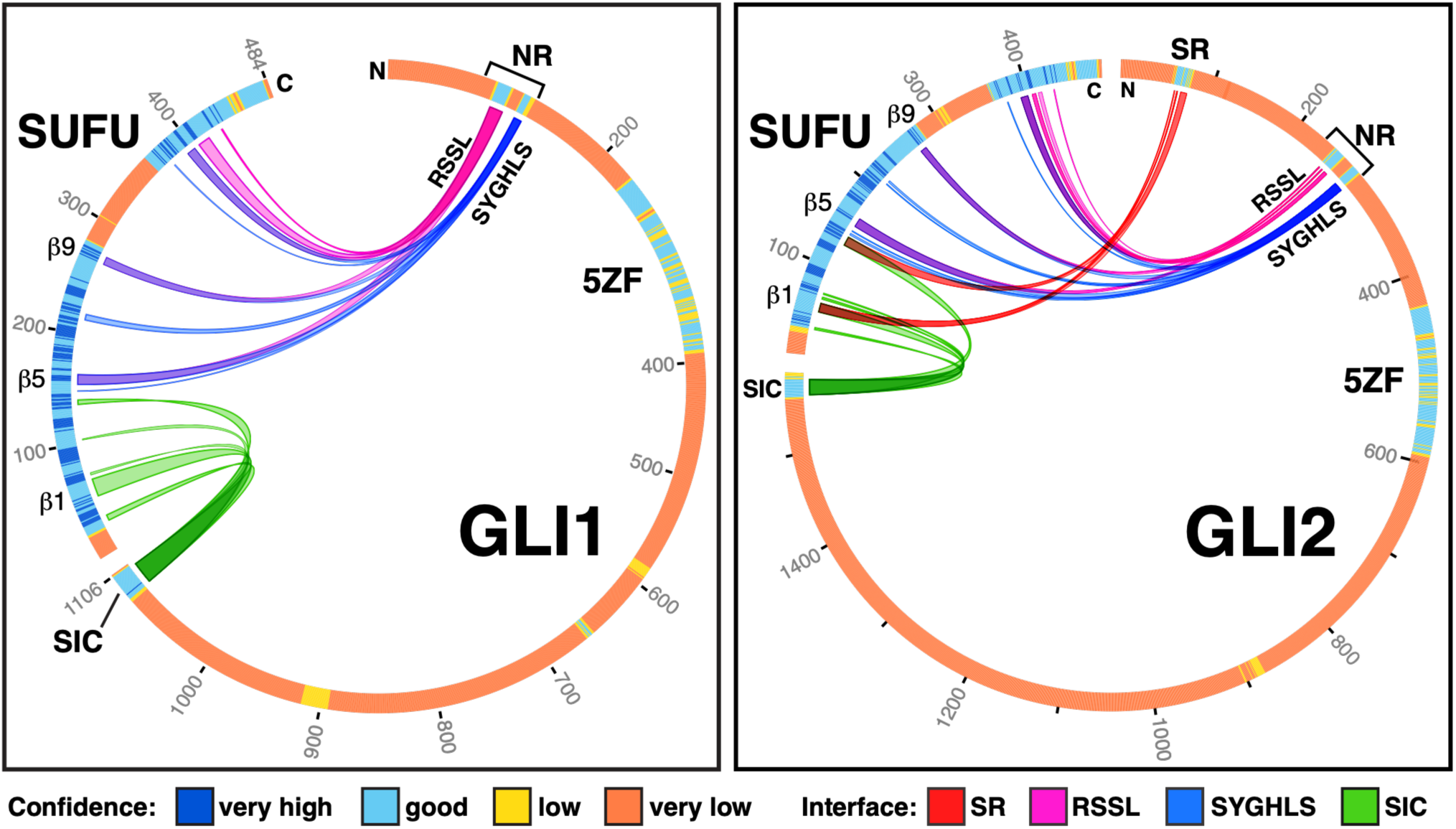
Chord Diagrams of the GLI1-SUFU and GLI2-SUFU interactions. The primary structures of the interacting proteins are displayed radially around the circle, colored by AlphaFold confidence score (pLDDT); note that very low confidence regions (orange) correspond to regions that are likely unstructured/intrinsically-disordered. Contacts between the interacting proteins are drawn as arcs connecting them, colored by interface. The position of the 5-zinc finger DNA-binding domain (’5ZF’) is indicated, as well as the SR motif, the NR domain (which contains the RSSL and SYGHLS motifs), the SIC domain, and SUFU b1, b5 and b9. **Left** GLI1 5z SUFU, CSM 261#. **Right** GLI2 5z SUFU, CSM 91#

### Global quality metrics and negative controls indicate high confidence in the predicted GLI-SUFU complexes

The quality of AlphaFold multiprotein complex predictions can be assessed by three global measures, the ipTM, the pTM and the Model Confidence (MC). All three metrics have a range of 0-1, with scores near 1 indicating higher confidence. The ipTM score measures the accuracy of the predicted interface(s) in a protein-protein complex, pTM measures confidence in the structure of the entire complex, and MC is a linear combination of ipTM and pTM that weights ipTM more. A glossary with more complete descriptions of all scoring metrics is available in the Methods section.

AF3 jobs run with full-length GLI proteins and full-length SUFU had average ipTM scores of 0.78 and MC scores of 0.7, just below the threshold for high confidence. These scores were improved by using SUFU derivatives lacking part or all of IDR2, such as the SUFU-D and SUFUd60 derivatives used in the two published co-crystallization studies. Even higher confidence was obtained by using SUFU-D25, a version of SUFU-D that also lacks the N-terminal 25 amino acids (IDR1). Co-folding of SUFU-D25 with full-length GLI proteins resulted in an average ipTM of 0.9 and average MC of 0.8, values that indicate high confidence (Table S1).

The highest confidence models were obtained by co-folding only the approximately 40 amino acid region of GLI containing both the RSSL and SYGHLS motifs (residues 93-128 in GLI1) with SUFU-D25, or by co-folding this polypeptide and the SIC domain polypeptide with SUFU-D25. This approach did not alter the position of the predicted binding sites, yet resulted in ipTM, pTM and MC scores averaging 0.9 or greater for all three GLI-SUFU complexes (Table S1). These scores indicate very high confidence predictions for both the contact interfaces and the overall structures.

For the *Drosophila* Ci-dSufu complex, co-folding of full-length proteins resulted in confident predictions for the SIC domain interface, but not for the RSSL or SYGHIS motifs. Confident predictions for these interfaces were obtained by co-folding the RSSL+SYGHIS region (residues 226-263) with full-length dSufu, or by co-folding both this polypeptide and the SIC domain (residues 1371-1397) with dSufu. Representative models of GLI-SUFU and Ci-dSufu complexes colored by confidence (pLDDT) are shown in Figure S4.

Other observations support the quality of the AF3-predicted GLI-SUFU complexes. First, in replicate models obtained with different random seeds, the structured portions of the models were highly superimposable with each other, indicating that the same structures and interfaces were repeatedly generated with different random seeds and diffusion sampling (Figures S5, S6). Second, the SUFU and SYGHLS elements of the predicted structures superimposed well with the published co-crystal structures, with average root mean square deviations of less than 1 Å (Table S2, Figure S7). Third, predicted aligned error (PAE) diagrams (Figure S8) also display strong signals for the novel interfaces.

Lastly, we used AF3 to predict the structures of six additional GLI-SUFU complexes from vertebrate model organisms (including GLI1-, GLI2- and GLI3-SUFU from mouse, as well as GLI2-SUFU from chicken, frog and zebrafish) and a further five Gli/Ci-Sufu complexes from model organisms representing diverse invertebrate phyla (Figures S9-S11). These structures show that the GLI-SUFU interfaces described in this work are conserved at the sequence and structural level across metazoans.

As a negative control and to provide a point of comparison, we ran jobs in which glutathione-S-transferase (GST) was used in place of either GLI or SUFU. Biochemical experiments have shown that GST does not bind to GLI or SUFU, even when used at high concentrations (*77, 78*). Consistent with expectations for non-interacting proteins, AF3 co-folding jobs run with GST and GLI or SUFU consistently had ipTM scores of less than 0.5, indicating a failed result for a potential interaction. Furthermore, these models also lacked interfacial contacts between GST and GLI or SUFU that met the confidence metric thresholds described in the next section. Thus, AF3 accurately predicted a lack of interaction between GST and GLI or SUFU (Figure S12).

### Identification of consistent, confident contact residues

To identify high-confidence GLI-SUFU contact residues in AlphaFold3 (AF3) models, we combined AF3 confidence metrics with traditional geometric proximity criteria. We first enumerated interfacial residue pairs with any heavy-atom distance between 1 and 5 Å, then retained only those for which both residues had pLDDT > 70 and a pairwise PAE < 4 Å (a glossary with descriptions of all scoring metrics is available in the Methods section). This definition is intentionally more stringent than the previously described “C+” contacts (pLDDT > 50, PAE < 15 Å, and 1–5 Å separation; (*79*)). Using this pipeline, we compiled contact lists from the top-scoring AF3 models of GLI1/2/3 bound to SUFU, including full-length and IDR-pruned variants, and filtered them to retain only contacts observed in ≥80% of these models across paralogs. All retained contacts were then verified by analysis of the AF3-generated .cif files in ChimeraX, which identifies contact residues based on van der Waals overlap and atomic distance. Contacts which met these criteria are herein designated ‘high-quality’ contacts, as their close spatial positioning is supported by AF3-based confidence metrics, consistent across multiple runs/models/seeds, conserved across paralogs/orthologs, and verified by molecular visualization and analysis software. We recognize that this high threshold may exclude some bona fide interactions, but it enabled us to focus on the most robust, highest-confidence interfaces in this study.

The result of this process was the identification of 9 GLI residues in the RSSL motif, 8 residues in the SYGHLS motif, and 11 residues in the SIC domain that are identical or conserved in all three GLI proteins and contact the same residues in SUFU (Tables S3-S5). The majority of these are conserved in Ci and make similar contacts in the predicted Ci-dSufu structures (Tables S3-S5). In the GLI2/GLI3-specific SR motif, 8 conserved contact residues were identified (Table S6). Figure S13 is an alignment of human SUFU with fly dSufu that shows the predicted GLI/Ci contact residues in both proteins.

### RSSL motif structure and interactions

The first novel structured interface with SUFU predicted by AF3 – the RSSL motif – runs from GLI 95-107, and has four major contact points with SUFU (Figure 4, Figure S14, Table S3). The first half of this sequence forms a short α-helix. Hydrophobic residues (Leu, Met, Ile) on the SUFU-facing side of this helix make hydrophobic contacts with SUFU His394, Thr396, Ala405 and Trp430, providing the first contact point with SUFU. GLI1 Gln96 does not contact SUFU directly; rather, along with Ile99 and Arg100, it sits over the SYGHLS motif and shields GLI1 Ser120, Tyr121 and His123 from solvent.

**Figure 4.**
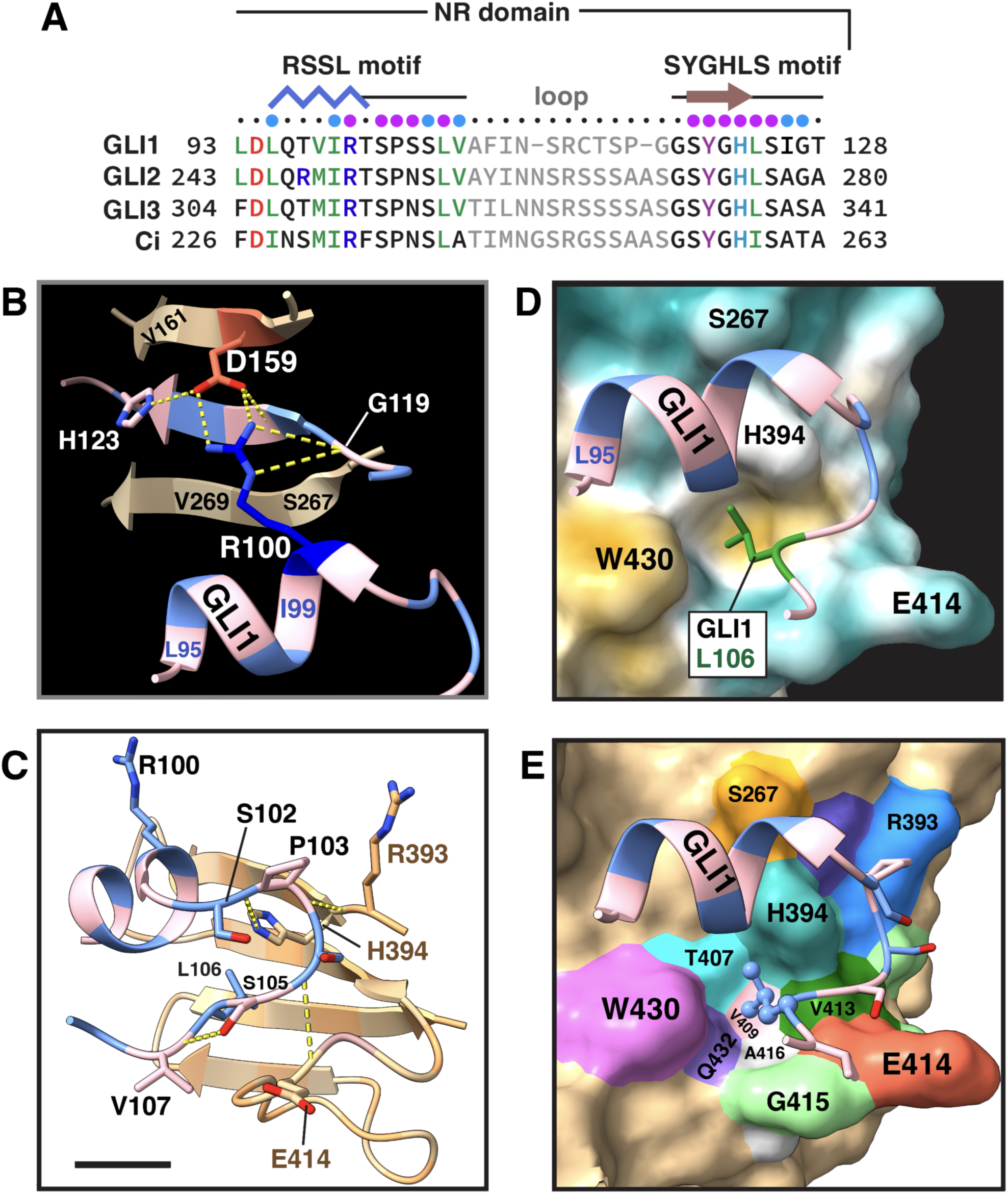
RSSL motif structure and interactions. **A** Alignment of the C-terminal regions of the NR domains of human GLI1, GLI2, GLI3 and *D. melanogaster* Ci. Residue numbers are indicated. Select conserved amino acids are colored as follows: positive/basic residues (K,R,H) blue; negative/acidic (D,E) red, aliphatic hydrophobic (V,I,L,M) green, aromatic (Y,W,F) purple, amides and histidine (N, Q, H) teal. The residues in the disordered loop are colored light gray. The secondary structure for interface regions is shown above, with blue zig-zags for α-helix and brown arrows for β-strands. Cyan and magenta-colored circles denote residues that contact SUFU, with magenta indicating the involvement of at least one H-bond in the contact. The NR domain extends about 15-20 residues N-terminal to the sequence shown here (see Figure S21), but this region is unstructured in the AF3 predictions generated in this work. **B** Hydrogen bonding network involving GLI1 Arg100 (R100) and SUFU Asp159 (D159). GLI1 is colored alternating blue and light pink; SUFU β5 and β9 are colored brown. Select GLI1 and SUFU residues are identified with the single letter amino acid code and their residue number. Model is CSM 70#, GLI1 93-128 SUFU. **C** Hydrogen bonding interactions of the SPSS residues of the GLI1 RSSL motif. GLI1 residues are numbered in black text; SUFU residues are numbered in brown text. SUFU residues 393-435 are shown. **D** Leu106 hydrophobic pocket. The surface of SUFU is colored by hydrophobicity (dark goldenrod = more hydrophobic, dark cyan = less). GLI1 Leu106 (L106) is colored green with side chain shown. **E** SUFU residues that comprise the hydrophobic pocket for Leu106. As in **D**, but slightly different orientation. Leu106 is blue, shown in ball and stick style. SUFU is shown with its surface colored tan and select contact residues in other colors.

The next point of contact between GLI and SUFU on the RSSL motif α-helix is GLI1 Arg100 (Figure 4B). In the predicted structures, this highly-conserved arginine inserts itself between SUFU β5 and the bound SYGHLS motif, forming a classic bidentate salt bridge with SUFU Asp159 in which side chains of Arg100 and Asp159 simultaneously engage in two hydrogen bonds with each other (*80*). GLI1 Arg100 also makes hydrophobic contacts with SUFU Cys156 and GLI Tyr121, and forms two intrachain H-bonds with GLI1 G119.

The short α-helix terminates at GLI1 Thr101, which does not contact SUFU, but does donate a side chain hydrogen to an H-bond with the carbonyl oxygen of GLI1 Thr97. After that, GLI1 residues 102-107 form an extended docking arm that makes two more major contacts with SUFU (Figure 4C, Figure S11). The first of these contact points is the GLI1 Ser102-Pro103 dyad. Both of these residues hydrogen bond to SUFU His394. The carbonyl oxygen of Ser102 accepts an H-bond from the NE2 atom of the His394 imidazole ring, and the carbonyl oxygen of Pro103 accepts an H-bond from the amide nitrogen of His394. Both of these H-bonds are partially shielded from solvent by additional hydrophobic contacts with the aliphatic stem of SUFU Arg393 and the imidazole ring of SUFU His394. GLI1 Ser104 also participates in a shielded main-chain hydrogen bond with SUFU Glu414, which is maintained in GLI2/3 and Ci where Asn is substituted for Ser.

Near the end of the docking arm, the highly conserved GLI1 Leu106 is the fourth major contact point between the RSSL motif and SUFU (Figure 4D, 4E). Leu106 functions as a hydrophobic knob that inserts into a hydrophobic pocket formed by the methyl groups of SUFU Thr407, Val409, Val413 and the face of the His394 imidazole. Weak polar contacts with SUFU His394 and Gln432 also add a stabilizing contribution. Leu106 qualifies as an anchor residue, burying over 70% of its surface area in the SUFU pocket (*81, 82*), and also forms a backbone hydrogen bond with SUFU Glu414.

The side chain hydroxyl of highly-conserved GLI1 Ser105 also donates a hydrogen bond to GLI1 Val107, which may help orient GLI1 Leu106 towards its cognate pocket on SUFU. GLI1 Ser105 and Val107 also make more minor polar and hydrophobic contacts with SUFU.

To reiterate, although we have used human GLI1 and SUFU residue numbers in the above analysis, the contacts described are also all found in the predicted GLI2-SUFU, GLI3-SUFU and Ci-dSufu structures (Table S3, Figures S14, S15).

To summarize, the RSSL motif of GLI1/2/3/Ci adopts a mixed helix-extended structure that makes four main points of contact with SUFU: a hydrophobic patch, a bidentate salt bridge, three shielded H-bonds, and a hydrophobic knob-pocket interaction. Experimental validation of key GLI residues in this motif, and of their contact residues in SUFU, is presented further below.

### SYGHLS motif structure and interactions

Following the RSSL motif are two more elements: an 11-13 residue loop, predicted by AF3 to be unstructured, and the SYGHLS motif. The SYGHLS motif is a well-established, experimentally-demonstrated SUFU-binding motif (*66, 67*). In the AF3-predicted structures and the published co-crystal structures, this motif forms a β-strand, with the SYGH residues participating in a near-canonical parallel β-sheet with SUFU β5 and β9. This binding modality is a type of β-strand addition mechanism known as β-clamping (*83*). Figure 5 shows the predicted hydrogen bonding pattern for GLI2 and Ci, for which there are no existing co-crystal structures. For GLI1 and GLI3, the AF3-predicted SYGHLS-motif structures are almost identical to those observed in the published co-crystal structures (Figure S16). One notable difference is that in the predicted structures, the GLI tyrosine side chain (which is repositioned due to the presence of the salt bridge between GLI1 Arg100 and SUFU Asp159) forms a side-chain to side-chain hydrogen bond with SUFU Ser270. This is also seen in the Ci-dSufu structure between Ci Tyr256 and dSufu Asn266, which replaces Ser270 in dSufu β9. Table S4 lists the high-quality contacts for all residue pairs.

**Figure 5.**
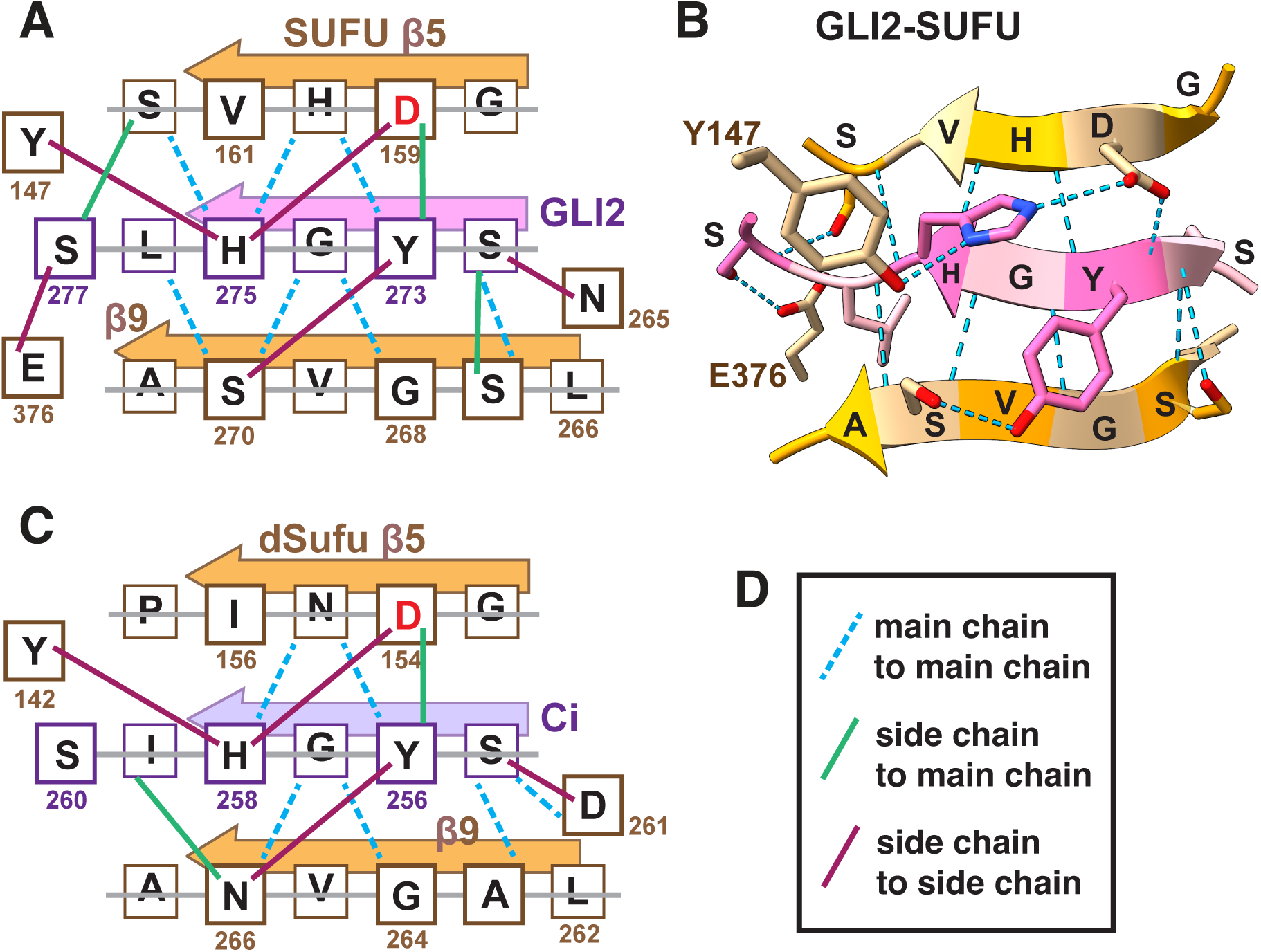
SYGHLS motif interactions. **A** Consensus of AF3-predicted H-bonds in GLI2-SUFU CSMs. The side chains of residues in the larger boxes above the gray line point up relative to the plane of the sheet; those below the line point down. H-bonds are shown as lines or dashes; see **D** for key. **B** Example from GLI2 CSM 255#. All H-bonds shown as blue dashes. **C** Consensus of AF3-predicted H-bonds in Ci-dSufu CSMs. **D** Key for H-bond representation in **A** and **C**.

The RSSL and SYGHLS motifs combine to form a larger interface, herein referred to as the NR domain. (Although the full NR domain, as defined by evolutionary conservation and the presence of other regulatory motifs, extends about 15-20 residues N-terminal to the RSSL motif, as detailed further below.) The RSSL motif and the SYGHLS motif contact each other, and both make extensive contacts with SUFU. Moreover, the α-helix of the RSSL motif sits above the SYGHLS motif and shields the first serine, the histidine and particularly the tyrosine from solvent (Figure S14). As a result, in the predicted GLI-SUFU complexes, the SYGHLS residues become completely buried by the combination of clamping from the side and bottom by SUFU β5 and β9 and capping from the top by the RSSL α-helix. Further, SUFU residues Asp159 in β5 and Ser267 in β9 contact both RSSL (GLI1 Arg100 and Ile99, respectively, Table S3) and SYGHLS (GLI1 Ser120 and Y121/H123 respectively, Table S4).

### SIC domain interactions

The SIC domain resides at the end of the GLI linear sequence and includes the carboxy-terminal residue. In the AF3-predicted structures, the SIC domain is composed of a short β-strand and long α-helix, both of which make contacts with the N-terminal lobe of SUFU (Figure 6). GLI2 residues 1563-1567, Met-Ala-Val-Gly-Asp (and the corresponding residues in GLI1, GLI3 and Ci) are predicted to form an antiparallel β-strand that hydrogen bonds to the exposed edge of SUFU β1 (residues 52-56), adding to a pre-existing 7-stranded β-sheet in the SUFU upper lobe (Figure 6B, 6C). This is a type of β-strand-addition mechanism known as β-strand augmentation (*83*). At the N-end of the strand, GLI2 Asn1562 pins the strand in place via main and side chain hydrogen bonds with SUFU Ile57, Val58 and Asp66, as well as additional polar and nonpolar interactions with these and other (Ala56, Pro65) residues. The side chains of GLI2 Met1563 and Val1565 point down into SUFU, where they contact SUFU Gln53, Val54, Thr55, Thr135 and Gln142 (Table S5).

**Figure 6.**
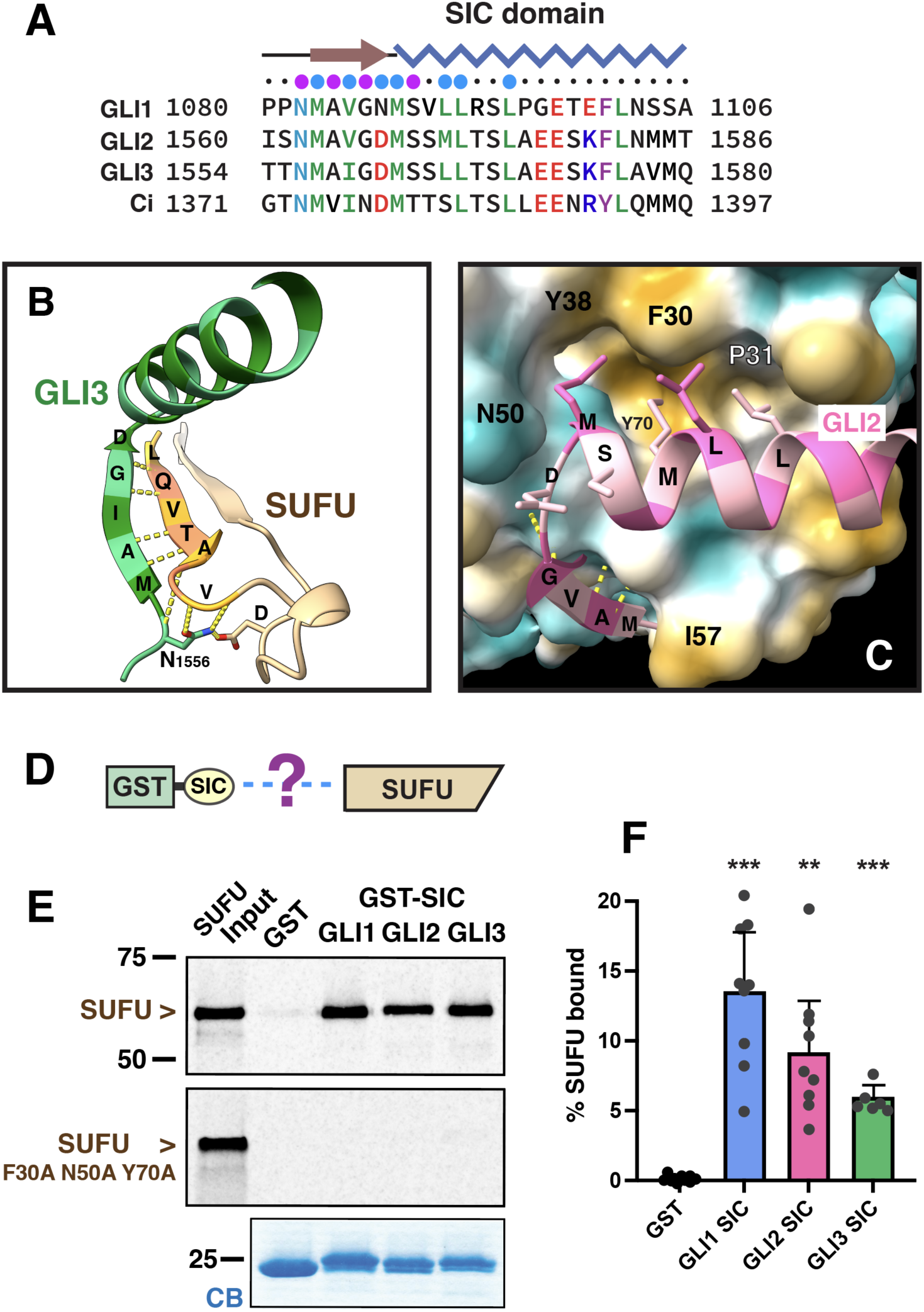
SIC domain structure and interactions. **A** Alignment of the SIC domains of human GLI1, GLI2, GLI3 and *D. melanogaster* Ci. Other details as in Figure 4A. **B** Hydrogen bonds (shown as yellow dashes) between the β-strand of the SIC domain and SUFU β1; model is GLI3 CSM 275#, shown in shades of green. SUFU β1, β2 and connecting residues are shown (tan/yellow/orange). **C** Interaction between the SIC domain amphipathic α-helix and the hydrophobic groove on SUFU. The surface of SUFU is colored by hydrophobicity (dark goldenrod = more hydrophobic, dark cyan = less), with select residues denoted by letters and numbers (e.g. N50). GLI2 is colored alternating shades of pink, with select residues denoted with letters only. Model is CSM 168#. **D** The SIC domains of human GLI1, GLI2 and GLI3, fused to GST, were tested for binding to full-length human SUFU protein. **E** GLI SIC domains are sufficient for binding to SUFU. As shown in **D**, ^35^S-radiolabeled full-length wild-type human SUFU protein (or the F30A N50A Y70A mutant) was prepared by *in vitro* translation and partially purified by ammonium sulfate precipitation, and 10% of the amount added in the binding reactions was resolved on a 12% SDS-polyacrylamide (SDS-PAGE) gel (“SUFU Input”). For binding reactions, 1 pmol of SUFU was incubated with 25 μg of GST or GST-SIC bound to glutathione-Sepharose beads, and the resulting bead-bound protein complexes were isolated by sedimentation and resolved by 12% SDS-PAGE on the same gel. The gel was analyzed by staining with a Coomassie-blue-based reagent for visualization of the bound GST fusion protein (lower panel) and by phosphor imaging for visualization of the bound radiolabeled SUFU (upper panel). The migration of molecular weight markers is indicated on the left. **F** Quantification of 6-10 independent repetitions of the binding assay shown in D and E. 95% confidence intervals are shown; individual data points are displayed as dots. Binding of SUFU to all three SIC domains is significantly different from the minimal binding to GST alone: **, p < 0.01, ***, p < 0.001.

GLI2 Met1568-Thr1586 form a continuous 5-turn α-helix. This helix is amphipathic, with GLI2 Met1568, Met1571, Leu1572, and Leu1575 all lying on the same face. In the predicted complexes, these hydrophobic residues are inserted into a surface-exposed hydrophobic/aromatic groove formed by residues in or near SUFU α2, β1, β2 and α3 (Figure 6C). At the front end of this cleft is a deeper pocket lined by SUFU Tyr38, Asn50, Pro51, and Gln53, into which GLI2 Met1568 buries over 70% of its surface area. Next, GLI2 Met1571 and Leu1572 contact SUFU Phe30, Leu34, Gln53 and Tyr70. Finally, GLI2 Leu1575 interacts mainly with SUFU Pro31, Tyr70 and Tyr99 (Table S5).

GLI1, GLI2, GLI3 and Ci are all predicted to form similar structures and contacts (Figure 6, Figure S15, Table S5). GLI1 has a helix breaking Pro-Gly dyad in the middle of the 5-turn helix just after Leu1095, and in many of the AF3 models we examined, the GLI1 helix was indeed kinked or completely broken at/near this point. However, this kink occurs after the major SUFU contact region.

### Biochemical validation of SIC domain binding to SUFU and of important contact residues

In order to assess the predicted interaction of the GLI SIC domain with SUFU, we fused the C-terminal 27 amino acids of human GLI1, GLI2 and GLI3 to GST, purified the resulting fusion proteins from bacteria, and tested them for binding to full-length human SUFU produced by *in vitro* translation (Figure 6D). As shown in Figure 6E, 6F and S17, SUFU did not bind to GST alone, yet bound with an affinity of ∼50 micromolar to all three GLI SIC domain constructs, validating the predicted interactions. These experiments complement previous results showing that the SIC domains of fly Ci and mouse Gli2 are able to co-sediment fly and mouse Sufu from cell lysates (*72, 84*).

As indicated above, in the AF3-predicted structures, SUFU residues Phe30, Asn50 and Tyr70 make extensive contacts with the hydrophobic knob residues that protrude from the SIC domain α-helix. Consistent with this prediction, a SUFU mutant with these three residues changed to alanine (SUFU F30A, N50A, Y70A) exhibited no detectable binding to the GLI1, GLI2 and GLI3 SIC domains (Figure 6E, Figure S17). Furthermore, substitution of the four knob residues in the GLI1 or GLI2 SIC domains with alanine also abolished SIC-SUFU binding, as did alanine substitution of both residues of the Asn-Met (NM) dyad at the start of the SIC domain β-strand (Figure S17). Collectively, these experiments strongly suggest that the predicted SIC-SUFU interface is correct in many of its details.

### Ci and Fused bind to the same elements of dSufu

The predicted SIC domain-SUFU complexes, particularly in the Ci-dSufu complex, are similar to the co-crystal structure of the SUFU-binding site (SBS) of *D. melanogaster* Fused bound to the N-terminal domain of dSufu (*85*). Indeed, there is previously unrecognized sequence homology between the Fused SBS and the GLI/Ci SIC domain that is concordant with the structural similarity (Figure S18A). In the Fused SBS-dSufu structure, residues 378-382 of Fused form a β-strand that pairs up with the β1 strand of dSufu and is capped by an N-terminal Asn residue, very much as in the predicted GLI/Ci structures (Figure S18B). Moreover, residues 383-387 of Fused form a 3_10_-helix-like structure, with residues Met383 and Phe387 protruding into the dSufu hydrophobic cleft and making contacts similar to those made by Ci Met1379 and Leu1383 (Figure S18C).

### SR (SUFU-binding, repressor domain) motif

The SR motif (GLI2 residues 61-75 and GLI3 residues 134-148) is predicted to interact both with the folded SIC domain and with residues in the N-terminal lobe of SUFU (Figures S19, S20, Table S6). The sequence of the SR motif is positioned in an N-terminal region of GLI2/3 that has been broadly defined as a repressor domain (*86–88*). Both the sequence and predicted structure of the SR motif are conserved between GLI2/GLI3 and multiple vertebrate and invertebrate GLI orthologs (Figures S9-S11), but it is not found in GLI1 or Ci (Figures 1–3, Figure S3). The SR motif is unfolded and unstructured in GLI-SUFU CSMs lacking the SIC domain, including the GLI3 repressor fragment (Figure S20).

His141 in the GLI3 SR motif has been shown to be important for GLI3’s transcriptional repression activity (*89*). To our knowledge, however, there is no experimental evidence that this region of GLI2 or GLI3 binds directly to SUFU or to any other proteins, although it should be noted that the N-terminal SR domain might not be expected to bind to SUFU in the absence of the C-terminal SIC domain. We did not attempt to validate the SR motif-SUFU interaction in this study.

### Validation of the critical role of GLI1 R100 and L106 in SUFU binding

Two residues in the NR domain of GLI1 that make critical contacts in the new predicted structure are Arg100 (R100) and Leu106 (L106; we switch to the single letter amino acid abbreviations for reporting experimental observations). As shown in Figure 4B, in the AF3 structure, R100 sits above the SYGHLS strand, forming hydrogen bonds with both Gly119 (G119) in GLI1 and Asp159 (D159) in SUFU, the latter a classic Arg-Asp bidentate salt bridge. Indeed, GLI1 R100 and SUFU D159 combine to form multiple sidechain-sidechain and sidechain-mainchain H-bonds with the SYGHLS strand. The homologous GLI2/GLI3/Ci residues R250/R311/R233 are similarly positioned and make equivalent contacts (Figure S15, Table S3). L106 (and the homologous residues L256/L317/L239) qualifies as a key anchor residue that buries 70% of its surface area in a hydrophobic pocket formed by multiple residues in SUFU.

To validate the importance of these residues for the GLI1-SUFU interaction, we individually mutated them to alanine and to glutamate and tested their binding to SUFU. These substitutions were made in a plasmid encoding GLI1 residues 1-232 downstream of a T7 RNA polymerase promoter, because we previously found that this is a sensitive system for analyzing the effect of GLI1 substitution mutations on SUFU binding (*77*).

Next, full-length human SUFU protein was produced in, and purified from, *E. coli* bacteria as an N-terminal GST fusion protein. The purified GST-SUFU was then tested for binding to wild-type GLI1_1-232_ that had been produced and radiolabeled by *in vitro* transcription/translation, as well as to GLI1_1-232_ substituted in R100 or L106 (Fig 7A). Each residue was substituted either with Alanine (Ala, A) or Glutamate (Glu, E), resulting in a total of four variants.

**Figure 7.**
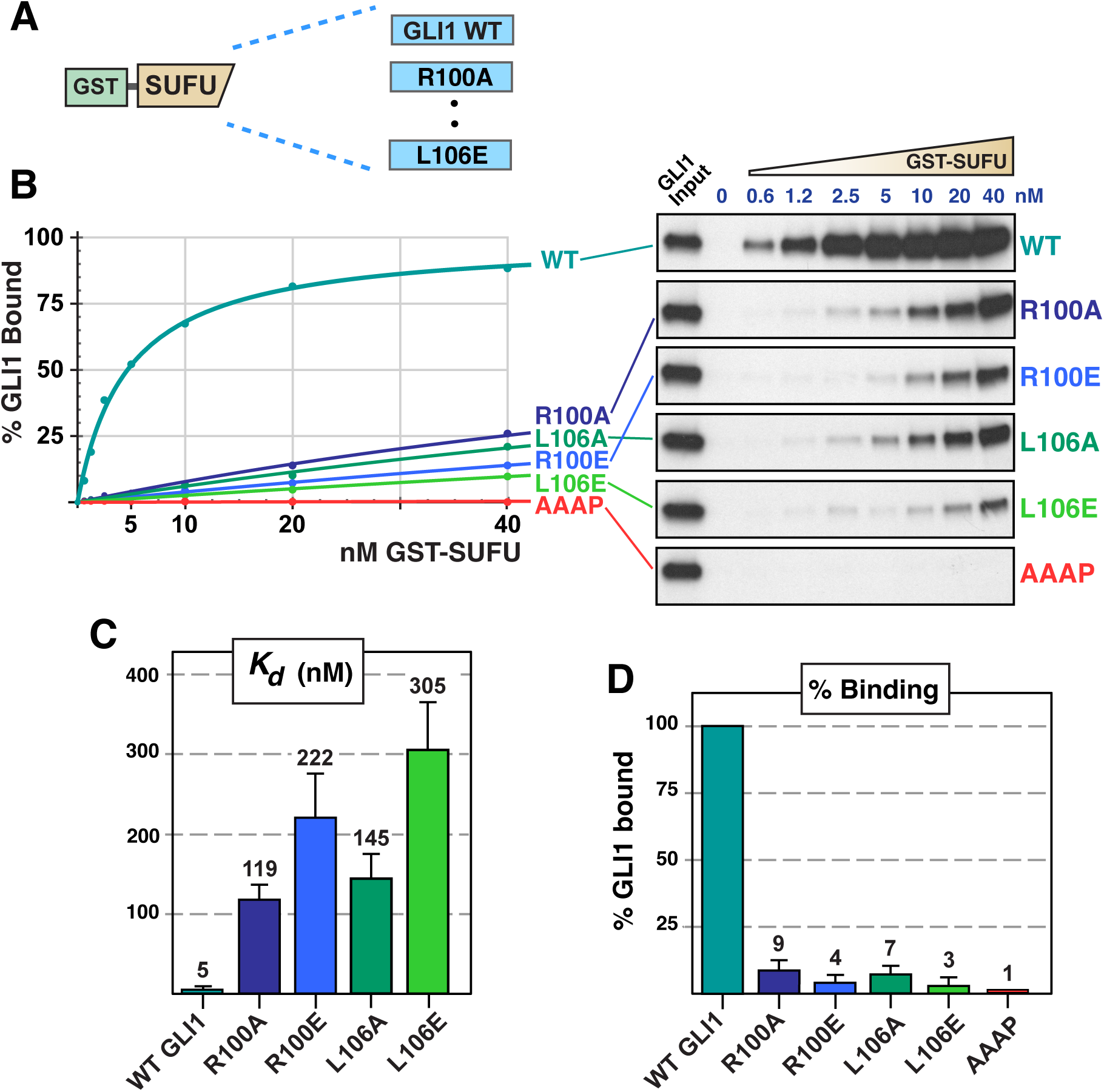
GLI1 R100 and L106 mutations decrease binding to SUFU. **A** Radiolabeled GLI1_1-232_ variants were assessed for binding to GST-SUFU protein. **B** GLI1-SUFU binding isotherms (left) and gel analysis of a representative experiment (right). The data points on the graph are an average of 3 independent experiments. The input lanes were loaded with 10% of the amount added to binding reactions. The ‘AAAP’ variant is a YGHL to AAAP change in the SYGHLS motif. **C** Quantification of dissociation constants (Kd) for the five GLI1_1-232_ alleles. Higher Kds equate to lower levels of binding. Each data point in 3 independent experiments was converted to a Kd as described in Materials and Methods, n = 21. Numbers above each bar are the mean Kd in nanomolar units; error bars show the 95% confidence interval. The difference between each of the four mutants and the wild-type is statistically significant with very high confidence, p < 0.0001. **D** Data from the 2.5, 5 and 10 nM GST-SUFU points from all replicate experiments were averaged to determine percent binding and normalized to wild-type, n = 9. Numbers above each bar are the mean percent binding; error bars show the 95% confidence interval. The difference between each of the five mutants and the wild-type is very highly significant.

As shown in Fig 7, GST-SUFU bound tightly to radiolabeled wild-type GLI1_1-232_, with a calculated dissociation constant (Kd) of 5 nM. As noted earlier, the GLI proteins do not detectably bind to GST alone, even at GST concentrations exceeding 1500 nM (*77, 78*). Both substitutions in R100 and both substitutions in L106 had a dramatic effect on GLI1-SUFU binding, increasing the dissociation constant (and correspondingly decreasing the binding affinity) by 24-61 fold.

Increases in the dissociation constant correspond to proportional decreases in affinity, as Kd is the reciprocal of the affinity constant. Similarly, increases in Kd correspond to decreases in the amount of protein complex formed. For example, a 20-fold increase in Kd can result in up to a 95% decrease in the amount of heterodimer formed, and a 60-fold increase in Kd can result in up to a 98% decrease, depending on the concentrations of the interacting proteins relative to the Kd of their interaction (e.g. Figure 7D).

These results confirm that R100 and L106 are critical determinants of GLI-SUFU binding affinity, consistent with the structures predicted by AlphaFold3.

### Scanning Mutagenesis of GLI1 reveals NR domain contact residues

As previously noted, the RSSL motif and the SYGHLS motif are part of a larger 55-60 amino acid domain called the NR (N-terminal Regulatory) domain or the SIN (SUFU-Interacting, N-terminal) domain (*71, 72*). Other than the 5ZF domain, the NR domain is the most conserved region between vertebrate and invertebrate GLI proteins, and was shown to be sufficient to bind to SUFU in early studies (*71, 90*). As defined by sequence conservation, the NR domain consists of GLI1 residues 78-131 and the corresponding residues in GLI2/3 and Ci (Figure S21, Figure 8A; as before, residue numbering from human GLI1 and human SUFU will usually be used). The structure of the N-terminal portion of the NR domain (GLI 78-94) is not confidently predicted by AF3 and does not contact SUFU based on our strict definition of contact residues. This region has had multiple functions ascribed to it, including functioning as a ciliary-localization signal (*91*), a docking site for MAP kinases (*77, 78*), and a target site for phosphorylation by multiple protein kinases (*92, 93*).

**Figure 8.**
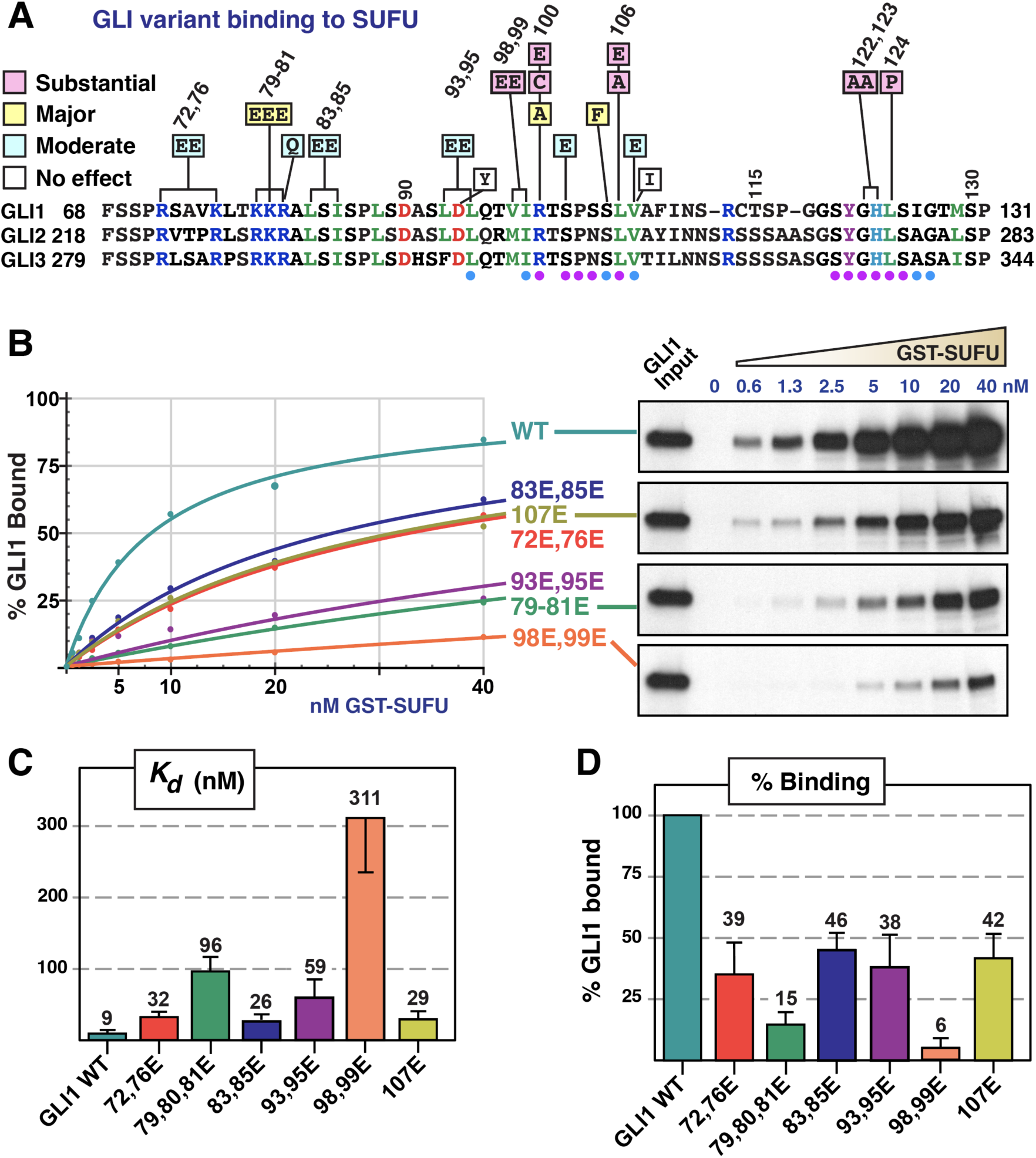
Effect of additional GLI1 mutations on binding to SUFU. **A** Chart summarizing the effect of GLI1 NR domain mutations on binding to SUFU from this study and from (*77*). The chart shows which substitutions in GLI1_1-232_ had no effect (defined as an affinity change that was either less than 2-fold, and/or was not statistically significant), a minor effect (affinity decrease between 3 and 10-fold that was statistically significant), a major effect (greater than 10-fold, significant), or a substantial effect (greater than 30-fold, significant). Fourteen additional “no effect” substitutions in non-contact residues can be found in (*77*). An alignment of human GLI1 residues 68-131 with the corresponding regions of human GLI2 and GLI3 is also shown. Amino acid coloring as in Figure 4A; circles below indicate contacts as in Figure 4A. **B** Radiolabeled GLI1_1-232_ variants were assessed for binding to GST-SUFU protein. GLI1-SUFU binding isotherms (left) and gel analysis of a representative experiment (right). The input lanes were loaded with 10% of the amount added to binding reactions. The data points on the graph are an average of 2 independent experiments. **C** Quantification of dissociation constants (Kd) for the GLI1_1-232_ alleles. Each data point in the 2 experiments was converted to a Kd as described in Materials and Methods, n = 14. Numbers above each bar are the mean Kd in nanomolar units; error bars show the 95% confidence interval. The difference between each of the six mutants and the wild-type is statistically significant, p < 0.0025 or better. **D** Data from the 2.5, 5 and 10 nM GST-SUFU points from all replicate experiments were averaged to determine percent binding and normalized to wild-type, n = 6. Numbers above each bar are the mean percent binding; error bars show the 95% confidence interval. The difference between each of the six mutants and the wild-type is very highly significant.

In a previous study, we engineered over 30 different mutations in the NR domain of GLI1, both to investigate the role of MAP kinase phosphorylation of GLI1 serines S102, S116 and S130 on SUFU release, and to probe the effect of tumor-derived mutations in this region on SUFU binding (*77*). Of these, the most relevant to the present work are GLI1 R100C, S102A, S102E and S105F, all in the RSSL motif. The GLI1 R100C mutation was found in four different somatic tumor samples, and resulted in a 35-fold decrease in GLI1-SUFU binding affinity in the same assay as used in this study. Mutations in GLI1 S105 were also found in multiple somatic tumor samples, with the S105F substitution resulting in a 13-fold decrease in GLI1-SUFU binding affinity. Finally, GLI S102A and S102E – substitutions in a MAPK phosphorylation site – resulted in 3.5- and 8-fold affinity reductions, respectively. These prior results support the importance of key RSSL motif residues in SUFU binding.

Another tumor-derived mutation we previously examined is L124P in the GLI1 SYGHLS motif. This mutation, from a drug-resistant basal cell carcinoma patient, diminished SUFU binding by ∼250-fold (*77*).

To assess the involvement of additional GLI residues in the NR domain in SUFU binding, scanning mutagenesis of GLI1 was performed. In order to maximize our ability to detect important residues for SUFU binding, two or three closely-spaced residues were simultaneously substituted, making radical substitutions that switched charge (e.g. positively-charged arginine or lysine to negatively-charged glutamate) or replaced a hydrophobic residue with glutamate. These experiments were performed relatively early in this study before we had developed our definition of high-quality contact residues; thus, some non-contact residues were included in the analysis.

The first set of substitutions was in the N-terminal portion of the NR domain. As shown in Figure 8, the double substitution GLI1 R72E/R76E in this region had a relatively minor effect on SUFU binding, corresponding to a 4-fold reduction in affinity. Likewise, the double substitution GLI1 L83E/I85E resulted in a minor 3-fold reduction in SUFU binding. In contrast, the triple substitution GLI1 K79E/K80E/R81E reduced SUFU binding affinity by 11-fold. These three Lys/Arg residues – which constitute the basic submotif of the D-site for MAPK docking – do not contact SUFU directly in the majority of AF3 models we examined. The predicted models do show, however, that these positively charged residues are spatially proximal to a large negatively charged patch on SUFU (Figure S22). Thus, they may interact with SUFU using the same “fuzzy” electrostatic mechanism that has been proposed for the basic submotif of MAPK D-sites (*94*).

Radical substitutions in the RSSL motif included L93E/L95E, V98E/I99E and V107E; of these, L95, I99 and V107 make high-quality contacts with SUFU. These mutations resulted in 7-, 35-, and 3-fold decreases in binding affinity, respectively (Figure 8).

Figure 8A summarizes the results of all the GLI1 substitution mutants examined in this study, as well as selected tumor-derived substitutions analyzed in our 2022 study (*77*). The graphic highlights that mutations in the RSSL motif can dramatically impact GLI1-SUFU binding.

### SUFU mutations in contact residues

To further validate the AF3-predicted models and the predicted contacts between GLI and SUFU, we constructed a set of 18 single amino acid substitutions in SUFU residues predicted to make high-quality contacts with the GLI RSSL motif, and a further 6 mutations in non-contact residues that we nevertheless thought it would be interesting to test.

The first SUFU variants we constructed were radical substitutions of predicted contact residues, including SUFU D159R, A405E, T407E, V409E, E414G/R, and W430E. A second group of mutations that we tested was chosen from those deposited in the ClinVar database (*95*). These mutations – sequenced from patients or their families afflicted with, or thought to be at risk for, medulloblastoma or Gorlin syndrome – were anonymously deposited in ClinVar by clinical genetics laboratories such as Ambry Genetics, Labcorp, etc., or by academic medical center labs such as those at Baylor and Mass General Brigham (*96*). These 13 “patient mutations” were C156R, G392R, R393W, H394D, M404K, I406T, T407M, G412E, V413G, G415D, A416D, W430S, and Q432P. To this we added two variants found in sporadic tumors, A405T and W430C, to make a set of 15 patient/tumor mutations. Of these 15, 11 are in contact residues, while four – G392R, M404K, I406T, and G412E– are adjacent to contact residues.

These mutations were introduced into full-length human SUFU, and the SUFU variants were then produced by *in vitro* transcription/translation and tested for binding to GLI1 in a quantitative assay (Figure 9A). GST-GLI1_68-232_ was chosen as the binding partner for full-length SUFU in these assays for the following reasons: (1) it is readily expressed in soluble form and purified from bacteria; (2) it begins just before the start of the NR domain and ends just before the start of the 5ZF domain, and thus contains both the RSSL and SYGHLS motifs, but no other SUFU-interacting regions; (3) it was used extensively and thoroughly characterized in both our 2010 and 2022 studies (*77, 78*).

**Figure 9.**
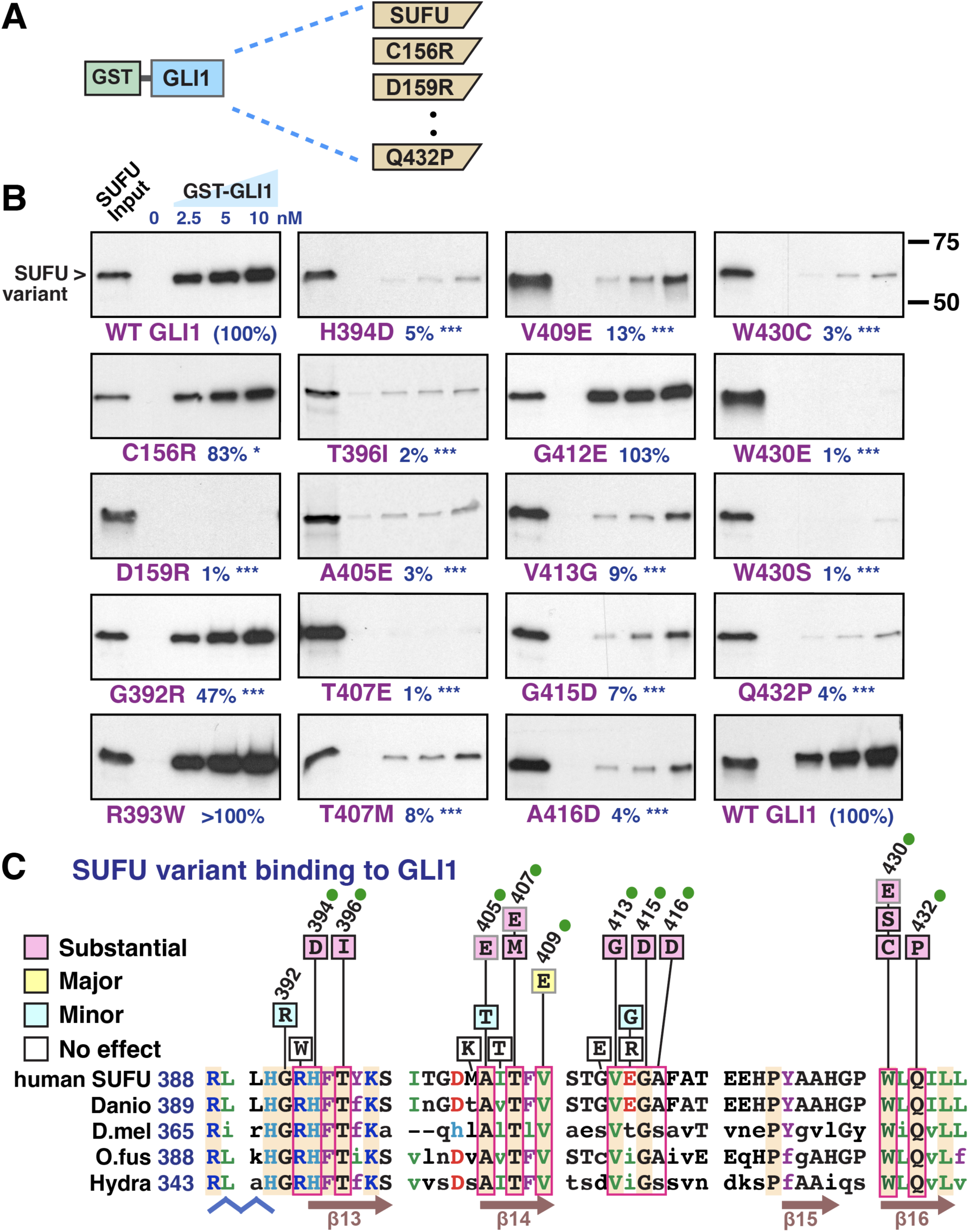
SUFU mutations in GLI contact residues decrease GLI1-SUFU binding. **A** Radiolabeled SUFU variants were assessed for binding to GST-GLI1_68-232_ protein. **B** Representative gel analysis results of the experiment described in **A**. The input lanes were loaded with 10% of the amount added to binding reactions. The mean percent binding (average of all three concentrations) from 2-4 independent experiments is shown below (n = 6-12). The statistical significance of the difference from wild-type is also shown: *, significant; **, highly significant; ***, very highly significant. The position of molecular weight markers is shown to the right of the W430C image. **C** Chart summarizing the effect of 21 SUFU mutations on binding to GLI1. The chart shows which single substitutions in human SUFU had no effect (binding not decreased compared to wild-type), a minor effect (binding between 50% and 20% of wild-type), a major effect (binding between 20% and 10% of wild-type), or a substantial effect (binding less than 10% of wild-type). The variants are patient/tumor mutations except for A405E, T407E, V409E and W430E. Variants confirmed defective for binding GLI2 and GLI3 (see Figures 10 and S23) are indicated with a green circle. An alignment of human SUFU residues 388-435 with the corresponding regions of SUFU from one vertebrate (zebrafish *Danio rerio,* NP_958466) and three invertebrate model organisms (fruit fly *Drosophila melanogaster*, NP_536750; annelid worm *Owenia fusiformis*, CAH1784014; cnidarian polyp *Hydra vulgaris,* XP_012560454) is shown below. Residues in upper case are identical to human SUFU; residues identical in all five species are highlighted in orange; high-quality contact residues are boxed with pink outlines. Amino acid coloring as in Figure 4A. Secondary structure is shown below.

The results of these experiments are shown in Figure 9 and Figure S23. Because we wanted to be able to test more variants, we did not attempt to fit the data to isotherms as we did in Figures 7 and 8, but instead tested each SUFU mutant against 3 different concentrations of GST-GLI1 in the pseudo-linear range of the isotherm. Thus, results are reported as percent of wild-type binding efficiency only, and not as fold decreases in affinity.

SUFU D159R served as a positive control, as it has been previously tested by Zhang *et al*. and shown to be defective in GLI1 and GLI2 binding (*67*). In both the empirical and the AF3-predicted structures, D159 hydrogen bonds to GLI1 Y121 and H123 in the SYGHLS motif. In the AF3-predicted structures SUFU D159 also makes critical contacts to GLI1 R100; these same interactions occur between dSufu D154 and Ci R233, Y256 and H258 in the fly structure. As shown in Figure 9B, the SUFU D159R mutation substantially decreased binding to GST-GLI1 to 1% of wild type.

SUFU H394, located at the start of β13, contacts several residues in the GLI RSSL motif. It hydrogen bonds with GLI1 S102 and P103, and makes hydrophobic contacts with GLI1 I99 and L106. Equivalent interactions are seen between dSufu H371 and Ci S235, P236, I232, and L239. H394D is a patient mutation found in three different reports submitted to ClinVar; as shown in Figure 9B, this mutation substantially decreased SUFU binding to GST-GLI1 to 5% of wild type. SUFU T396, on the same face of β13 as H394, also contacts GLI1 I99. A T396I mutation substantially decreased SUFU binding to GST-GLI1 to 2% of wild type (Figure 9B); this substitution is equivalent to a mouse Sufu mutation associated with limb defects, as is expanded upon further below.

SUFU W430, located at the start of β16, makes multiple atomic contacts with GLI1 L95. In GLI2, GLI3 and Ci it may also contact a methionine residue three amino acids further on, although this was not supported by all our confidence metrics. We tested three different SUFU variants with W430 mutations: W430E, a radical substitution; W430C, from a sporadic tumor; and W430S, from a patient sequence deposited in ClinVar. All three mutations substantially decreased SUFU binding to GST-GLI1 to 3% or less of wild type.

GLI1 L106 buries over 70% of its surface area in a hydrophobic pocket formed by multiple SUFU residues, including H394, T407, V409, V413, A416, and Q432 (Figure 4D, 4E). Pocket residue substitutions H394D, T407E, T407M, V409E, V413G, A416D and Q432P each had a dramatic effect on complex formation, decreasing binding to between 1%-8% of wild type.

Overall, we constructed and tested SUFU mutations in 14 different predicted GLI-contact residues. For 11 of those contact residues, one or more of the variants we tested exhibited a major/substantial defect in GLI1 binding (Figure 9, Figure S23). In contrast, 5 of the 6 non-contact residues tested did not affect GLI1 binding, while one (G392R) had a minor defect. For the 15 patient/tumor mutations tested, 8 displayed a substantial defect in GLI1 binding (all 8 of these were also contact residues), two (G392R and I405T) displayed a modest reduction, and 5 (C156R, R393W, M404K, I406T and G412E) had no effect.

These results demonstrate that the vast majority of AF3-predicted contact residues in SUFU are in fact important for GLI1 binding.

### SUFU mutations - effect on GLI2 and GLI3 binding

As stated previously, in the AF3-predicted structures, the SUFU residues that contact GLI1 make equivalent contacts with GLI2 and GLI3. For example, R250/R311 in GLI2/GLI3 form a salt bridge with SUFU D159, while S252/S313 in GLI2/GLI3 form a hydrogen bond with SUFU H394, and L256/L317 in GLI2/GLI3 insert into the same hydrophobic pocket as GLI1 L106. Hence, we strongly predicted that SUFU variants defective in binding to GLI1 would also be defective in binding to GLI2 and GLI3.

To verify this prediction, we first expressed and purified GST-GLI2_218-433_ and GST-GLI3_279-480_ from bacteria. These constructs are similar to GST-GLI1_68-232_ in that they begin just before the start of the RSSL motif and end just before the start of the 5ZF domain, and they are soluble and readily purified. Next, we verified that purified GST-GLI2 and GST-GLI3 bound to wild-type full-length human SUFU with low nanomolar affinity (Fig 10). Finally, we tested their binding to 11 SUFU variants with mutations in SUFU-GLI contact residues – including 8 patient/tumor mutations – all of which were found to be substantially defective in GLI1 binding. The results are shown in Figure 10 and Figure S23, which also contain additional data points with GST-GLI1 for comparison. As can be seen, all 11 variants tested were indeed defective in GLI2 and GLI3 binding, as expected based on the results obtained with GLI1.

**Figure 10.**
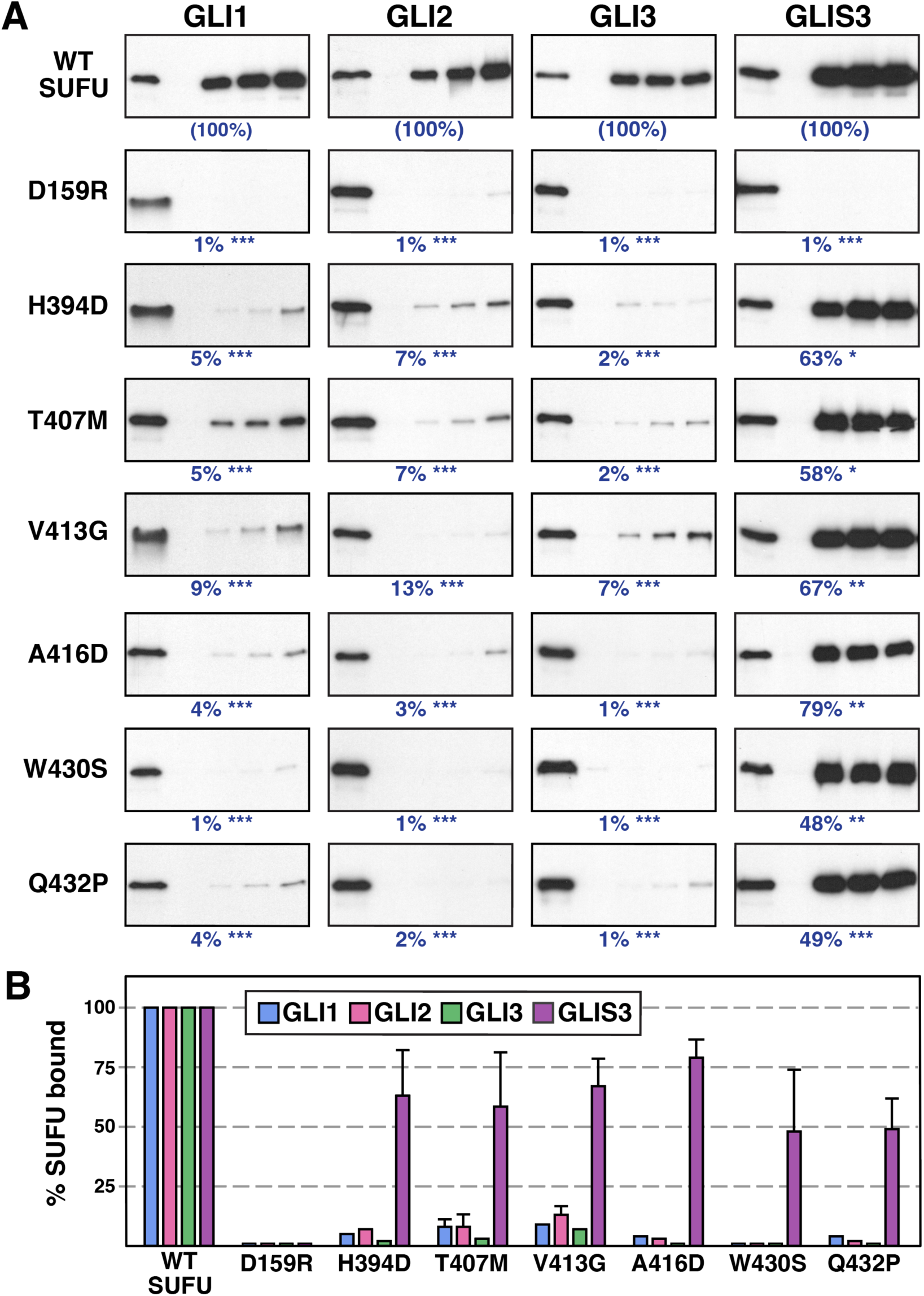
SUFU mutations in GLI contact residues compromise binding to all 3 GLI proteins but not to GLIS3. **A** Radiolabeled SUFU variants were assessed for binding to GST-GLI1_68-232_, GST-GLI2_218-433_, GST-GLI3_279-480_, or GST-GLIS3_105-284_ proteins. Representative gel analysis results of 2-4 independent experiments are shown. Protein concentrations and other details are as in Figure 9. **B** Data from the 2.5, 5 and 10 nM GST-GLI/GLIS points from all replicate experiments were averaged to determine percent binding and normalized to wild-type, n = 6-12. Error bars show the 95% confidence interval.

These results provide strong experimental support for the majority of AlphaFold-predicted contacts between the GLI1, GLI2 and GLI3 RSSL motifs and SUFU, and confirm the dramatic effect of patient mutations in predicted contact residues on GLI-SUFU binding.

### GLIS3 binds differentially to SUFU

Human GLIS3 (GLI similar 3) is a known SUFU-binding protein that has sequence similarity with GLI1/2/3 in the 5-zinc finger DNA-binding domain and in the NR domain. This similarity extends to key residues in the RSSL motif (Figure S24). Furthermore, GLIS3 has a YGH sequence that aligns with the GLI SYGHLS motif, and mutation of this sequence to AAA reduced GLIS3-SUFU binding (*22*). GLIS3 does not play a role in Hedgehog signaling, but functions as a transcription factor required for proper development of the kidney, thyroid and β-cells of the pancreas. Homozygous loss-of-function mutations in *GLIS3* cause autosomal recessive neonatal diabetes mellitus with congenital hypothyroidism syndrome (NDH syndrome) (*25*).

Based on the NR domain similarity, we set out to ask if GLIS3-SUFU binding was affected by the same SUFU point mutations that disrupted GLI-SUFU binding. To do so, we first expressed and purified GST-GLIS3_105-284_ from bacteria. As with the GST-GLI proteins used in this study, GST-GLIS3_105-284_ begins just before the start of the NR domain and ends just before the start of the 5ZF domain. It should be noted, however, that there is no notable sequence similarity between GLIS3 and the GLI proteins in the region between the end of the NR domain and the 5ZF domain.

GST-GLIS3_105-284_ binds to wild-type SUFU with low nanomolar affinity (Figure 10). Moreover, this interaction is substantially disrupted by the SUFU D159R mutation: SUFU D159R bound to GST-GLIS3 with only 1% the efficiency of wild type. Surprisingly, however, the remaining SUFU variants tested showed a much more moderate effect on GLIS3 binding, as they were able to bind with 45-79% wild-type efficiency (Figure 10, Figure S23).

To investigate potential reasons for this interesting difference, we used the AlphaFold server to co-fold human GLIS3_105-284_ with full-length human SUFU. The results confirm that GLIS3 RSSL- and YGH-mediated contacts with SUFU are very similar to those seen with the GLI proteins, but also delineate two additional contact regions that may explain the increased relative affinity of GLIS3 for SUFU mutants that otherwise disrupt GLI binding (Figures S24, S25).

### Some contact residue mutations have been assessed in cells and animals

Because our goal was to systematically test AF3-predicted interfaces, we focused our validation experiments on quantitative *in vitro* binding assays. Nevertheless, several SUFU and GLI residues that we identify as key components of the NR-domain interface have previously been analyzed in cell- and animal-based studies, and the reported phenotypes are consistent with impaired SUFU-mediated repression of GLI activity. Human and mouse SUFU proteins are both 484 amino acids long, 98% identical, and have concordant residue numbering. Merchant *et al*. showed that substituting mouse Sufu residues 391-394 with four alanines, or 395-398 with four alanines, severely disrupted the Gli1-Sufu interaction and Sufu-dependent repression of Gli1-driven transcription (*97*). Both of these 4-residue blocks include residues that our models place at the RSSL contact surface (Figure 9, Table S3).

Several single-residue variants further support the functional importance of this interface. SUFU Thr396 contacts the hydrophobic patch on the RSSL motif α-helix (Table S3); Makino *et al*. isolated a Thr396Ile allele of mouse Sufu in an ENU mutagenesis screen and showed that it caused polydactyly when homozygous, with concomitant destabilization of Gli3 (*98*). We found that the orthologous substitution in human SUFU, T396I, was substantially defective in binding to GLI1, GLI2, and GLI3 (Figure 9, Figure S23). SUFU Thr407 contributes to the hydrophobic pocket that accommodates GLI1 Leu106 (Figure 4D,4E, Table S3); Takenaka *et al*. generated mouse Sufu Thr407Glu as a phosphomimetic of a proposed GSK3β phosphorylation site, and showed that it reduced Gli1-Sufu co-IP, Sufu-dependent repression of Gli1-driven transcription, and Sufu-dependent inhibition of HH-ligand stimulated differentiation (*99*). Consistent with those observations, we found that human SUFU T407E was substantially defective for GLI1 binding, and the T407M patient variant was defective for binding to GLI1, GLI2, and GLI3 (Figures 9 and 10). Additionally, multiple studies have verified the importance of SYGHLS-motif contact residues for SUFU function in cells (*66–68, 77*). This includes SUFU Asp159, which contacts both the RSSL and SYGHLS motifs.

In summary, multiple different SUFU residues that were shown in this study to make important contacts with the GLI NR domain perturb GLI-SUFU binding and HH signaling *in vivo* when mutated.

### Phosphorylation of GLI-SUFU contact regions

Several residues within, or adjacent to, the GLI-SUFU contact surfaces defined here are known phosphorylation sites with functional consequences consistent with altered SUFU binding and regulation. In particular, GLI1 Ser102 (and the corresponding residues in GLI2 and GLI3, Ser252 and Ser313) lies at the core of the RSSL motif, where it is positioned to hydrogen-bond with SUFU His394 (Figure 4C, Table S3). GLI1 Ser102 has been reported to be phosphorylated by MAP kinases and by AMP-activated protein kinase, and GLI1 phosphomimetic variants (in combination with other substitutions) are hyperactive in cell-based assays (*52, 77*). In mouse cells, phosphorylation of Gli2 Ser248 (equivalent to human GLI2 Ser252) increases in response to Hedgehog stimulation (*100*). Furthermore, DYRK2 has been reported to phosphorylate GLI2 Ser252 and GLI3 Ser313, promoting their dissociation from SUFU (*101*).

Additional phosphorylation events in the NR region have been linked to SUFU regulation. Several studies report functional phosphorylation of GLI2 Ser230 and Ser232 (or equivalent residues in GLI1 and Ci) (*84, 92, 93*); these residues lie near basic, positively-charged residues that we propose may contribute to SUFU association via electrostatic (“fuzzy”) interactions (Figures 8 and S22), raising the possibility that nearby phosphorylation could weaken binding by introducing local negative charge.

Finally, GLI2 Ser1570, Thr1573 and Ser1574 lie on the solvent-exposed face of the amphipathic SIC domain helix, and do not directly contact SUFU (Figure 6). Nevertheless, phosphomimetic substitutions in the equivalent residues in mouse Gli2 and fly Ci have been shown to cooperate with NR-domain phosphomimetics to weaken Sufu binding and repression (*84*). Together, these observations support the idea that the NR and SIC domains are recurrent targets of regulatory phosphorylation positioned to modulate the GLI-SUFU interaction.

## DISCUSSION

A central problem in Hedgehog signal transduction is how the SUFU tumor suppressor protein regulates the activity of GLI transcription factors and how this is modulated during pathway activation. A closely related question is how *SUFU* mutations –particularly missense alleles– may compromise SUFU function in patients with Shh-type medulloblastoma, Gorlin syndrome, Joubert syndrome, neonatal diabetes, and other diseases. In this study, we used AlphaFold3 (AF3) to predict high-confidence structures of the human GLI1-SUFU, GLI2-SUFU, and GLI3-SUFU complexes, the structure of the fruit fly (*D. melanogaster*) Ci-dSufu complex, and the structures of eleven other orthologous vertebrate and invertebrate GLI-SUFU complexes (Figures 1–3, S9-S11). These structures revealed three new domains/motifs in GLI and their structural interfaces with SUFU. The first of these, the NR domain, contains both the previously-characterized SUFU-binding peptide (designated the SYGHLS motif) and the novel RSSL motif, which together form an integrated SUFU-binding domain that contacts SUFU β5, β9 and multiple additional residues in the C-terminal lobe of SUFU (Figures 4, 5). The second new structural domain, the C-terminal SIC domain, contacts SUFU β1 and additional residues in the N-terminal lobe of SUFU (Figure 6). The third, the SR motif, is present in the repressor domains of GLI2 and GLI3. In the predicted structures, it sits over the SIC domain-SUFU interface, making contacts with both the SIC domain and SUFU (Figures S19, S20). Other than these compact SUFU-binding modules and the DNA-binding domain, most of the GLI polypeptide is predicted to be unstructured/disordered (Figures 3, S3, S4A, S12), consistent with a model in which short, conditionally-folded domains embedded in intrinsic disorder form the key contacts with SUFU and other GLI-binding proteins (*102*).

To validate some of the novel predictions, we showed that the SIC domain of human GLI proteins was sufficient to bind SUFU in biochemical binding assays, and that alanine substitutions of predicted contact residues severely impaired this interaction (Figure 6). More extensively, we assessed the predicted interface between the GLI NR domain and the C-terminal lobe of SUFU by systematically testing the effect of a large number of GLI and SUFU variants – including many disease-associated variants – on GLI-SUFU complex formation (Figures 7–10). The outcome of these experiments strongly supports the AF3-predicted interfaces. Overall, this study substantially revises and extends the structural model of the GLI-SUFU interaction, and suggests several testable mechanisms by which disease mutations, competing protein-protein interactions and post-translational modifications could blunt or tune SUFU repression.

### A rigorous multi-step pipeline provides confidence in the predictions

The goal of this study was to use AlphaFold3 – augmented by experimental verification – to further delineate SUFU-binding domains of GLI transcription factors and their contact points with SUFU. Our computational procedure was to obtain (at least) 25 models for each protein pair (5 different seeds x 5 diffusion samples per seed). Protein pairs included full-length GLI and SUFU (with zinc ions to coordinate with the cysteine and histidine residues in the GLI zinc fingers) as well as GLI and SUFU variants in which unstructured/disordered regions were deleted prior to co-folding. Interfacial residues in these models were analyzed by residue-level confidence metrics, and only those surpassing stringent thresholds (and doing so consistently in multiple models) were retained as potential contacts. Finally, these provisional contacts were verified for the top 5 models of each protein-pair using the popular molecular visualization and analysis tool ChimeraX, which considers additional criteria beyond geometric proximity. Furthermore, taking advantage of the fact that there are three human GLI paralogs, we focused on contacts conserved in all three. The results of this meticulous, multistep procedure were lists of high-quality contacts for each GLI domain and the interacting residues on SUFU (Tables S3-S6).

Additional confidence in the predictions presented in this paper is provided by the following observations:

1. Standard AlphaFold global confidence metrics (ipTM, pTM, MC) are good in models of full-length GLI and SUFU, and very high in pruned models where sequences known or predicted to be intrinsically disordered were deleted prior to co-folding. Local confidence metrics (pLDDT, PAE) are high/very high for interfacial residues in all models (e.g. Figures S3, S4, S8; Table S1).
2. The replicate models resulting from different random seeds are highly consistent with each other and superimposable (Figures S5, S6).
3. The SUFU and SYGHLS elements of the predicted structures superimpose with the published co-crystal structures (Figures S7, S16; Table S2).
4. The GLI1, GLI2 and GLI3 interfaces with SUFU are highly consistent with each other and superimposable (e.g. Figures 5, S3, S14-S16).
5. The GLI1, GLI2 and GLI3 interfaces with SUFU all feature very similar high-quality contacts (Tables S3-S6).
6. The novel interfaces we identified are highly conserved at the sequence and structural level in GLI-SUFU orthologs in vertebrate and invertebrate model organisms (Figures S9-S11).
7. GLI-SUFU contact residues have substantial overlap with Ci-dSufu contact residues (Tables S3-S5, Figure S13).
8. Negative control co-foldings with GST confirmed the expected lack of interaction (Figure S12).
9. Prior studies of mutated GLI/Ci variants support the importance of the RSSL motif and the SIC domain for SUFU binding and SUFU-dependent repression *in vivo* (*72, 77, 84, 97–99, 103*). However, these studies did not provide structures for either domain, nor did they determine the corresponding contact points with SUFU.
10. We experimentally validated the AF3-predicted interaction between the C-terminal SIC domains of GLI1, GLI2 and GLI3 and SUFU (Figure 6). We further validated key predicted contact residues in the SIC domains of GLI1 and GLI2 and in the N-terminal lobe of SUFU (Figure 6, Figure S17).
11. Lastly, for the novel RSSL motif, we experimentally validated multiple key predicted contact residues in GLI1 (Figures 7, 8) and SUFU (Figures 9, 10, S23).

### The GLI NR domain comprises an expanded SUFU-binding interface

Prior structural work established that short GLI1- and GLI3-derived peptides containing the SYGHLS motif bind as a β-strand in the narrow channel between the N- and C-terminal lobes of SUFU (*66, 67*). In the AF3 models generated in this study, the SYGHLS element occupies essentially the same position as in published co-crystal structures. This result is not surprising given that AF3’s training set included these structures. However, AF3 consistently predicts an additional, extensive set of contacts contributed by an adjacent motif that we term the RSSL motif, which lies within the broader NR domain. In the predicted complexes, the RSSL motif sits over and shields the SYGHLS strand, contacting two key SUFU residues (SUFU Asp159 and Ser267) that are also contacted by SYGHLS, and also makes numerous additional contacts with the SUFU C-terminal lobe (Figures 4, 5, S14-S16, Tables S3, S4).

The AF3-predicted structures identified two residues in the GLI RSSL motif that act as “anchor” determinants of affinity: Arg100 and Leu106 in GLI1, and the corresponding, identical residues in GLI2, GLI3 and Ci. GLI1 Arg100 participates in an extended hydrogen-bonding/salt-bridge network that stabilizes the region around the SYGHLS β-strand, while GLI1 Leu106 inserts deeply into a hydrophobic pocket formed by multiple SUFU residues. Consistent with this, substitutions at GLI1 Arg100 or Leu106 dramatically weaken SUFU binding *in vitro* (Figure 7), and multiple different substitutions of SUFU pocket residues reduce binding to a small fraction of wild-type (Figures 9 and 10). Other highly-conserved residues in the RSSL motif (GLI1 Ser102, Pro103, Ser104, Ser105, Val107) also make important contacts with SUFU; mutation of these residues also compromises GLI-SUFU binding (Figure 8). Additionally, we found evidence that further binding affinity may be provided by fuzzy contacts between positively-charged residues in the N-terminal portion of the NR domain (Lys79, Lys80, Arg81 in GLI1) and a negatively-charged surface patch on SUFU (Figure 8, Figure S22).

### The conserved SIC domain is a second SUFU-binding interface

Our AF3 models and validation experiments also strongly support a second SUFU-binding interface contributed by the SUFU-interacting C-terminal (SIC) domain, which consists of the C-terminal ∼27 residues of GLI/Ci. In the predicted complexes, this region folds into a β-strand followed by an amphipathic helix; the β-strand binds the SUFU N-terminal domain via β-strand augmentation with SUFU β1, and the helix inserts hydrophobic residues into a surface groove on SUFU (Figures 6, S15, S17). The SIC interface provides putative structural insight into ‘dual-binding’ and ‘multi-binding’ models in which SUFU can engage GLI proteins through multiple regions and may not completely dissociate during pathway activation (*72, 97, 103–105*). Interestingly, there is no overlap between SUFU residues that contact the SIC domain and those that contact the NR domain, consistent with the possibility of independent regulation (Tables S3-S5, Figure S13).

Unexpectedly, the predicted SIC–SUFU interaction closely resembles the co-crystal structure of the SUFU-binding site (SBS) of Drosophila Fused bound to dSufu. In support of this structural similarity, we detected previously unrecognized sequence similarity between the Fused SBS and the GLI/Ci SIC domain (Figure S18). This new finding may inform models of the structure, function and phosphoregulation of the Ci-dSufu-Fused complex (GLI-SUFU-ULK3/STK36 in humans) (*72, 84, 85, 93, 103, 104, 106, 107*).

### The SR motif may help explain GLI2/GLI3-specific regulatory behaviors

In the GLI2-SUFU and GLI3-SUFU structures – but not in GLI1-SUFU – AF3 predicts a third interface involving a short motif located in the N-terminal repressor domains of GLI2 and GLI3. This “SR” motif is predicted to contact both the folded SIC domain and residues in the N-terminus of SUFU. The SR motif was not present in the Ci-dSufu structure, and is generally absent from *Drosophilidae* Ci orthologs. It is, however, widespread in Gli/Ci orthologs from other invertebrates, including other insects (Figures S9-S11). Although we did not experimentally validate the SR-SUFU contact in the current work, the prediction is intriguing because it provides a potential structural rationale for functional differences among GLI paralogs. GLI2 and GLI3 (unlike GLI1) undergo regulated processing and possess well-characterized repressor activities; it is therefore plausible that GLI2/3 evolved/retained an additional SUFU-coupled interaction that helps coordinate repression, processing, or trafficking. If the SR motif indeed engages SUFU preferentially when the SIC domain is present (as the models suggest, Figures S19, S20), then perhaps this interaction regulates repression or processing when SUFU is bound to full-length GLI2/3, but not when GLI2/3 are in their truncated repressor forms.

### Some SUFU mutations differentially affect GLI and GLIS3 binding

SUFU is not exclusively a HH-pathway regulator; it also binds and regulates GLIS3, a transcription factor with important developmental roles and disease relevance. GLIS3 shares similarity to GLI proteins in the 5ZF and NR domains, and it contains a YGH sequence aligned with the GLI SYGHLS motif. In our biochemical tests, GLIS3 bound SUFU with low-nanomolar affinity. Surprisingly, however, many SUFU variants that strongly impaired GLI binding had only modest effects on GLIS3 binding (Figure 10, Figure S23). This suggests that some *SUFU* mutations may differentially affect Hedgehog signaling versus GLIS3-regulated programs, perhaps impacting disease symptoms.

On a related note, in addition to its well-known role as a tumor suppressor, SUFU has also been shown to have tumor-promoting activities (*17, 108*). It will be interesting to explore whether certain SUFU missense variants may diminish some SUFU functions while maintaining others.

### Disease-associated SUFU variants of residues contacting the NR domain often behave as binding-defective alleles

A key question, both mechanistically and clinically, is which SUFU missense variants compromise SUFU repression by disrupting physical binding to GLI proteins, which compromise SUFU function through other mechanisms (e.g., localization, stability, recruitment of additional regulators), and which are effectively silent. When we tested an extensive panel of SUFU substitutions (including multiple patient mutations), ∼80% of AF3-predicted contact-residue substitutions caused substantial defects in GLI1 binding. We then tested a subset of these against GLI2, GLI3, and GLIS3, and found that all of them – including 7 of 8 patient/tumor mutations tested – also substantially reduced binding to GLI2 and GLI3 (Figures 9, 10, S23). A practical implication is that AF3-derived structural context may help triage *SUFU* missense variants into those likely to broadly impair GLI binding (and therefore likely to be pathogenic) versus those which are benign or which may perturb SUFU biology by other mechanisms. Of course, such inference must be done cautiously, because a residue can be structurally “contacting” yet not energetically critical, and because cells can partially buffer against weakened interactions. Nonetheless, the strong concordance between predicted contacts and biochemical phenotypes suggests that structure-guided interpretation could be a powerful complement to variant scoring based on other methods (*109, 110*).

In this regard, it should be noted that most variants associated with genetic diseases are missense mutations, and most missense mutations are classified as ‘variants of uncertain clinical significance’ (*110*). Improving this situation would be beneficial for patients for many reasons, including the fact that screening recommendations are based on the suspected pathogenicity of the mutation. For example, in the case of *SUFU* mutations, the International Society of Pediatric Oncology recommends infant screening for medulloblastoma and meningioma for SUFU pathogenic variant carriers (*111*).

### Conclusion and future directions

The structures presented herein highlight the role of SUFU as a multi-interface scaffold/adaptor protein whose binding partners exploit different combinations of surfaces to achieve distinct regulatory logics. Furthermore, they provide a structural framework that will be useful for understanding existing experimental findings and suggesting new hypotheses.

As explicated in the results, mutations of some of the contact residues identified and verified in this work have been shown in other studies to result in Hedgehog signaling defects *in vivo*. Nevertheless, important future directions include (1) assessing the effects of mutations in additional contact residues – particularly the patient mutations – in cells and animal model organisms; (2) using the ability to precisely disrupt GLI-SUFU contact interfaces to delineate the combinatorial role of the SR, NR and SIC-domain interactions in SUFU/GLI signaling; (3) exploring at scale how AlphaFold3 and related tools may assist in the prediction of disease variant effects in diseases where perturbations to macromolecular interactions are known to drive pathogenesis.

## Materials and Methods

### Genes

The human genes used in this study were human *GLI1* (NCBI accession number NM_005269), *GLI2* (NM_001371271), *GLI3* (NM_000168), *GLIS3* isoform b (NM_001438909), and *SUFU* (NM_016169). *D. melanogaster* Ci (P19538) and dSufu (Q9VG38) protein sequences are the canonical sequences from UniProt.

### Plasmids

The vector used for generating glutathione-*S*-transferase (GST) fusion proteins was pGEX-LB, a derivative of pGEX-4T-1. In pGEX-LB, an encoded Pro residue is replaced with a Gly-Gly-Gly-Gly-Gly-Ser-Gly coding sequence to promote the independent functioning of the GST and fusion moieties (*112*). Plasmid GST-SUFU encodes a fusion of GST to full-length human SUFU (*77*). The construction of plasmid GST-GLI1(68-232), which encodes a fusion of GST to residues 68-232 of human GLI1, was previously described, as was that of plasmid GST-GLI3(279-480) (*78*); plasmids GST-GLI2(218-433) and GST-GLIS3(105-284) were constructed using a similar procedure. Plasmids GST-GLI1-SIC, GST-GLI2-SIC and GST-GLI3-SIC were constructed by inserting the C-terminal 27 amino acids plus the stop codon of each of the human GLI proteins, flanked *by BamH*I and *Sal*I restriction sites, into pGEX-LB.

Plasmid pGEM3Z-SUFU encodes full-length (residues 1-484) human SUFU downstream of a promoter for T7 RNA polymerase, and was constructed by subcloning from pGEM4Z-SUFU (*77*); plasmid T7-GLI1(1-232) is also described in this previous study.

### Protein purification

GST fusion proteins were expressed in bacteria, purified by affinity chromatography using glutathione-Sepharose, eluted from the matrix by reduced glutathione, dialyzed to remove the glutathione, quantified by Bradford assay and Nanodrop analysis, and flash frozen in aliquots at -80 °C, as described elsewhere (*78, 112, 113*).

### *In vitro* transcription and translation

Proteins labeled with [^35^S]-methionine were produced by coupled transcription and translation reactions (T7, Promega). Translation products were partially purified by ammonium sulfate precipitation (*114*), and resuspended in ‘Binding Buffer’ (20 mM Tris-HCl, pH 7.5, 125 mM KOAc, 0.5 mM EDTA, 5 mM DTT, 0.1% (v/v) Tween20, 12.5% (v/v) glycerol) prior to use in binding assays.

### Protein binding assays

For the experiments shown in Figs 7 and 8, ^35^S-radiolabeled GLI1_1-232_ protein and variants thereof were prepared by *in vitro* translation and partially purified by ammonium sulfate precipitation. Approximately 1 pmol of ^35^S-GLI1 was added to each 200 μL binding reaction; 10% of this amount was loaded in the ‘Input’ lane. Purified GST-SUFU was added at concentrations that varied from 0.625 to 40 nM. The buffer for binding reactions was Binding Buffer (described above) to which 1 mg/mL molecular-biology grade bovine serum albumin was added to block non-specific protein-protein or protein-bead interactions. The binding reactions were incubated for 15 min at 30 °C. Then, glutathione-Sepharose beads (20 μl of a 50% slurry) were added, the samples were rocked for 1 h at room temperature, and the resulting bead-bound protein complexes were isolated by sedimentation, washed thoroughly with binding buffer to remove unbound protein, and resolved by 10% SDS-PAGE on the same gel. Other details of the protein binding assays are as described elsewhere (*112, 113*).

For the experiments shown in Figs 6, 9, 10, S17 and S23, ^35^S radiolabeled full-length human SUFU protein and variants thereof were prepared by *in vitro* translation and partially purified by ammonium sulfate precipitation. Approximately 1 pmol of ^35^S-SUFU was added to each 200 μL binding reaction; 10% of this amount was loaded in the ‘Input’ lane. The buffer for binding reactions was Binding Buffer + BSA, as described above. The binding reactions were incubated for 15 min at 30 °C, then rocked for 1 h at room temperature. The resulting bead-bound protein complexes were isolated by sedimentation, washed thoroughly with binding buffer to remove unbound protein, and resolved by 10% or 12% SDS-PAGE on the same gel.

Quantification of binding was performed on a Typhoon TRIO+ Imager using phosphor imaging mode. Percent binding was determined by comparing the input with the amount that was co-sedimented.

### Statistical analysis and fitting

Statistical analysis of binding assay results was performed using Welch’s ANOVA test with planned two-way comparisons. Multiple hypothesis testing was accounted for using Dunnett’s T3 multiple comparison test. The GraphPad Prism software package was used for statistical analysis.

Statistical analysis of normalized data was performed as described elsewhere (*115*); briefly, significance was assessed based on the distance between the mutant variant mean and the normalized value of 100, measured in units of the 95% confidence interval (CI) of the mean of the variant.

Binding isotherm analysis for Figures 7 and 8 was conducted as follows: Emax and Kd for the wild-type protein were determined by fitting wild-type data from each experiment to the formula

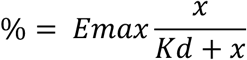

where ‘%’ is the percent of the radiolabeled protein bound, and *x* is the concentration of GST-SUFU. The Emax for wild-type was then used to normalize the wild type and all mutant variants analyzed in the same experiment.

For the experiments shown in Figures 6, 7, 8 and S17, Kd values for each concentration were determined from the formula

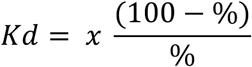

where ‘%’ and *x* are as defined above.

### Computed structure modeling tools

Our computational pipeline using the AlphaFold server is described in the main text. pLDDT values and PAE values of interacting residues were calculated with the aid of the Predictomes server (*116*). Chord diagrams were made using the AlphaBridge server on default confidence settings (*75*). PAE graphics for Figure S8 were made using PAE viewer (*117*). Molecular graphics and analyses were performed with UCSF ChimeraX (*118*).

## Glossary of AlphaFold-related confidence metrics

### 1. Global Confidence Metrics

**pTM** (predicted Template Modeling score) is an integrated measure of how well AlphaFold has predicted the overall structure of the complex, based on comparing the predicted structure and the hypothetical true structure. pTM scores range from 0 to 1; a pTM score above 0.5 means the overall predicted fold for the complex might be similar to the true structure (*119*).

**ipTM** (interface pTM) evaluates the accuracy of predicted protein-protein complex interfaces. ipTM scores range from 0 to 1; values higher than 0.8 represent confident high-quality predictions, values below 0.6 suggest a failed prediction, and values between 0.6 and 0.8 are a gray zone. Disordered regions and regions with low pLDDT score may negatively impact the ipTM score even if the structure of the complex is predicted correctly.

**MC** (Model Confidence) is a combined metric equal to ipTM x 0.8 + pTM x 0.2 (*120*). MC scores range from 0 to 1; values of 0.8 or above indicate confident high-quality predictions.

### 2. Local Confidence Metrics

**pLDDT** (predicted Local Distance Difference Test) is a per-residue measure of AF’s confidence in the local positioning of that residue. It is scaled from 0 to 100, with higher scores indicating higher confidence and a likely more accurate prediction. Note that AF3 calculates pLDDT values per atom, but the Predictomes server used in our pipeline averages these over the whole amino acid residue.

**PAE** (Predicted Aligned Error) is a measure of how confident AF is in the relative position of two amino acid residues within the predicted structure, including across interfaces. PAE is defined as the expected positional error at residue X, measured in Ångströms (Å), if the predicted and hypothetical true structures were aligned on residue Y. For residues at the interface of a protein-protein interaction, a PAE of less than 5 indicates that AF3 is confident that the two residues in question are within 5 Å of each other. AF3 calculates PAE values per token (e.g. per atom), but the Predictomes server used in our pipeline converts these to per-residue-pair values.

## Acknowledgements

This work was supported by the National Cancer Institute of the National Institutes of Health under award number P30CA062203, by the UC Irvine Chao Family Comprehensive Cancer Center using Anti-Cancer Challenge funds, and by a UC Irvine Chancellor’s Discretionary award. Additional research funds were provided by the UC Irvine Charlie Dunlop School of Biological Sciences and Department of Developmental and Cell Biology.

## Author Contributions (CRediT format)

AJ Bardwell: formal analysis, investigation, methodology, supervision, validation, visualization, writing.

U Arif: investigation.

A Salgado: investigation.

F Muhammad: investigation.

CN Salinas Auris: investigation.

L Bardwell: conceptualization, formal analysis, funding acquisition, investigation, methodology, project administration, supervision, validation, visualization, writing.

