## Supplementary Materials for "An expanded structural and biochemical map of GLI-SUFU recognition identifies binding-defective disease variants"


| Figure or Table | Page | Title/description |
|---|---|---|
| Figure S1 | 2 | 5ZF domain superposition |
| Figure S2 | 3 | Variable positioning of the 5ZF domain in different CSMs |
| Figure S3 | 4 | Chord diagrams of additional computed structure models |
| Table S1 | 5 | Model confidence metrics |
| Figure S4 | 6 | pLDDT values of selected computed structure models |
| Figure S5 | 7 | Comparison of GLI1/2-SUFU models from 5 different seeds |
| Figure S6 | 8 | Comparison of GLI3-SUFU and Ci-dSufu models from 5 seeds |
| Table S2 | 9 | RMSD values from superposition of CSMs and x-ray structures |
| Figure S7 | 10 | Superposition of GLI CSMs with empirical co-crystal structures |
| Figure S8 | 11 | Predicted aligned error (PAE) diagrams |
| Figure S9 | 12,13 | Sequence alignment of SUFU-binding domains/motifs from select vertebrate and invertebrate GLI orthologs |
| Figure S10 | 14 | Chord diagrams of additional GLI-SUFU computed structure models from selected vertebrate and invertebrate species |
| Figure S11 | 15,16 | Comparison of AF3-predicted GLI-SUFU structures of select vertebrate and invertebrate species |
| Figure S12 | 17 | GST does not interact with GLI or SUFU in control co-foldings |
| Table S3 | 18 | GLI-SUFU contact residues in RSSL motif |
| Table S4 | 19 | GLI-SUFU contact residues in SYGHLS motif |
| Table S5 | 20 | GLI-SUFU contact residues in SIC domain |
| Table S6 | 21 | GLI-SUFU contact residues in SR motif |
| Figure S13 | 22,23 | Alignment of human and fruit fly SUFU with contact residues shown |
| Figure S14 | 24 | GLI1 and GLI2 RSSL motif H-bonds; burying of SYGHLS by RSSL |
| Figure S15 | 25 | Comparison of GLI1, GLI2 and GLI3 CSMs |
| Figure S16 | 26 | SYGHLS motif H-bonds in GLI1, GLI3 CSMs vs. x-ray structures |
| Figure S17 | 27 | Substitutions of SIC domain contact residues with alanine abolish SIC-SUFU binding |
| Figure S18 | 28 | Ci and Fused bind to the same elements of dSufu |
| Figure S19 | 29 | SR motif structure and interactions |
| Figure S20 | 30 | SR motif is unstructured in co-foldings lacking the SIC domain |
| Figure S21 | 31 | Alignment of the NR domains of GLI1, GLI2, GLI3 and Ci |
| Figure S22 | 32 | Fuzzy electrostatic interaction of GLI1 K79, K80, R81 with SUFU |
| Figure S23 | 33 | Additional SUFU mutations assessed for GLI and GLIS3 binding |
| Figure S24 | 34,35 | Similarity of GLIS3 and GLI1 interfaces with SUFU |
| Figure S25 | 36 | Chord diagrams of additional computed structure models |


Version 11/9/2026

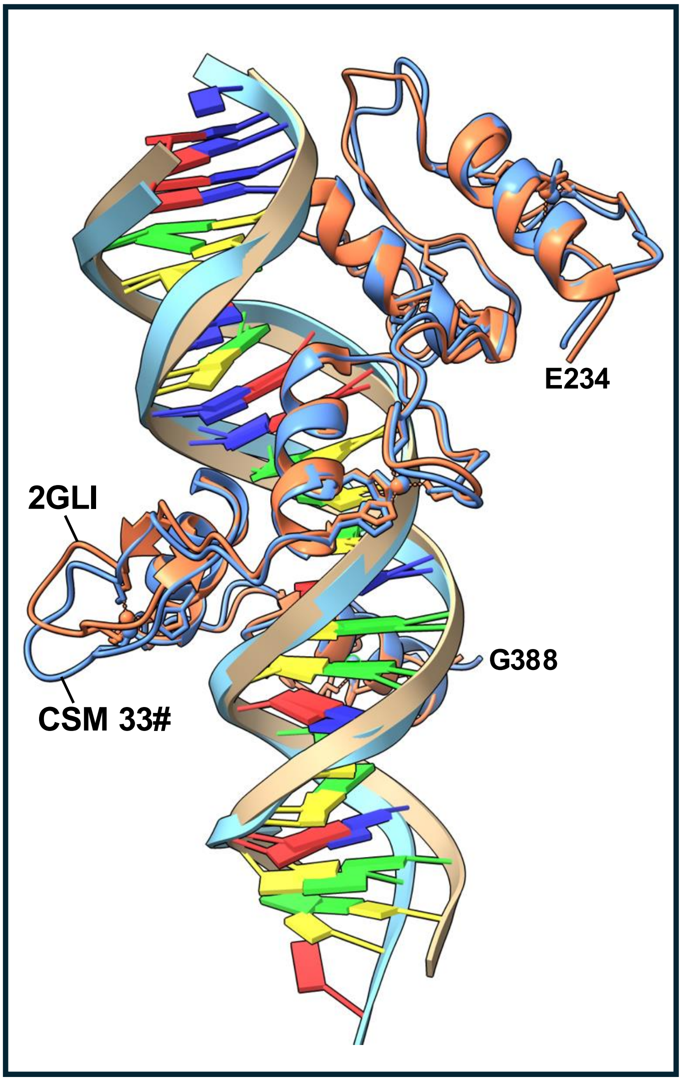


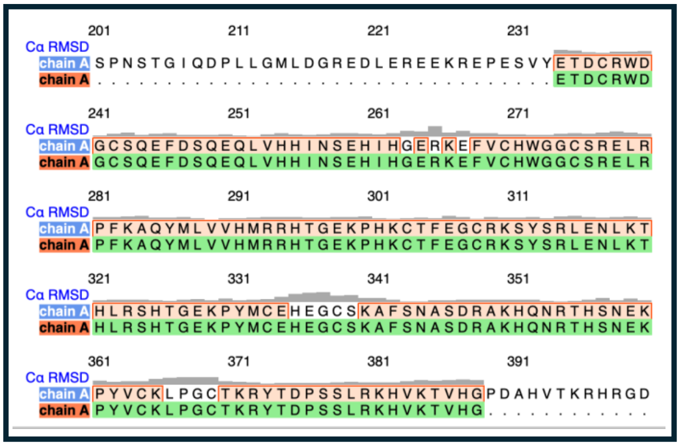


**Figure S1**. Superposition of x-ray and AF3-predicted structures of GLI1 5-zinc finger DNA-binding domain bound to target DNA sequence.

Structural superposition of x-ray structure 2GLI with residues 234-388 of GLI1 from Computed Structure Model (CSM) 33#, GLI1 5z SUFU. This was an AlphaFold3 co-folding of full-length human GLI1 with full-length human SUFU and 5 zinc ions.

The computed Cα root mean squared deviation (RMSD) between 143 pruned atom pairs is 0.886 angstroms; across all 155 pairs: 1.260 angstroms.

CSM 33# was generated by cofolding the following components using the AlphaFold server:

1. Full-length human GLI1 (1106 residues)
2. Full-length human SUFU (484 residues)
3. 5 zinc ions
4. DNA strand **5′-TTCGTCTTGGGTGGTCCACG**
5. DNA strand **3′-AAGCAGAACCCACCAGGTGC**

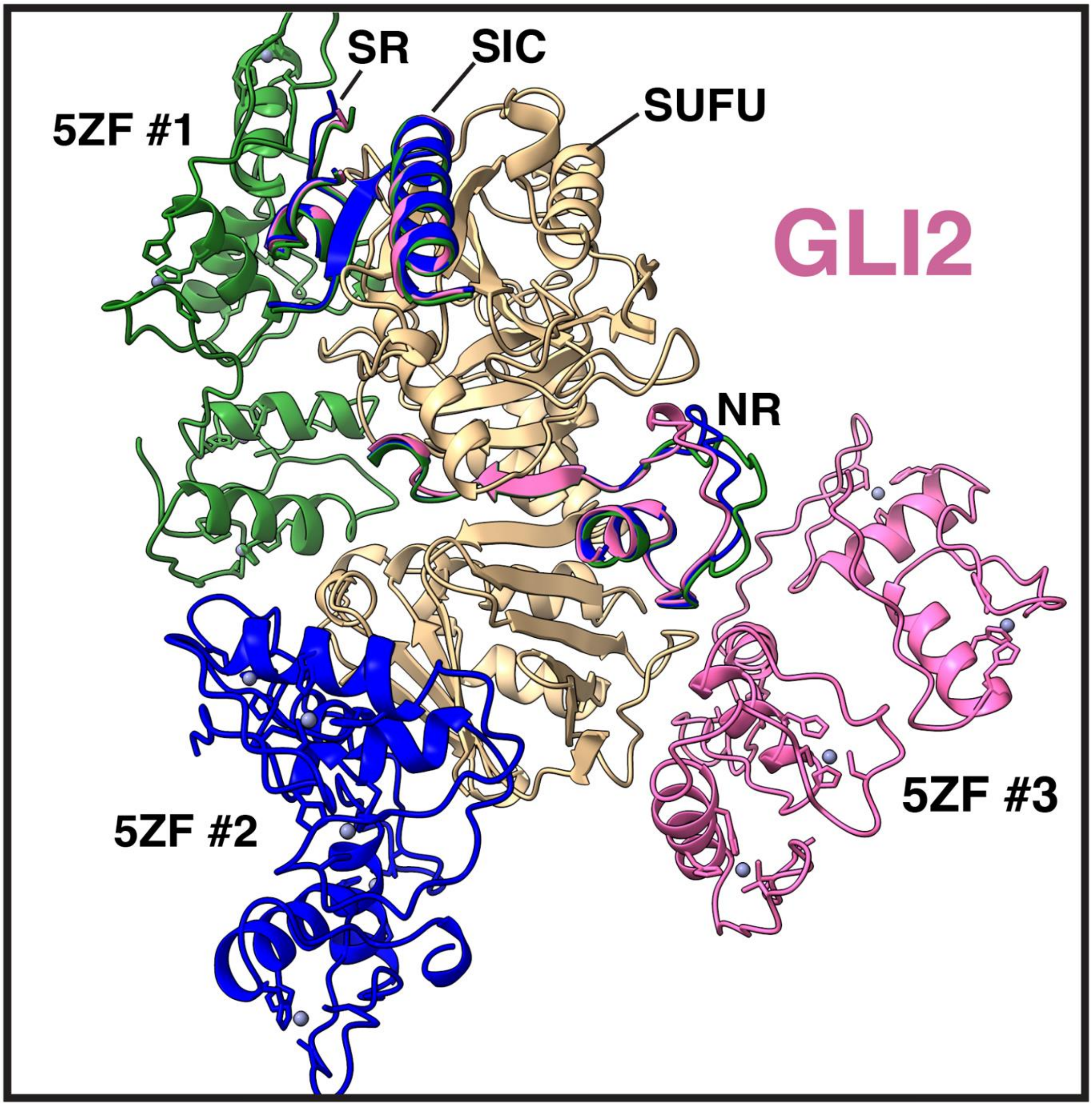


**Figure S2**. Variable positioning of the 5 zinc finger (5ZF) DNA-binding domain of GLI2 in three different GLI2 5z SUFU computed structure models.

Top models from 3 different seeds of GLI2 FL 5z SUFU (colored green, blue and pink) were superimposed with respect to SUFU (tan). The SR, NR (RSSL-loop-SYGHLS), 5ZF and SIC domains of GLI2 are shown. The rest of the GLI2 sequence (disordered and low confidence) is hidden. Also hidden are the disordered regions of SUFU (residues 1-25 and 279-362). In two additional models (not shown), the 5ZF domain was placed in the approximate position of 5ZF #2.

Models are CSM 91# (pink), 92# (green) and 267# (blue)

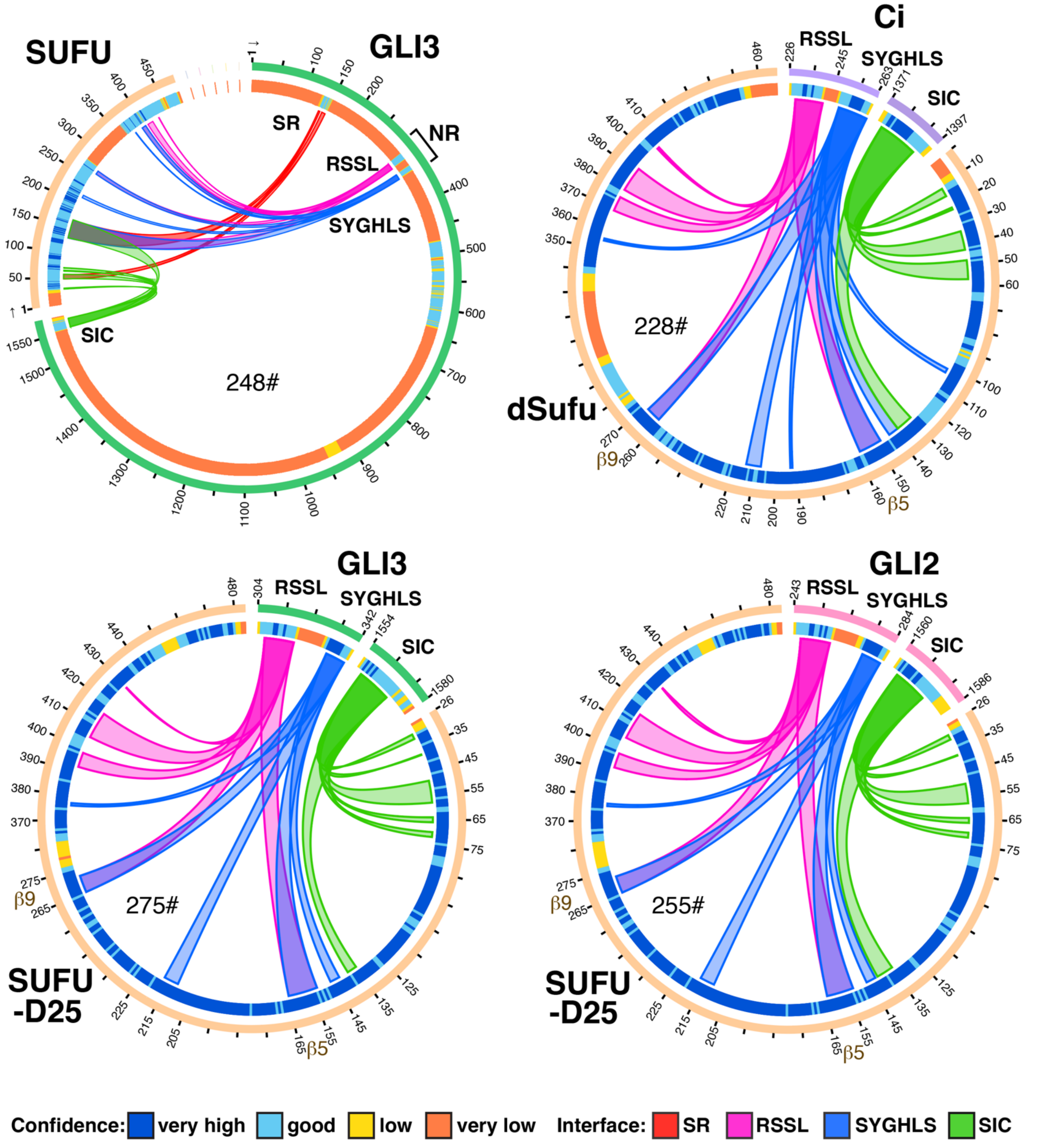


**Figure S3**. Chord diagrams of additional computed structure models:

1. CSM 248#, full-length human GLI3 co-folded with full-length human SUFU and 5 zinc ions.
2. CSM 228#, *D. melanogaster* Ci residues 226-263 and 1371-1397 co-folded with full-length dSufu.
3. CSM 275#, GLI3 residues 304-342 and 1554-1580 co-folded with SUFU-D25.
4. CSM 255#, GLI2 residues 243-284 and 1560-1586 co-folded with SUFU-D25.

Chord Diagrams were generated with AlphaBridge (https://alpha-bridge.eu/) using default confidence settings and then enhanced with Adobe Illustrator.

| SUFU / GLI1 | WT | WT | WT | d60 | d60 | d60 | D | D | D | D25 | D25 | D25 |
|---|---|---|---|---|---|---|---|---|---|---|---|---|
| | ipTM | pTM | MC | ipTM | pTM | MC | ipTM | pTM | MC | ipTM | pTM | MC |
| GLI1 5z DNA | 0.73 | 0.42 | 0.67 | 0.81 | 0.43 | 0.73 | 0.81 | 0.42 | 0.73 | 0.85 | 0.41 | 0.76 |
| GLI1 5z | 0.77 | 0.42 | 0.70 | 0.84 | 0.41 | 0.76 | 0.86 | 0.41 | 0.77 | 0.89 | 0.42 | 0.80 |
| GLI1 NRC (93-128) | 0.87 | 0.77 | 0.85 | 0.85 | 0.85 | 0.85 | 0.86 | 0.87 | 0.86 | **0.90** | **0.91** | **0.90** |

| SUFU / GLI2 | WT | WT | WT | D25 | D25 | D25 |
|---|---|---|---|---|---|---|
| | ipTM | pTM | MC | ipTM | pTM | MC |
| GLI2 5z DNA | 0.76 | 0.39 | 0.69 | 0.85 | 0.38 | 0.76 |
| GLI2 5z | 0.78 | 0.38 | 0.70 | **0.91** | 0.38 | 0.80 |
| GLI2 NRC (243-280) | 0.84 | 0.77 | 0.83 | **0.90** | **0.91** | **0.90** |
| GLI2 SIC (1560-1586) | 0.85 | 0.70 | 0.82 | 0.82 | 0.75 | 0.81 |
| GLI2 NRC & SIC | 0.85 | 0.78 | 0.83 | **0.91** | **0.91** | **0.91** |

**Key**

| | |
|---|---|
| (pink) | = low confidence |
| (white) | = moderate confidence |
| (green) | = high confidence |
| **0.90** | = very high confidence |

| SUFU / GLI3 | WT | WT | WT | D25 | D25 | D25 |
|---|---|---|---|---|---|---|
| | ipTM | pTM | MC | ipTM | pTM | MC |
| GLI3 5z DNA | 0.77 | 0.38 | 0.69 | 0.86 | 0.38 | 0.76 |
| GLI3 5z | 0.78 | 0.39 | 0.71 | **0.91** | 0.38 | 0.80 |
| GLI3 NRC (304-341) | 0.83 | 0.77 | 0.82 | 0.89 | **0.90** | 0.89 |
| GLI3 SIC (1554-1580) | 0.82 | 0.70 | 0.80 | 0.80 | 0.74 | 0.79 |
| GLI3 NRC & SIC | 0.83 | 0.78 | 0.82 | **0.90** | **0.91** | **0.90** |

| dSufu / Ci | WT | WT | WT |
|---|---|---|---|
| | ipTM | pTM | MC |
| Ci 5z | 0.71 | 0.36 | 0.64 |
| Ci NRC (226-263) | 0.85 | 0.80 | 0.84 |
| Ci SIC (1371-1397) | 0.87 | 0.79 | 0.85 |
| Ci NRC & SIC | 0.86 | 0.86 | 0.86 |

**Table S1**. Model confidence metrics
Average of metrics for the top 3 computed structure models (CSM's) for each protein pair.
Each CSM included in the averaging is from a different job with a distinct random seed.

Abbreviations:

<u>AlphaFold metrics (see Glossary in Methods for definition)</u>
ipTM = interface predicted Template Modeling score
pTM = predicted Template Modeling score
MC = Model Confidence, equal to 0.8 x ipTM + 0.2 x pTM

<u>GLI</u>
GLI 5z DNA = full-length human GLI co-folded with 5 zinc ions, DNA target site oligonucleotide, and the indicated SUFU variant.
GLI 5z = full-length human GLI co-folded with 5 zinc ions and the indicated SUFU variant.
NRC = C-terminal half of NR domain, containing both the RSSL and SYGHLS motifs. Residue numbers in parenthesis.
SIC = SIC domain. The residue numbers that make up this domain are given in parenthesis.

<u>SUFU</u>
WT = wild type human SUFU (484 amino acids) or fly dSufu (468 aa's)
d60 = SUFUdelta60 derivative used in co-crystalization studies by Zhang et al.
D = SUFU-D derivative used in co-crystalization studies by Cherry et al.
D25 = SUFU-D with N-terminal 25 amino acid deleted

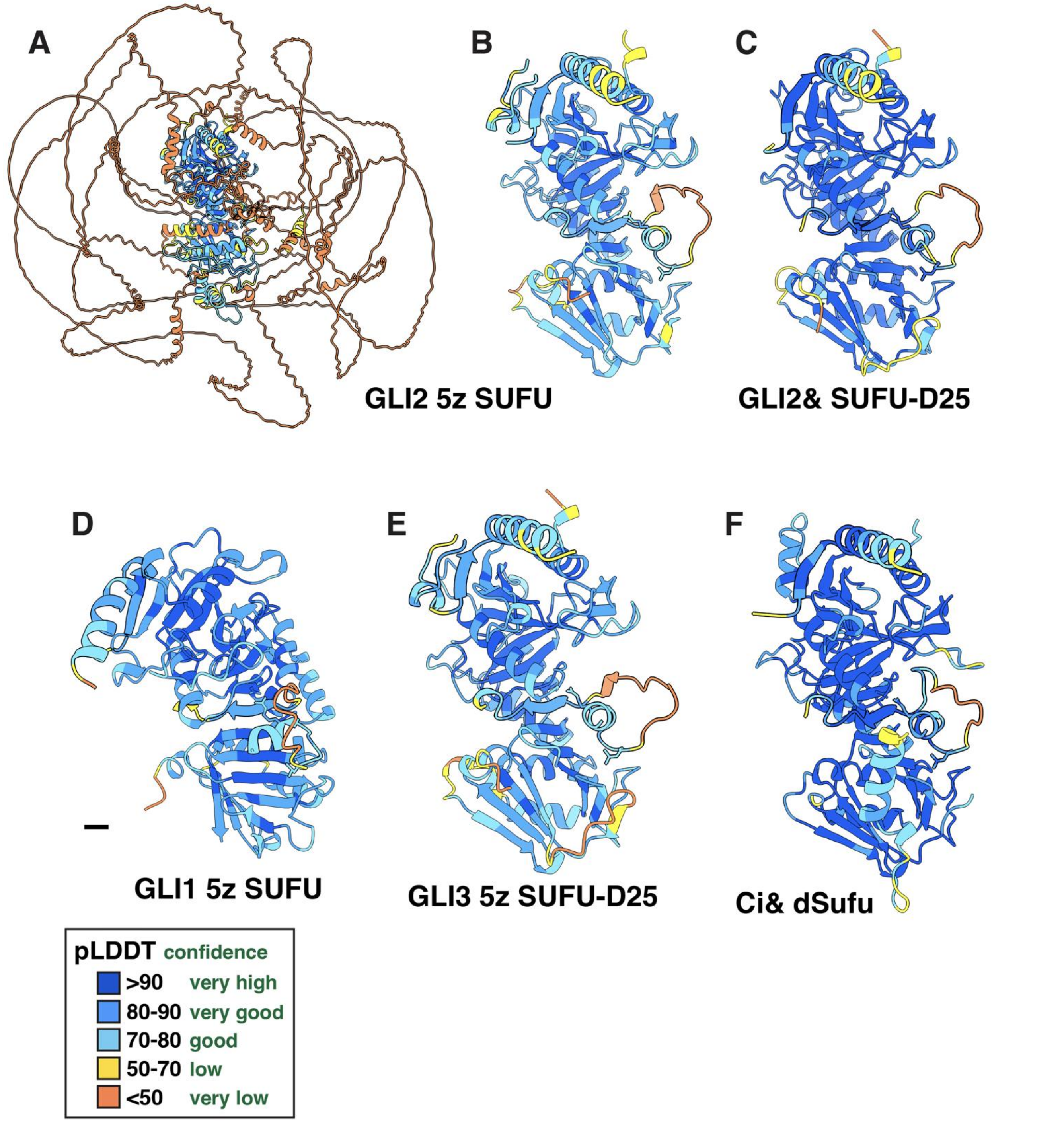


**Figure S4.** pLDDT values of selected computed structure models.

**A** CSM 92# GLI2 5z SUFU, all residues shown.
**B** CSM 92# GLI2 5z SUFU, with GLI2 1-60, 76-244, 281-1560 and SUFU 1-25 and 279-362 hidden.
**C** CSM 255# GLI2 243-280 & 1560-1586 SUFU-D25.
**D** CSM 261# GLI1 5z SUFU, with GLI1 1-94, 129-1080 and SUFU 1-25 and 279-362 hidden; scalebar = 5 A°.
**E** CSM 190# GLI3 5z SUFU-D25, with GLI3 1-133, 149-305 and 342-1554 hidden.
**F** CSM 228# Ci 226-263 & 1371-1397 dSufu, dSufu 1-11, 300-338 and 455-468 hidden.

Side chains are shown for GLI1 R100 and L106 and homologous residues in other CSMs.

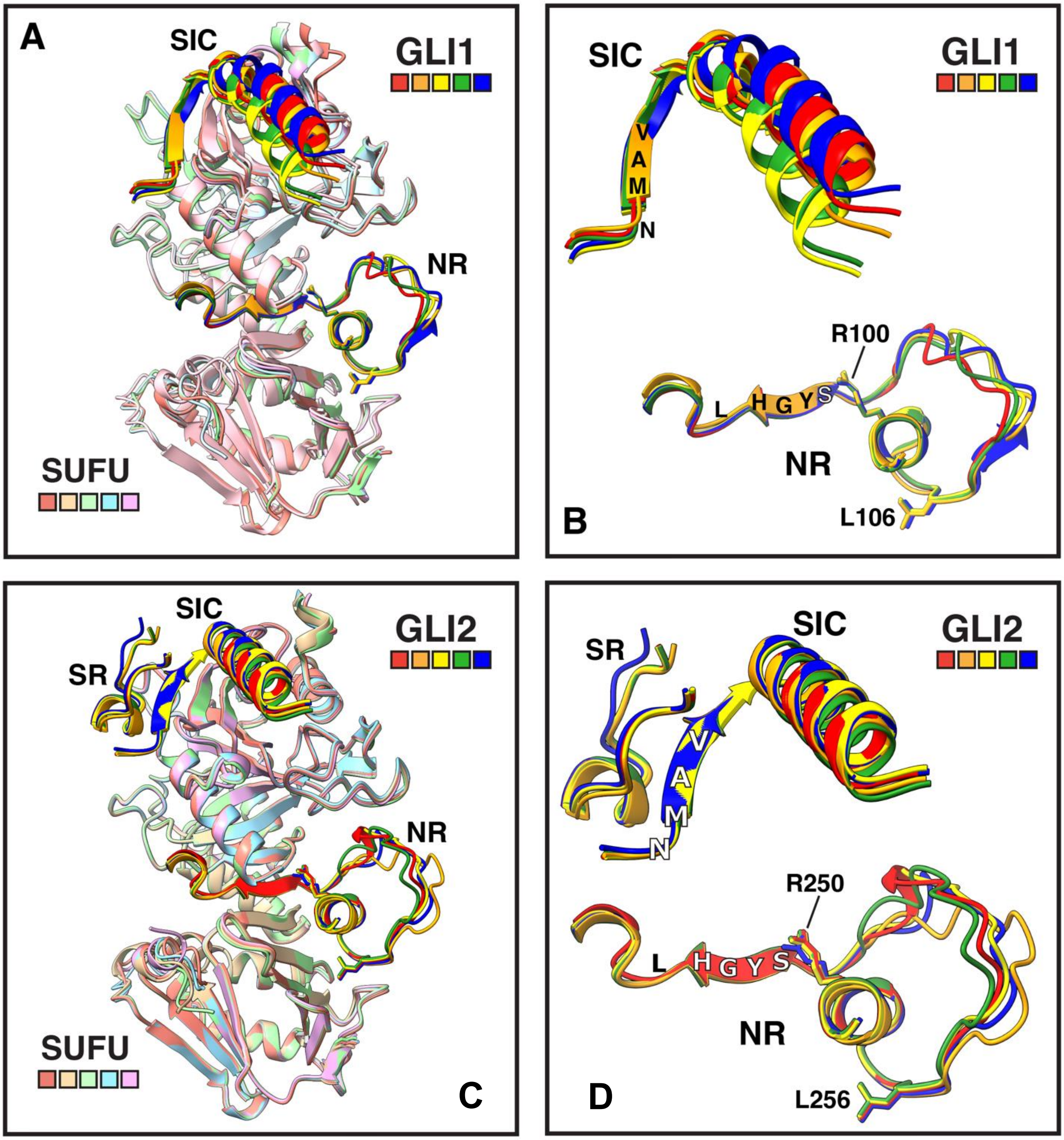


**Figure S5**. Comparison of GLI1-SUFU and GLI2-SUFU models from five different seeds.

**A** The top scoring models from 5 independent seeds of GLI1 5z SUFU were superimposed with respect to the SUFU structures. In each model, GLI1 is a primary color (red, orange, yellow, green or blue) and SUFU is a pastel color. The RSSL, SYGHLS and SIC domains of GLI1 are shown, the rest of the sequence (disordered and low confidence) is hidden. R100 and L106 in the RSSL domain are shown in stick representation. Also hidden are the disordered regions of SUFU (residues 1-25 and 279-362). Models are CSM 31#, 111#, 126#, 261#, and 262#.
**B** Same as A but with all of SUFU hidden. The GLI1 domains are also shown enlarged, and some key conserved amino acid residues are also indicated.
**C** Same as A, but for GLI2 5z SUFU. The SR domain of GLI2 is also shown. Models are CSM 91#, 92#, 264#, 266# and 267#.
**D** Same as B, but for GLI2 5z SUFU.

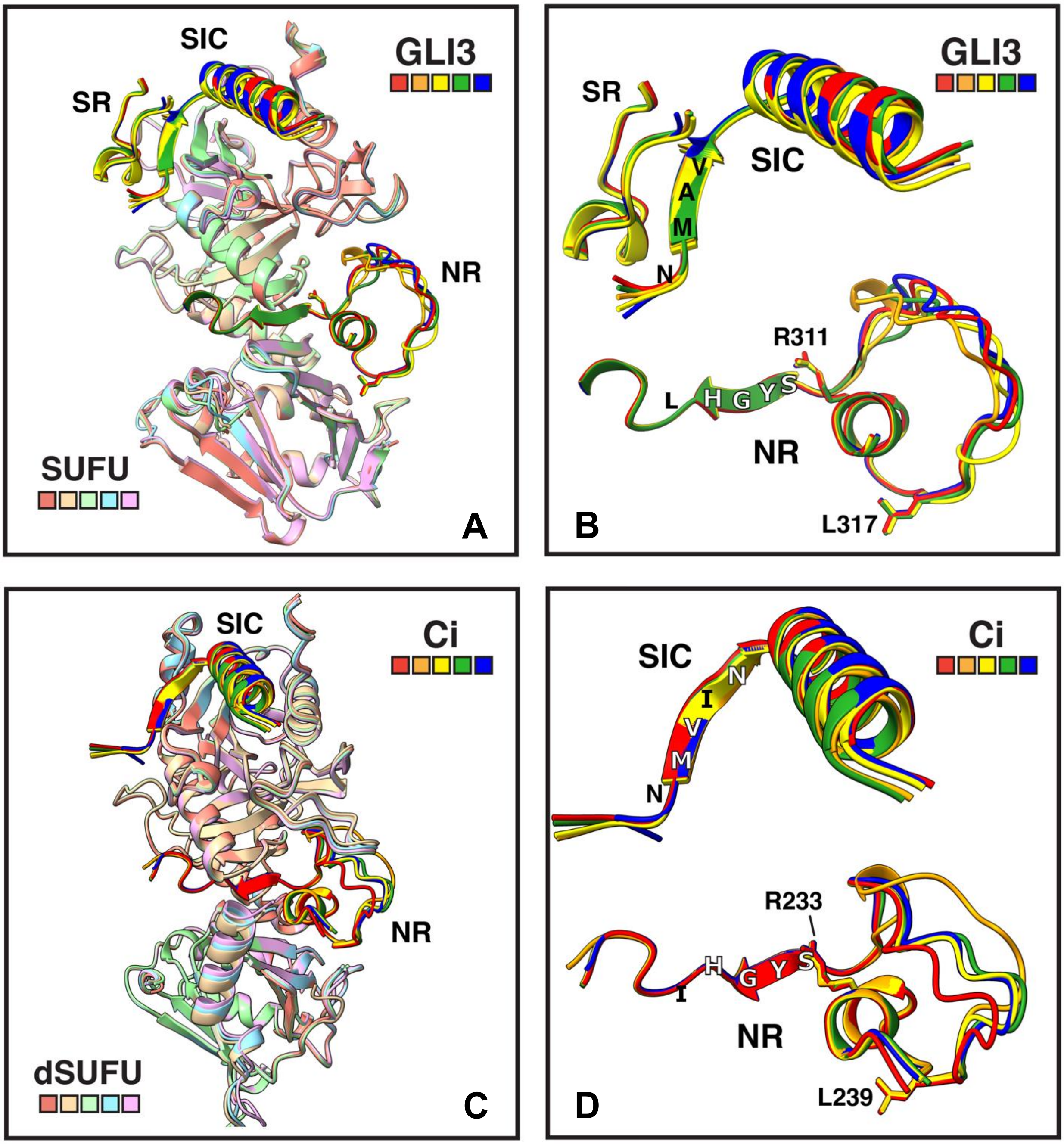


**Figure S6**. Comparison of GLI3-SUFU and Ci-dSufu models from five different seeds.

**A** The top scoring models from 5 independent seeds of GLI3 5z SUFU (primary colors: red, orange, yellow, green and blue) were superimposed with respect to the SUFU structures (pastel colors). The SR, RSSL, SYGHLS and SIC domains/motifs of GLI3 are shown, the rest of the sequence (disordered and low confidence) is hidden. Also hidden are the disordered regions of SUFU (residues 1-25 and 279-362). The side chains of R311 and L317 in the RSSL domain are shown. Models are CSM 188#, 189#, 248#, 249# and 268#.

**B** Same as A, but with all of SUFU hidden. The GLI3 domains are also shown enlarged, and some key conserved amino acid residues are also indicated.

**C** Same as A, but for Ci 226-263 & 1371-1397 dSufu. Models are CSM 226#, 228#, 229#, 230# and 265#.

**D** Same as B, but for Ci 226-263 & 1371-1397 dSufu.

**Compare SUFU to SUFU**

| | GLI1p SUFU | GLI3p SUFU | GLI1 5z SUFU | GLI2 5z SUFU | GLI3 5z SUFU |
|---|---|---|---|---|---|
| | 4BLB | 4BLD | CSM 261# | CSM 92# | CSM 248# |
| 4KMD | **0.6** | **0.7** | **0.5** | **0.7** | **0.6** |
| GLI1p SUFU | 334 | 302 | 356 | 357 | 358 |
| 4BLB | | **0.8** | **0.6** | **0.6** | **0.5** |
| GLI1p SUFU | | 353 | 340 | 341 | 339 |
| 4BLD | | | **0.9** | **0.7** | **0.8** |
| GLI3p SUFU | | | 331 | 336 | 334 |

Top number is RMSD
Bottom number is number of residue pairs compared (Cα atoms only)

**Compare SYGHLS to SYGHLS**

| | GLI1p SUFU | GLI3p SUFU | GLI1 5z SUFU | GLI2 5z SUFU | GLI3 5z SUFU |
|---|---|---|---|---|---|
| | 4BLB | 4BLD | CSM 261# | CSM 92# | CSM 248# |
| 4KMD | **0.8** | **1.0** | **1.1** | **1.2** | **1.2** |
| GLI1p SUFU | 46 | 46 | 46 | 46 | 46 |
| 4BLB | | **0.6** | **1.4** | **1.4** | **1.4** |
| GLI1p SUFU | | 46 | 46 | 46 | 46 |
| 4BLD | | | **1.4** | **1.3** | **1.4** |
| GLI3p SUFU | | | 46 | 46 | 46 |

Top number is RMSD
Bottom number is number of **atom** pairs compared

**Compare SYGHLS to SYGHLS**

| | GLI1p SUFU | GLI3p SUFU | GLI1 5z SUFU | GLI2 5z SUFU | GLI3 5z SUFU |
|---|---|---|---|---|---|
| | 4BLB | 4BLD | CSM 261# | CSM 92# | CSM 248# |
| 4KMD | **0.4** | **0.6** | **0.8** | **0.9** | **0.9** |
| GLI1p SUFU | 6 | 6 | 6 | 6 | 6 |
| 4BLB | | **0.5** | **0.8** | **0.9** | **0.9** |
| GLI1p SUFU | | 6 | 6 | 6 | 6 |
| 4BLD | | | **0.9** | **0.8** | **0.9** |
| GLI3p SUFU | | | 6 | 6 | 6 |

Top number is RMSD
Bottom number is number of residue pairs compared (Cα atoms only)

**Table S2.** Root mean squared deviation (RMSD) values (A°) when selected computed structure models (CSMs) are superimposed on empirical models 4KMD, 4BLB and 4BLD from the Protein Data Bank.

For any given comparison, the SUFU chain from the CSM was superimposed upon the SUFU chain from the co-crystal structure using the 'matchmaker' tool in ChimeraX, which reports the RMSD for the alpha carbons (top panel). The GLI chains were not independently aligned, but rather co-transposed with the SUFU chains. Then the RMSD values for the predicted vs. empirical SYGHLS residues were computed using the 'rmsd' command in ChimeraX, both for all atoms (middle panel) and for Cα atoms only (bottom panel).

RMSD values of less than 2 (white cells) are considered very good superposition.
RMSD values of less than 1 (green cells) are considered excellent superposition.

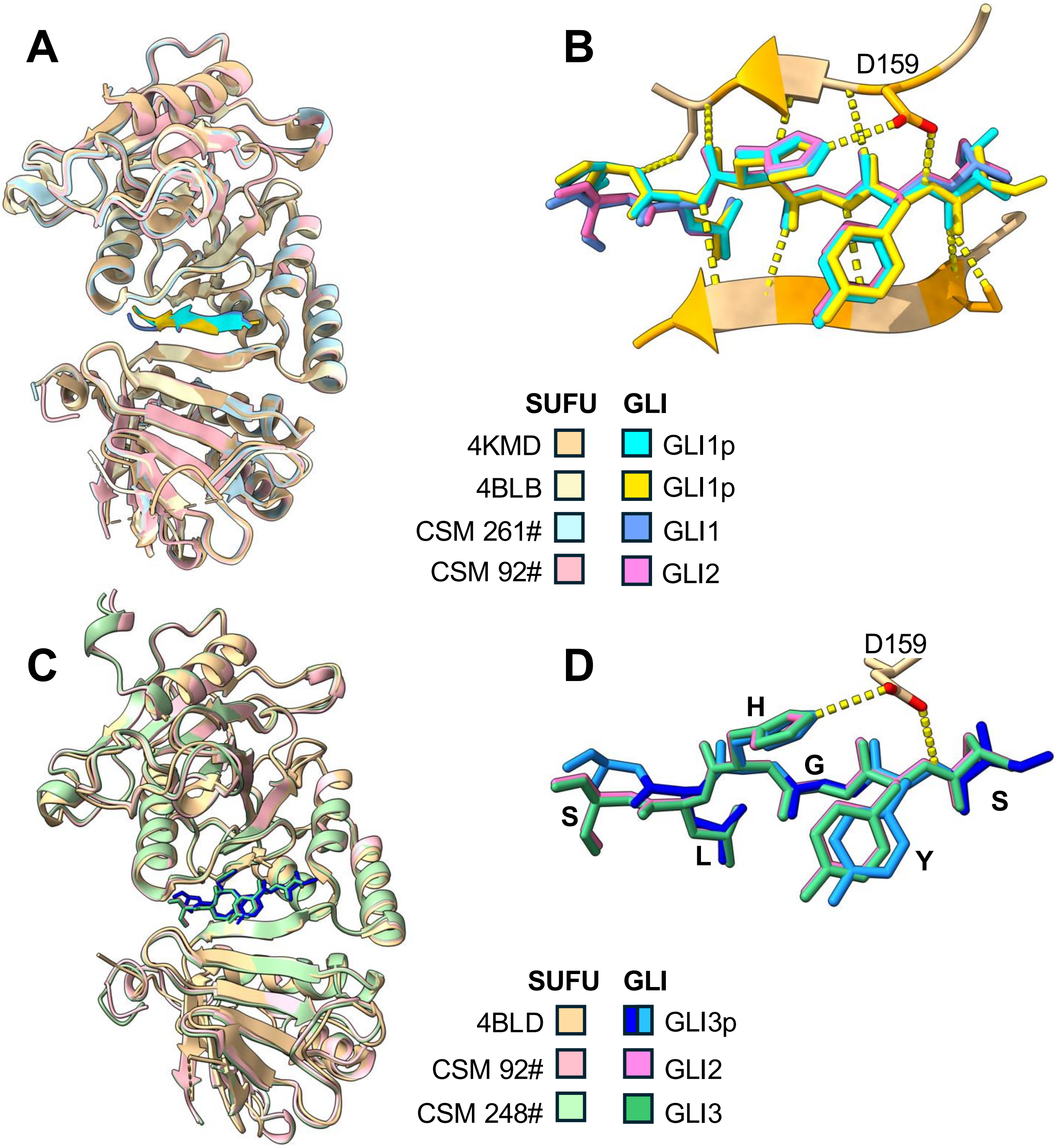


**Figure S7**. Structural superposition of GLI CSMs with empirical co-crystal structures from the protein data bank (PDB).

**A** Superposition of SUFU chains and SYGHLS peptides from PDB 4BLB, CSM 261# and CSM 92# with PDB 4KMD. The GLI chains were not independently aligned, but rather moved with the SUFU chains.

**B** Similar to **A,** but zoomed in on the SYGHLS motif, with side chains and key H-bonds shown.

**C** SUFU chains and SYGHLS motifs from CSM 92# and CSM 248# superimposed on PDB 4BLD.

**D** Similar to **C,** but zoomed in on the SYGHLS motif, with H-bonds from SUFU D159 shown.

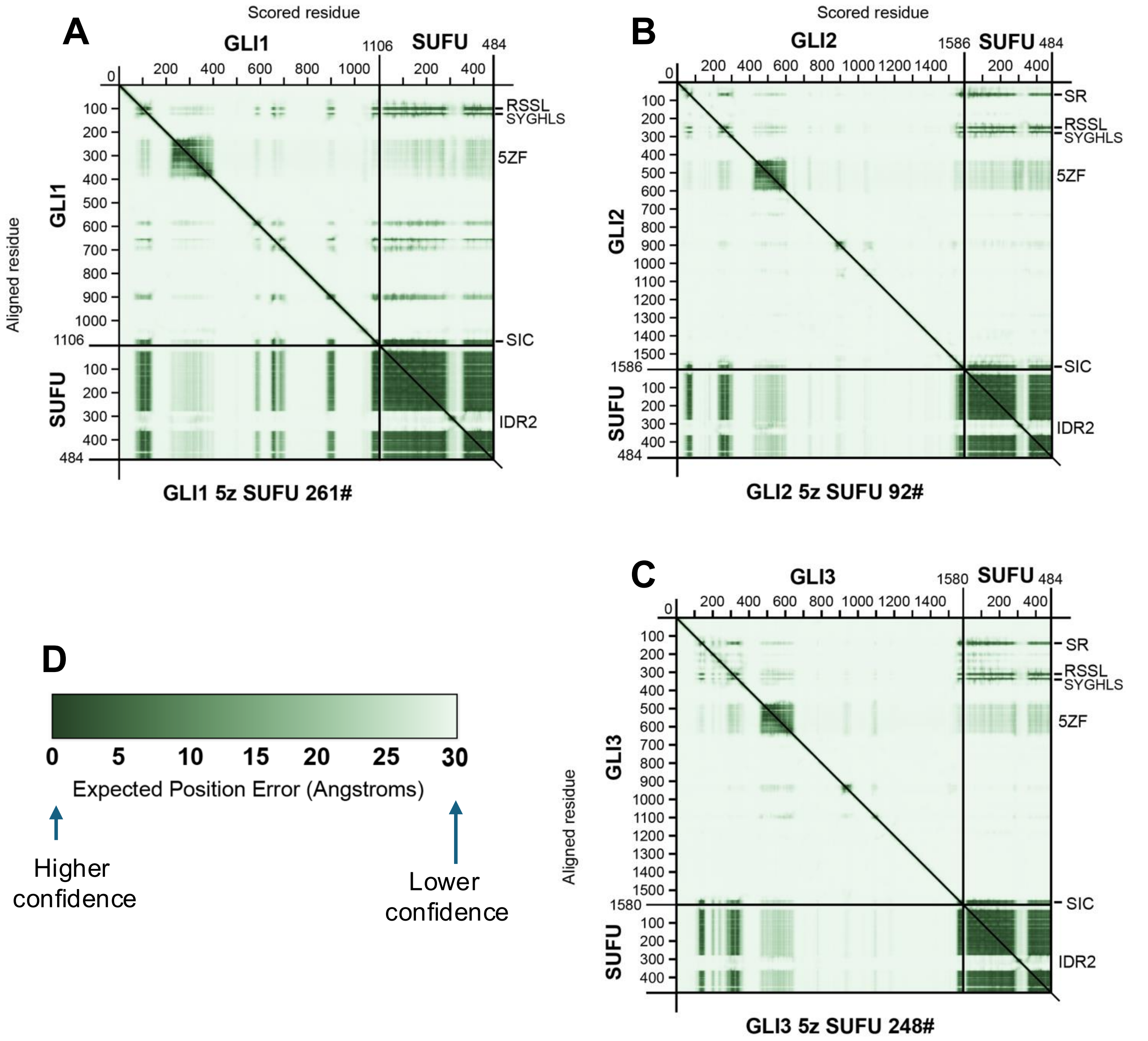


**Figure S8**. Predicted aligned error (PAE) diagrams.

**A** CSM 261# GLI1 5z SUFU
**B** CSM 92# GLI2 5z SUFU
**C** CSM 248# GLI3 5z SUFU
**D** Key; darker colors indicate lower predicted position error.

PAE diagrams are 2D plots that estimate the error in the relative position and orientation between two tokens (in this case, amino acid residues) in the predicted structure. The color at coordinates x,y represents the predicted position error of residue x if the predicted and true structures were aligned on residue y. Lower PAE values (darker green color) suggest well-defined relative positions and orientations in the prediction.

**A**

**SR motif**

| Phylum/Class | Species | Common Name | Protein | Accession # | Residues | Contacts | SR motif |
|---|---|---|---|---|---|---|---|
| | | | | | | w SUFU | ...●...●●.●.●.● |
| | | | | | | w SIC | ●●.●...●....... |
| Mammal | *H.sap* | human | GLI3 | NP_000159 | 134-148 | | PVPIDARHHEGRYHY |
| Mammal | *H.sap* | human | GLI2 | NP_001358200 | 61-75 | | PLPIDMRHQEGRYHY |
| Mammal | *M.mus* | mouse | Gli2 | NP_001074594 | 57-71 | | PLPIDMRHQEGRYHY |
| Bird | *G.gal* | chicken | Gli2 | NP_001258830 | 56-70 | | PLPIDMRHQEGRYHY |
| Amphibian | *X.tro* | frog | Gli2 | XP_002943329 | 52-66 | | PFPVEVRHQEGRYHY |
| Bony fish | *D.rer* | zebrafish | Gli2a | NP_571042 | 49-63 | | PIPIDMRHHEGRYHY |
| Echinoderm | *S.pur* | urchin | Gli/Ci | XP_030856399 | 10-24 | | PIPVNSRTHEGRYHF |
| Mollusc | *H.ruf* | abalone | Gli/Ci | XP_046357464 | 123-137 | | PLPINVSTLEGRYHW |
| Annelid | *O.fus* | tubeworm | Gli/Ci | CAH1773043 | 121-135 | | PMPINAHTHDGRYHW |
| Arthropod | *L.sac* | silverfish | Gli/Ci | XP_082967008 | 35-49 | | PIPVDQRTHEGRYIW |
| Arthropod | *D.mel* | fruit fly | Ci | NP_524617 | NA | | no SR found |
| Cnidarian | *N.vec* | anemone | Gli/Ci | XP_048579969 | NA | | no SR found |

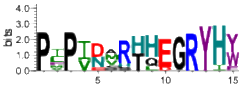

**B**

**NR domain**

| Species | Protein | Residues | D-site | RSSL | loop | SYGHLS |
|---|---|---|---|---|---|---|
| | | | | ..●....●●.●●●●●● | | .●●●●●●●●.. |
| human | GLI1 | 78-129 | TKKRALSISPLSDAS- | LDLQTVIRTSPSSLV | AFI----NSRCTSPG- | GSYGHLSIGTM |
| human | GLI3 | 289-342 | SRKRTLSISPLSDHS- | FDLQTMIRTSPNSLV | TIL---NNSRSSSSAS | GSYGHLSASAI |
| human | GLI2 | 228-281 | SRKRALSISPLSDAS- | LDLQRMIRTSPNSLV | AYI---NNSRSSSAAS | GSYGHLSAGAL |
| mouse | Gli2 | 224-277 | SRKRALSISPLSDAS- | LDLQRMIRTSPNSLV | AYI---NNSRSSSAAS | GSYGHLSAGAL |
| chicken | Gli2 | 225-278 | SRKRALSISPLSDAS- | IDLQTMIRTSPNSLV | AYI---NNSRSSSAAS | GSYGHLSAGTI |
| frog | Gli2 | 219-272 | SRKRALSISPLSDAS- | IDLQTMIRTSPNSLV | AYI---NNSRSSSAAS | GSYGHLSAGAI |
| zebrafish | Gli2a | 213-266 | SRKRALSISPVSDAS- | IDLQTMIRTSPNSLV | AYI---NNSRSSSAAS | SSYGHLSVGAI |
| *S.pur* | Gli/Ci | 163-216 | SRKRALSISPLFNES- | IDINAMIRTSPNSLV | AYV---NNSRSSSAAS | GSYGHLSAGTV |
| *H.ruf* | Gli/Ci | 266-321 | NRKRALSQSPISDY-- | LDIQSLTRSSEGSLQ | LTPFLHHNSRSSSAAS | GSYGHLSAASL |
| *O.fus* | Gli/Ci | 261-315 | SRKRALSHSPISDY-- | LDIQSLTRSSEGSLQ | LTPLM-HHSRSSSAAS | GSYGHLSAASF |
| *L.sac* | Gli/Ci | 197-248 | SRKRALSSSPYSDS-- | FDINSMIRFSPNSLV | SIV---NGSRSSSAS- | GSYGHLSAGAL |
| *D.mel* | Gli/Ci | 212-264 | SRKRALSSSPYSDS-- | FDINSMIRFSPNSLA | TIM---NGSRGSSAAS | GSYGHISATAL |
| *N.vec* | Gli/Ci | 202-254 | TRKRARSNTPASSVES | IDLNLLIRNSPDSLL | GGLI---GTRLSSA-- | GSFGHLSPIGF |

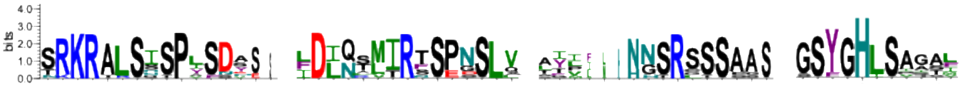

**C**

**SIC domain**

| Species | Protein | Residues | SIC domain |
|---|---|---|---|
| | | | .●●●●●●●●.●●...●........... |
| human | GLI1 | 1081-1106 | PNMAVGNMSVLLRSLPGETEFLNSSA. |
| human | GLI3 | 1555-1580 | TNMAIGDMSSLLTSLAEESKFLAVMQ. |
| human | GLI2 | 1561-1586 | SNMAVGDMSSMLTSLAEESKFLNMMT. |
| mouse | Gli2 | 1519-1544 | SNMAVGDMSSMLTSLAEESKFLNMMT. |
| chicken | Gli2 | 1503-1528 | SNMAIGDMSSMLTTLAEESKFLNMMS. |
| frog | Gli2 | 1444-1469 | GNMAIGDMSSMLTTLAEESKFLNLMP |
| zebrafish | Gli2a | 1414-1438 | GNMAIGDMSSMLTALAEESKFLLNMM |
| *S.pur* | Gli/Ci | 1274-1298 | SNMAVTDMSSMLTSLAEENRFLSIM-. |
| *H.ruf* | Gli/Ci | 1532-1556 | NNMAVNDMSSVLTTLAEENKYLNNMK. |
| *O.fus* | Gli/Ci | 1425-1450 | SNMVVNDMSSMLTTLAEENQYLSIRR. |
| *L.sac* | Gli/Ci | 1612-1637 | SNMVINDMTSSLTSLLEENRYLQMIQ. |
| *D.mel* | Gli/Ci | 1372-1397 | TNMVINDMTTSLTSLLEENRYLQMMQ |
| *N.vec* | Gli/Ci | 1082-1107 | GNMVVNDMNTLLNSLAEEDKYLEMRQ |

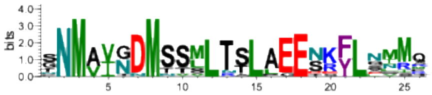

**Figure S9** (previous page). Sequence alignment of SUFU-binding domains/motifs from select vertebrate and invertebrate GLI orthologs.

Note that most invertebrates have only a single GLI ortholog.

**A** (Top) SR motif alignment. Residues that contact SUFU are indicated with cyan or magenta circles as in Figures 4 and 6. Select conserved amino acids are colored as follows: positive/basic residues (K,R,H) blue; negative/acidic (D,E) red, aliphatic hydrophobic (V,I,L,M) green, aromatic (Y,W,F) purple, amides and histidine (N, Q, H) teal.
Species compared (top to bottom) are *Homo sapiens, Mus musculus, Gallus gallus, Xenopus tropicalis, Danio rerio, Strongylocentrotus purpuratus, Haliotis rufescens, Owenia fusiformis, Lepisma saccharinum, Drosophila melanogaster, Nematostella vectensis.*
(Bottom) Sequence logo derived from the above sequences, excluding the mouse and chicken Gli2 sequences, which are 100% identical to human GLI2 in this region.

**B** (Top) NR domain alignment. The NR domain has been sub-divided into 4 parts, consisting of the MAPK-docking site region (D-site), the RSSL motif, the connecting loop, and the SYGHLS motif. Residue coloring as above, with the addition that known phosphorylation sites for proline-directed kinases are colored pink.
(Bottom) Sequence logos derived from the above sequences, excluding the mouse, chicken and frog Gli2 sequences, which are over 90% identical to human GLI2 in this region.

**C** (Top) SIC domain alignment.
(Bottom) Sequence logo derived from the above sequences, excluding the mouse and chicken Gli2 sequences, which are 90% identical to human GLI2 in this region.

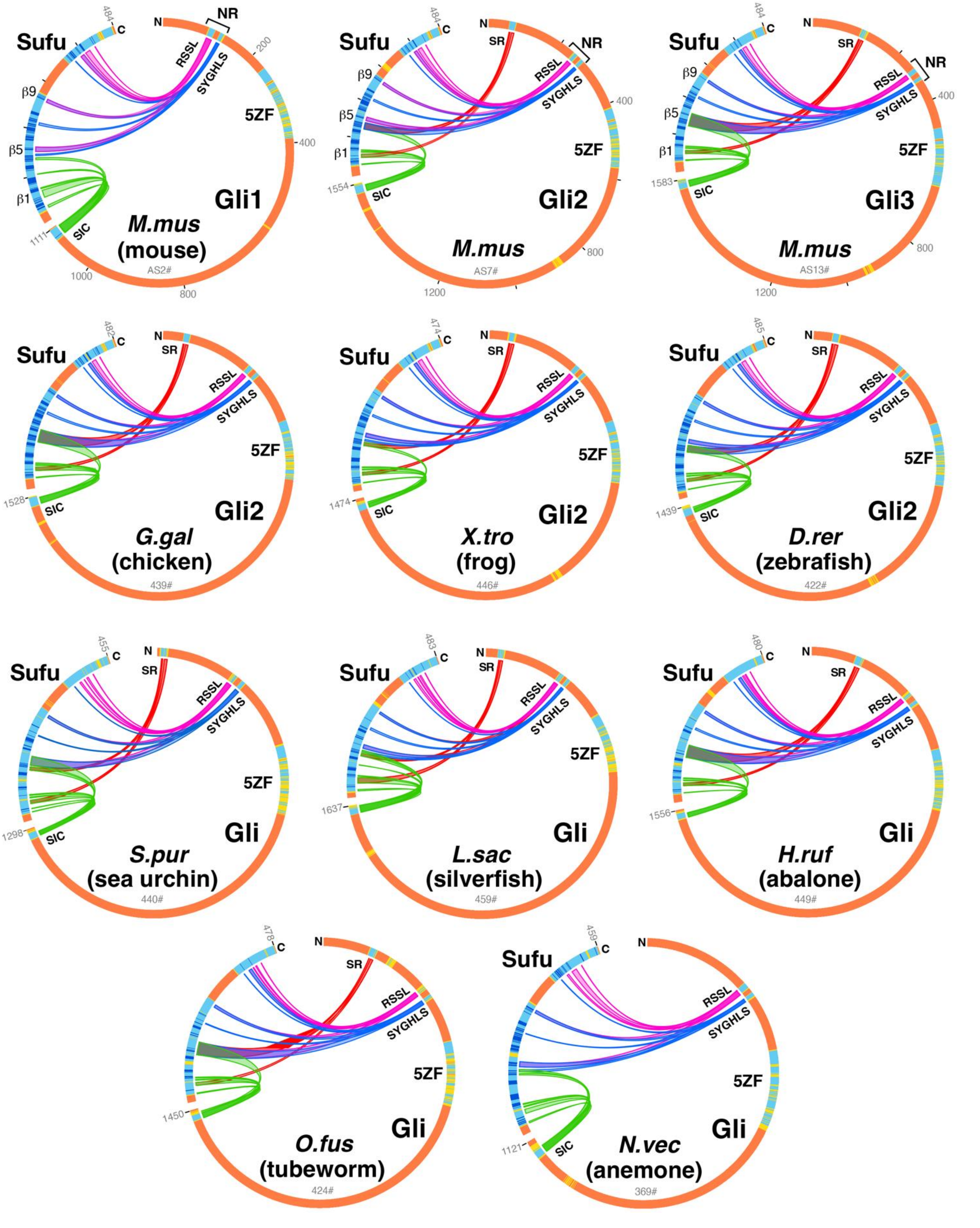


**Figure S10**. Chord diagrams of additional GLI-SUFU computed structure models from selected vertebrate (top 6) and invertebrate (bottom 5) species. Coloring as in Figure S3. The CSM# is shown below the name. Accession numbers can be found in Figure S9.

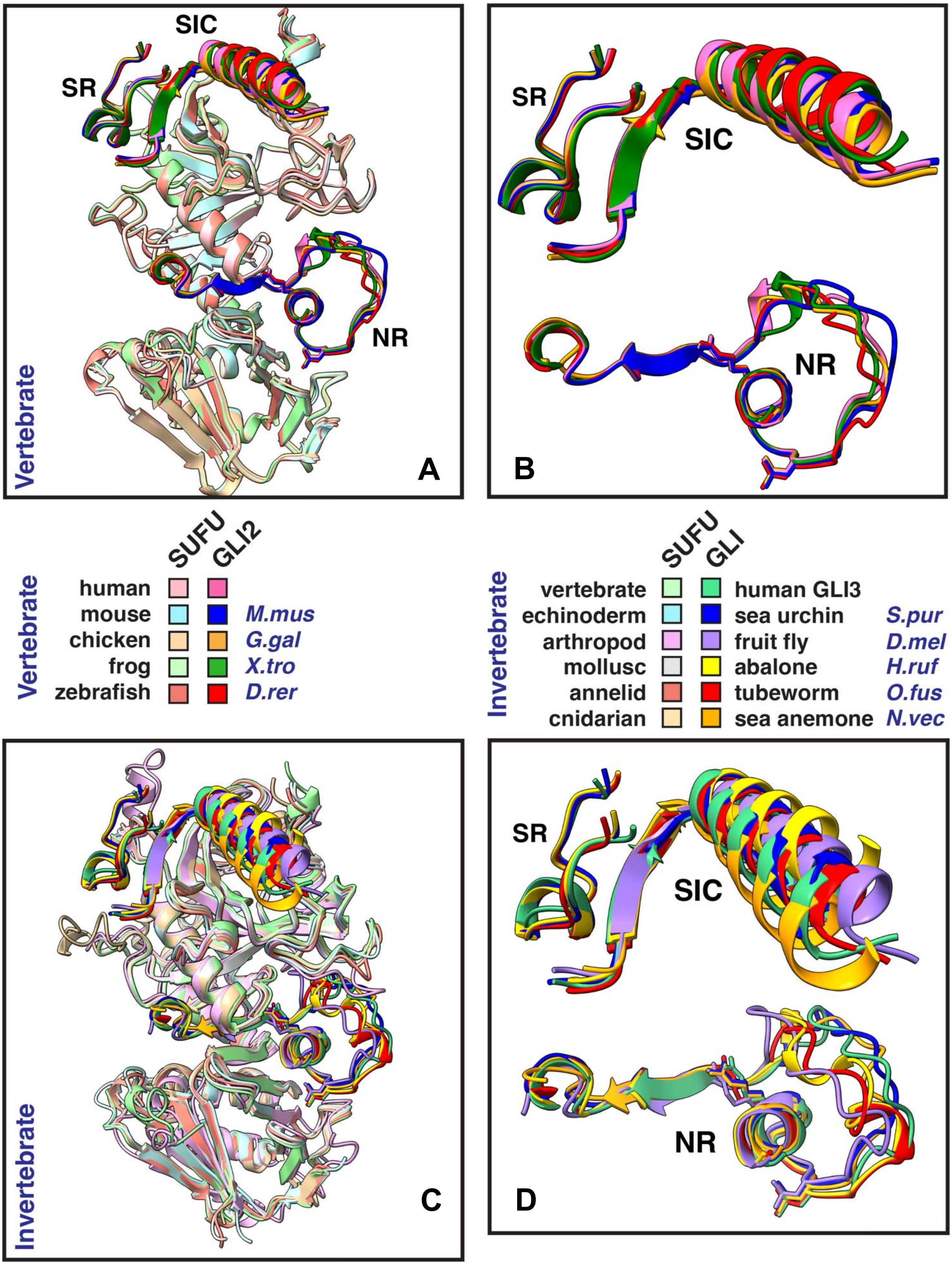
SIC
SR
NR
Vertebrate
A
B
SUFU
GLI2
Vertebrate
human
mouse
chicken
frog
zebrafish
M.mus
G.gal
X.tro
D.rer
SUFU
GLI
Invertebrate
vertebrate
echinoderm
arthropod
mollusc
annelid
cnidarian
human GLI3
sea urchin
fruit fly
abalone
tubeworm
sea anemone
S.pur
D.mel
H.ruf
O.fus
N.vec
Invertebrate
C
D

**Figure S11** (previous page). Comparison of AF3-predicted GLI-SUFU structures of select vertebrate and invertebrate species.

**A** The top scoring GLI2-5z-SUFU computed structure models from 4 vertebrate species (shown in key below) were structurally aligned with respect to their SUFU structures to human GLI2-5z-SUFU 92#. The average RMSD of the other vertebrate SUFUs with human SUFU was 0.44 over 370 $\alpha$-carbon atom pairs.

For the GLI2 models, only the SR, NR and SIC domains are shown. Also, the side chains of human GLI2 Arg250 and Leu256 are shown in stick representation, as are the corresponding Arg and Leu residues in the other species. For the SUFU models, regions corresponding to IDR1 and IDR2 are hidden.

**B** Same as A, but with all of SUFU hidden and the GLI2 domains zoomed.

**C** The top scoring Gli/Ci-SUFU computed structure models from 5 invertebrate species representing diverse phyla (shown in key below) were structurally aligned with respect to their Sufu structures to GLI3-5z-SUFU 248#. The average RMSD of the invertebrate Sufu structures with human SUFU was 0.75 over 330 $\alpha$-carbon atom pairs.

For the Gli/Ci models, only the SR, NR and SIC domains are shown. Also, the side chains of human GLI3 Arg311 and Leu317 are shown in stick representation, as are the corresponding Arg and Leu residues in the other species. For the Sufu models, regions corresponding to IDR1 and IDR2 are hidden.

**D** Same as C, but with all of Sufu hidden and the Gli/Ci domains zoomed.

**A**

| Protein 1 | Protein 2 | Other | | ipTM | pTM | MC |
|---|---|---|---|---|---|---|
| GST | GLI1 | 5 zinc | | 0.41 | 0.30 | 0.39 |
| GST | GLI2 | 5 zinc | | 0.45 | 0.29 | 0.42 |
| GST | SUFU | – | | 0.19 | 0.45 | 0.24 |
| GST | SUFU-D25 | – | | 0.20 | 0.48 | 0.26 |

= low confidence

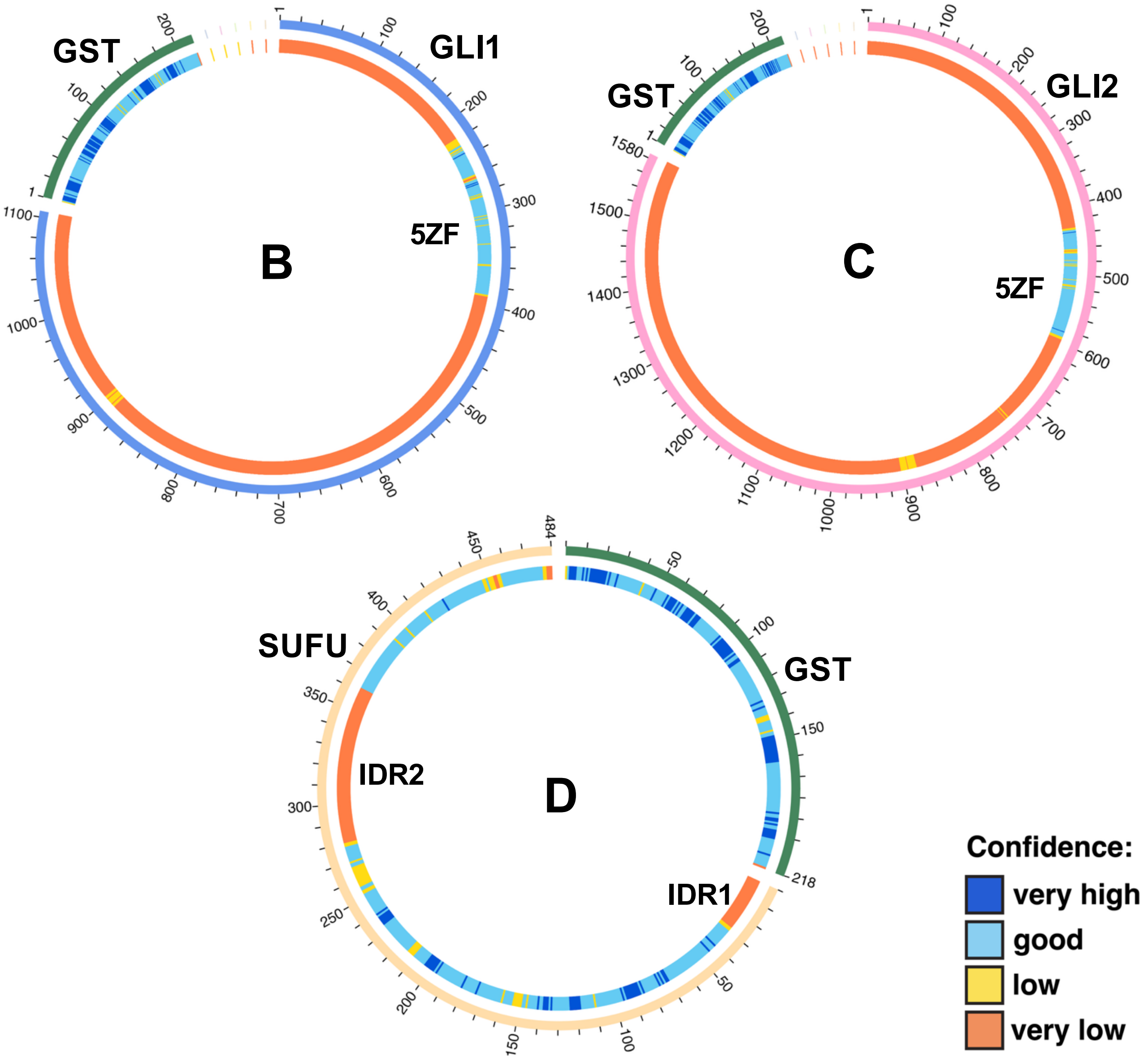


**Figure S12**. GST does not interact with GLI or SUFU in control co-foldings.

**A** Model confidence metrics for co-foldings of GST with GLI1, GLI2, SUFU or SUFU-D25. Average of top 3 models, each from a different job with a distinct random seed.
**B-D** Chord diagrams for representative models of GLI1 5z GST, GLI2 5z GST, and GST SUFU cofoldings. Lack of interaction is indicated by absence of chords connecting the proteins. **B** and **C** also show that SUFU-binding domains of GLI1 and GLI2 are predicted to be unstructured in the absence of SUFU.

Bardwell *et al*. 2026

Table S3

**GLI-SUFU contact residues in RSSL motif**

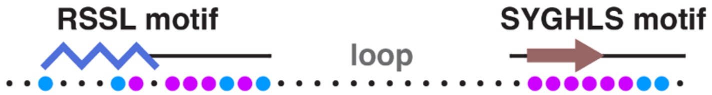


```
GLI1  93 LDLQTVIRTSPSSLVAFIN-SRCTSP-GGSYGHLSIGT 128
GLI2 243 LDLQRMIRTSPNSLVAYINNSRSSSAASGSYGHLSAGA 280
GLI3 304 FDLQTMIRTSPNSLVTILNNSRSSSSASGSYGHLSASA 341
  Ci 226 FDINSMIRFSPNSLATIMNGSRGSSAASGSYGHISATA 263
```

| | GLI contact residues | | | | SUFU contacts | |
|---|---|---|---|---|---|---|
| **Descriptor** | **GLI1** | **GLI2** | **GLI3** | **Ci** | **SUFU** | **dSufu** |
| Hydrophobic patch | Leu95 | Leu245 | Leu306 | Ile228 | Ala405<br>Trp430 | Ala380<br>Trp405 |
| | | Met248 | Met308 | Met231 | Trp430 | Trp405 |
| | Ile99 | Ile249 | Ile310 | Ile232 | *Ser267*<br>His394<br>Thr396 | *Ala263*<br>His371<br>Thr373 |
| Bidentate salt bridge | Arg100 | Arg250 | Arg311 | Arg233 | Cys156<br>***Asp159*** | Cys151<br>***Asp154*** |
| shielded H-bonds | Ser102 | Ser252 | Ser313 | Ser235 | **His394** | **His371** |
| | Pro103 | Pro253 | Pro314 | Pro236 | Arg393<br>**His394** | Arg370<br>**His371** |
| | Ser104 | Asn254 | Asn315 | Asn237 | Val413<br>**Glu414** | Val388<br>**Thr389** |
| | Ser105 | Ser255 | Ser316 | Ser238 | Val413<br>Glu414 | Val388<br>Thr389 |
| Hydrophobic knob | Leu106 | Leu256 | Leu317 | Leu239 | His394<br>Thr407<br>Val409<br>Val413<br>**Glu414**<br>Ala416<br>Trp430<br>Gln432 | His371<br>Thr382<br>Val384<br>Val388<br>**Thr389**<br>Ser391<br>Trp405<br>Gln407 |
| | Val107 | Val257 | Val318 | Ala240 | Gly415 | Gly390 |

**Bold** indicates that the SUFU residue participates in at least one hydrogen bond with the GLI residue. *Italics* indicates that the SUFU residue also contacts the GLI SYGHLS motif.

Grey, smaller text indicates a contact not fully supported by all the evidence:

- GLI2/GLI3 Met248/308 with SUFU Trp430 does not meet confidence thresholds
- GLI Leu106/256/317 with SUFU Ala416 and Trp430 are not ChimeraX contacts.

Bardwell *et al*. 2026
Table S4

**GLI-SUFU contact residues in SYGHLS motif**

SYGHLS motif

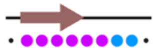

```
GLI1 119 GSYGHLSIGT 128
GLI2 271 GSYGHLSAGA 280
GLI3 332 GSYGHLSASA 341
  Ci 254 GSYGHISATA 263
```

| | **GLI contact residues** | | | | **SUFU contacts** | |
|---|---|---|---|---|---|---|
| **Descriptor** | **GLI1** | **GLI2** | **GLI3** | **Ci** | **SUFU** | **dSufu** |
| β-strand cap | Gly119 | Gly271 | Gly332 | Gly254 | -- | |
| | Ser120 | Ser272 | Ser333 | Ser255 | Gly158<br>**Asn265**<br>Leu266<br>***Ser267*** | Gly153<br>**Asp261**<br>Leu262<br>***Ala263*** |
| β-strand core | Tyr121 | Tyr273 | Tyr334 | Tyr256 | ***Asp159***<br>**His160**<br>Leu266<br>Gly268<br>Val269<br>**Ser270**<br>Lys398 | ***Asp154***<br>**Asn155**<br>Leu262<br>Gly264<br>Val265<br>**Asn266**<br>Lys375 |
| | Gly122 | Gly274 | Gly335 | Gly257 | His160<br>Leu266<br>**Gly268**<br>Val269<br>**Ser270** | Asn155<br>Leu262<br>**Gly264**<br>Val265<br>**Asn266** |
| | His123 | His275 | His336 | His258 | **Tyr147**<br>Phe155<br>**Asp159**<br>**His160**<br>Val161<br>**Ser162** | **Tyr142**<br>Leu150<br>**Asp154**<br>**Asn155**<br>Ile156<br>Pro157 |
| β-strand terminators | Leu124 | Leu276 | Leu337 | Ile259 | His160<br>Ser162<br>Gln212<br>Val269<br>**Ser270**<br>Ala271<br>Glu376<br>--<br>Leu380<br>Ile400 | --<br>--<br>--<br>Val265<br>**Asn266**<br>--<br>--<br>Val354<br>Tyr357<br>-- |
| | Ser125 | Ser277 | Ser338 | Ser260 | **Ser162**<br>**Glu376** | --<br>-- |
| | Ile126 | Ala278 | Ala339 | Ala261 | Tyr147<br>Val161 | Tyr142<br>Ile156 |
| | Gly127 | Gly279 | Ser340 | Thr262 | Trp163<br>-- | Trp158<br>Lys160 |

**Bold** indicates that the SUFU residue participates in at least one hydrogen bond with the GLI residue. *Italics* indicates that the SUFU residue also contacts the GLI RSSL motif.

Bardwell *et al*. 2026
Table S5

**GLI-SUFU contact residues in SIC domain**

**SIC domain**

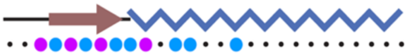

```
GLI1 1080  PPNMAVGNMSVLLRSLPGETEFLNSSA  1106
GLI2 1560  ISNMAVGDMSSMLTSLAEESKFLNMMT  1586
GLI3 1554  TTNMAIGDMSSLLTSLAEESKFLAVMQ  1580
  Ci 1371  GTNMVINDMTTSLTSLLEENRYLQMMQ  1397
```

| | **GLI contact residues** | | | | **SUFU contacts** | |
|---|---|---|---|---|---|---|
| **Descriptor** | **GLI1** | **GLI2** | **GLI3** | **Ci** | **SUFU** | **dSufu** |
| H-bond hub | Asn1082 | Asn1562 | Asn1556 | Asn1373 | Ala56<br>**Ile57**<br>**Val58**<br>Pro65<br>**Asp66** | Thr40<br>**Leu41**<br>**Leu42**<br>--<br>**Asp50** |
| β-strand cap | Met1083 | Met1563 | Met1557 | Met1374 | *Val54*<br>Thr55<br>Gln142 | Val38<br>Thr39<br>-- |
| β-strand | Ala1084 | Ala1564 | Ala1558 | Val1375 | *Val54*<br>**Thr55**<br>Ile57 | Val38<br>**Thr39**<br>Leu41 |
| | Val1085 | Val1565 | Ile1559 | Ile1376 | Gln53<br>Thr135 | Gln37<br>Thr130 |
| | Gly1086 | Gly1566 | Gly1560 | Asn1377 | *Leu52*<br>**Gln53** | Leu36<br>**Gln37** |
| β-strand terminator | Asn1087 | Asp1567 | Asp1561 | Asp1378 | Asn50<br>Pro51 | Asn34<br>Pro35 |
| Hydrophobic knob 1 | Met1088 | Met1568 | Met1562 | Met1379 | Tyr38<br>--<br>Asn50<br>Pro51<br>Gln53 | Ile22<br>Pro33<br>Asn34<br>Pro35<br>Gln37 |
| | Ser1089 | Ser1569 | Ser1563 | Thr1380 | **Asn50** | **Asn34** |
| Knob2 | Leu1091 | Met1571 | Leu1565 | Ser1382 | Gln53<br>--<br>Tyr70 | Gln37<br>Thr39<br>Tyr54 |
| Knob2 | Leu1092 | Leu1572 | Leu1566 | Leu1383 | Phe30<br>-- | --<br>Ile22 |
| Knob3 | Leu1095 | Leu1575 | Leu1569 | Leu1386 | Pro31<br>--<br>-- | Pro15<br>Tyr54<br>His83 |

**Bold** indicates that the SUFU residue participates in at least one hydrogen bond with the GLI residue.
*Italics* indicates that the SUFU residue also contacts the SR motif in GLI2 and GLI3.

Grey, smaller text indicates a contact not fully supported by all the evidence:

- GLI2/GLI3 Ser1569/1563 have no contacts in full-length structures 92# and 248#

Bardwell *et al*. 2026
Table S6

### GLI-SUFU contact residues in SR motif

```
                       SR motif
      GLI2  61   PLPIDMRHQEGRYHY  75
      GLI3 134   PVPIDARHHEGRYHY  148
SUFU contacts    •••●•••●●•●•●•●
 SIC contacts    ●●•●•••●•••••••
```

● contact does not involve H-bonds
● contact includes at least one H-bond

| SR Residue | | Target Residue | | |
|---|---|---|---|---|
| **GLI2** | **GLI3** | **SUFU** | **GLI2** | **GLI3** |
| Pro61 | Pro134 | | Ala1564<br>Val1565 | Ala1558<br>Ile1559 |
| Leu62 | Val135 | | Ala1564<br>**Val1565** | Ala1558<br>**Ile1559** |
| Ile64 | Ile137 | *Val54*<br>*Thr135* | Ala1564 | Ala1558 |
| His68 | His141 | Pro134<br>*Thr135* | Val1565 | Ile1559 |
| Gln69 | His142 | Pro134<br>*Thr135*<br>Trp136 | | |
| Gly71 | Gly144 | Pro134 | | |
| Tyr73 | Tyr146 | **Ser131**<br>Ala132<br>Pro133 | | |
| Tyr75 | Tyr148 | *Leu52* | | |

**Bold** indicates that the target residue participates in at least one hydrogen bond with the SR residue. *Italics* indicates that the SUFU residue also contacts the GLI SIC domain.

Only GLI-GLI contacts between the SR motif and the SIC domain are shown; intradomain contacts are not shown.

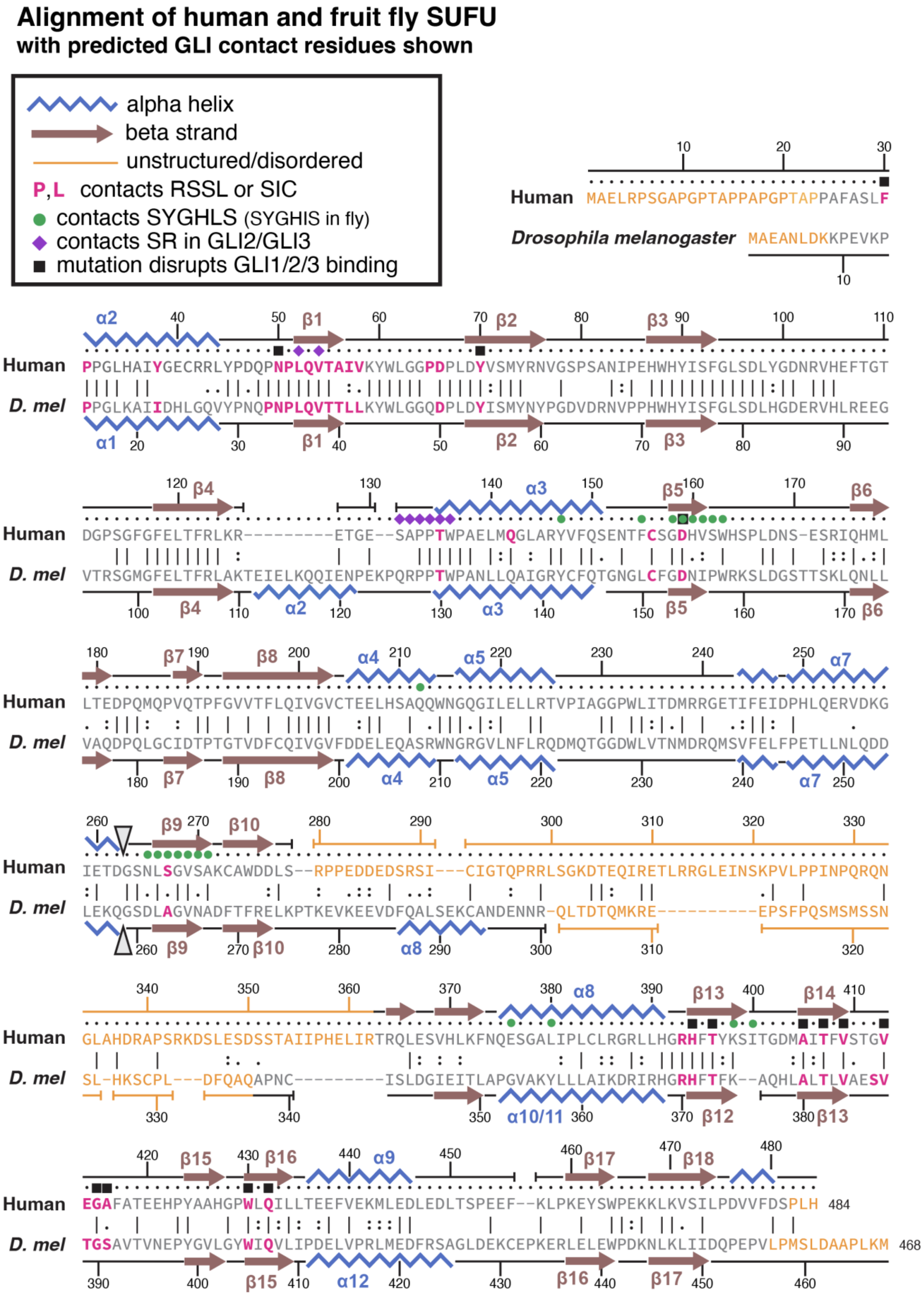


**Figure S13**. Alignment of human and fly SUFU with GLI contact residues shown.

**Figure S13** (previous page). Alignment of human SUFU with fruit fly (*D. melanogaster*) dSufu.

A key to aid interpretation is shown at the top.

Pink bold letters = residues that make high-quality contacts with the GLI/Ci SIC domain or RSSL motif (Tables S3, S5).

SIC-domain-contacting residues are found in the first half of the N-terminal lobe of SUFU, before the end of $\alpha$3. The first SIC-domain-contacting residue in human SUFU is F30 and the last is Q142.

RSSL-motif-contacting residues start after the end of $\alpha$3. The first RSSL-motif-contacting residue in human SUFU is C156 and the last is Q432.

Green dots = residues that contact the GLI/Ci SYGHLS motif. All the green dots are SYGHLS contacts for both GLIs and Ci (see Table S4), except E376 which is not found in Ci.

Purple diamonds = Residues that contact the GLI2 and GLI3 SR motifs (Table S6).

Black squares indicate SUFU residues whose mutation disrupted binding of the GLI SIC domain (F30, N50, Y70) or NR domain (all other black squares) in the experiments performed in Figures 6, 9, 10 and S23.

The triangles at the end of $\alpha$7 indicate the transition from the SUFU N-terminal domain to the C-terminal domain.

Helix and sheet numbering is from Zhang *et al*. 2013 Figure S1. Their $\alpha$1 helix (residues 27-29) is not shown.

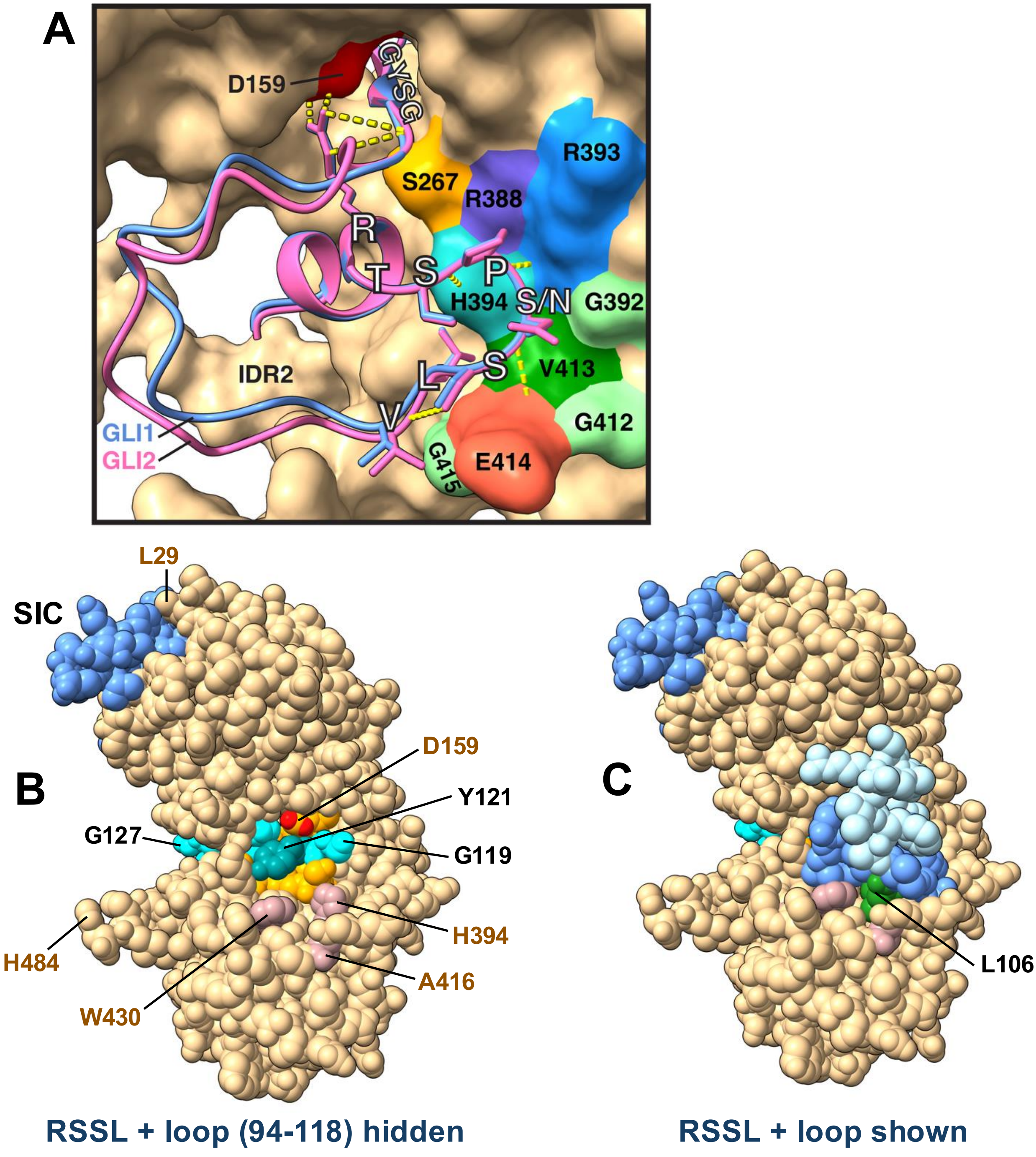


**Figure S14**. GLI1 and GLI2 RSSL motif H-bonds; burying of SYGHLS by RSSL.

**A** Interactions of the SP(S/N)S residues of the RSSL motif. Both GLI1 (CSM 70#) and GLI2 (CSM 155#) are shown after their SUFU chains were superimposed. SUFU (from CSM 70#) is shown with its surface colored tan and select contact residues in other colors. The orientation is similar to Figure 4C.

**B** GLI1 CSM 261# is shown in a similar orientation to Figure 1A but in sphere representation. The SYGHLS motif is colored cyan; the SIC domain is colored cornflower blue. β5 and β9 of SUFU are colored orange. The RSSL motif (94-107) and the 108-118 loop are hidden. Other residues are indicated with brown text for SUFU residues and black text for GLI residues.

**C** Same as **B** but the RSSL motif is now shown colored cornflower blue (with L106 green) and the 108-118 loop is colored light blue.

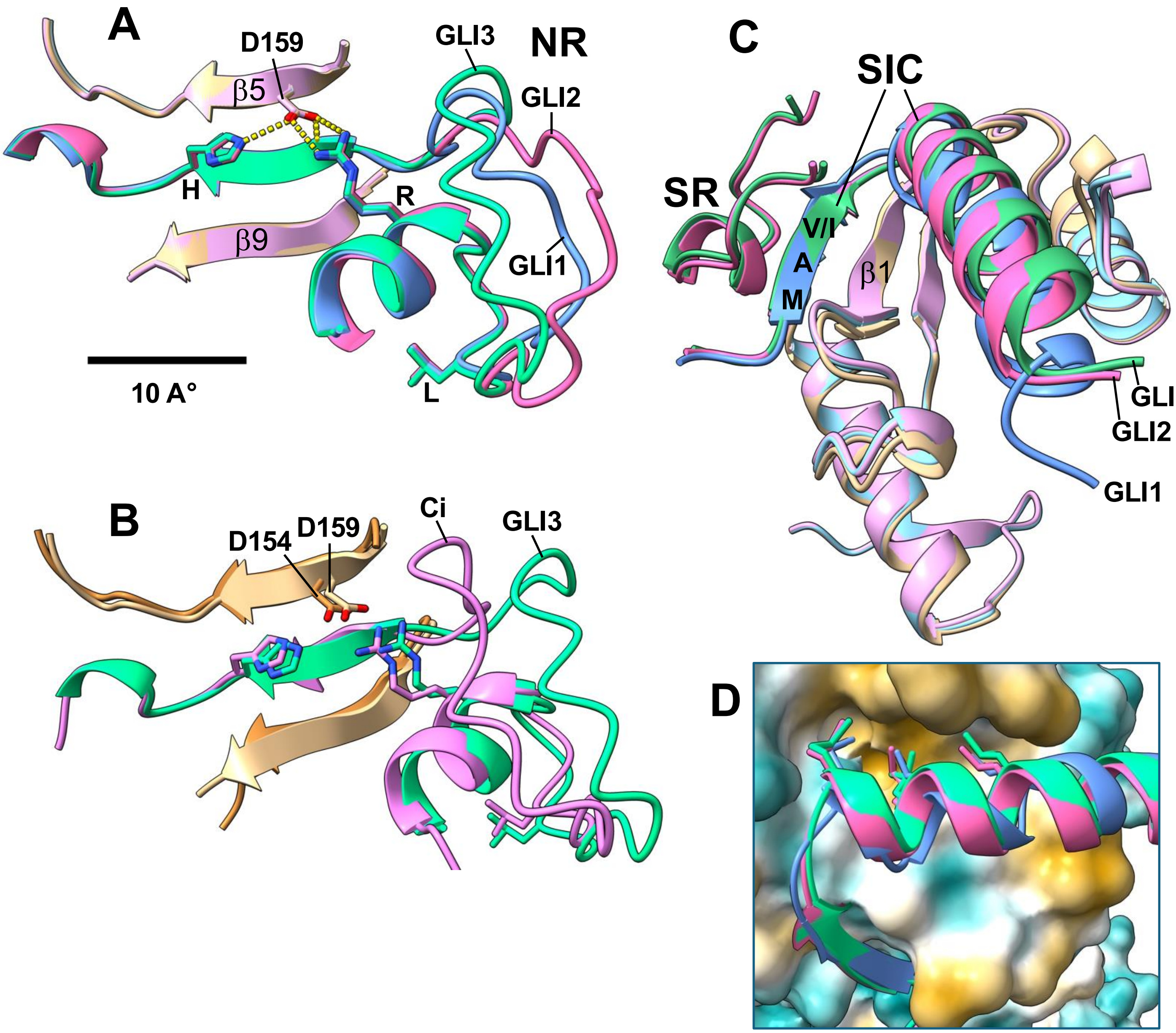


**Figure S15**. Comparison of GLI1, GLI2 and GLI3 CSMs.

**A** NR domains of GLI1/2/3. CSM 111# (GLI1 5z SUFU), 92# (GLI2 5z SUFU) and 248# (GLI3 5z SUFU) were superimposed with respect to the SUFU structures. SUFU β5 and β9 and the side chain of SUFU D159 are shown (pastel colors) and the H-bonds involving D159's side chain (yellow dashes). The sidechains of GLI1/GLI2/GLI3 R100/R250/R311, L106/L256/L317 and H123/H275/H336 are also shown. The GLI structures superimpose with RMSD <1, except in the unstructured loop that connects the RSSL and SYGHLS motifs. Scalebar = 10 A° (applies to **A**-**C**).

**B** NR domains of GLI3/Ci. Same as **A** except comparing CSM 228# (Ci NR & SIC, dSufu) and 248# (GLI3 5z SUFU). The sidechains of Ci/GLI3 R233/R311, L239/L317 and H258/H336 are shown.

**C** SIC domains of GLI1/2/3 and SR motifs of GLI 2/3. Same CSMs as **A**. SUFU residues 29-76 and 134-163 are shown (pastel colors).

**D** SIC domains of GLI1/2/3. Same CSMs as **C** but different orientation, with SUFU surface colored by hydrophobicity. GLI side chains shown left-to-right are M1088/M1568/M1562, L1092/L1572/L1566 and L1095/L1575/L1569.

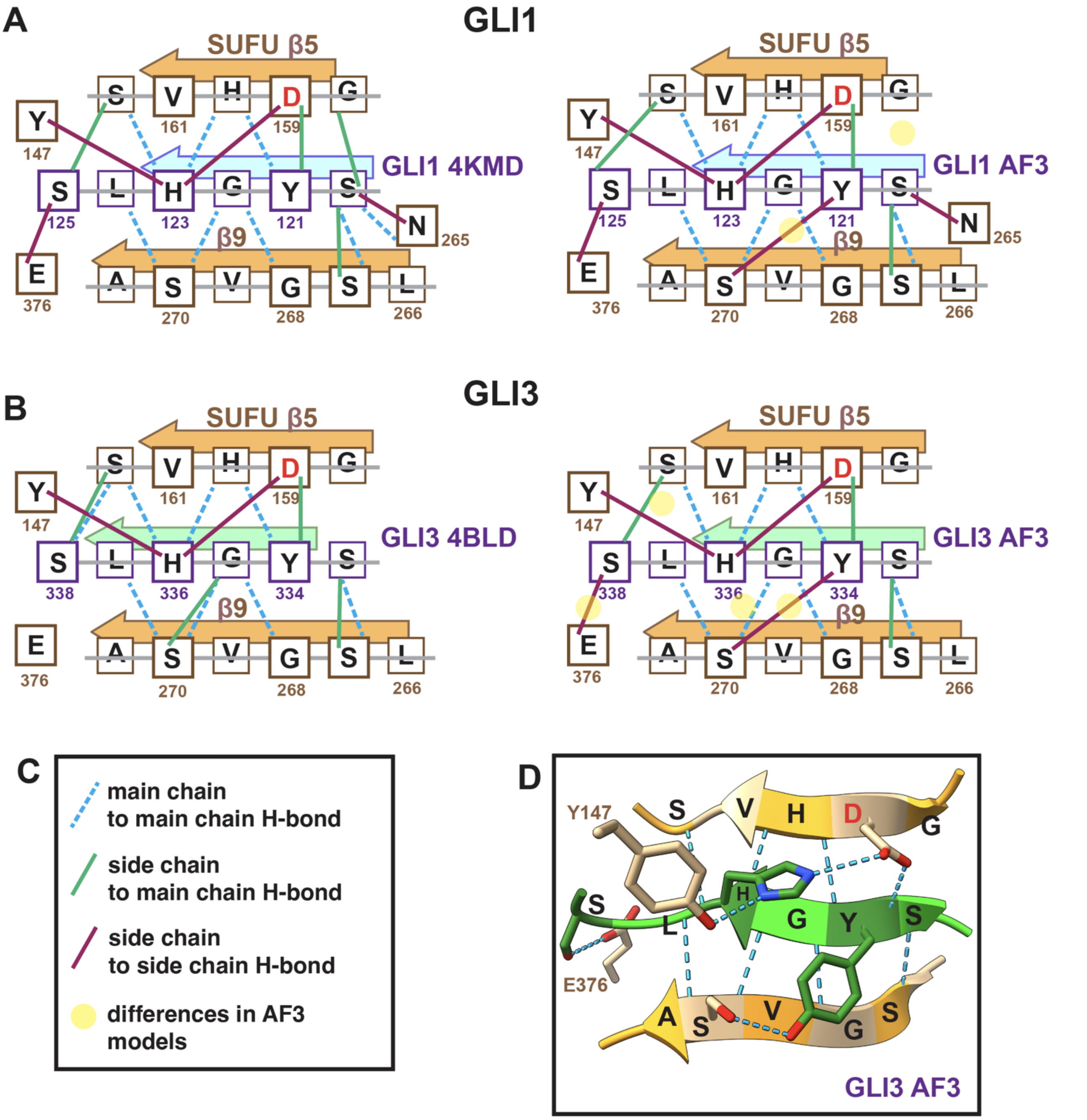


**Figure S16**. SYGHLS motif H-bonds for GLI1 and GLI3, empirical vs. AF3-predicted.

**A** Left: Hydrogen bonding pattern of GLI1 SYGHLS motif seen in PDB model 4KMD.
Right: Consensus of AF3-predicted H-bonds in GLI1 CSMs. Differences from 4KMD are highlighted with yellow circles.

**B** Left: Hydrogen bonding pattern of GLI3 SYGHLS motif seen in PDB model 4BLD.
Right: Consensus of AF3-predicted H-bonds in GLI3 CSM's. Differences from 4BLD are highlighted with yellow circles.

**C** Key for **A** and **B**.

**D** Example from GLI3 CSM 248#. All H-bonds shown as blue dashes.

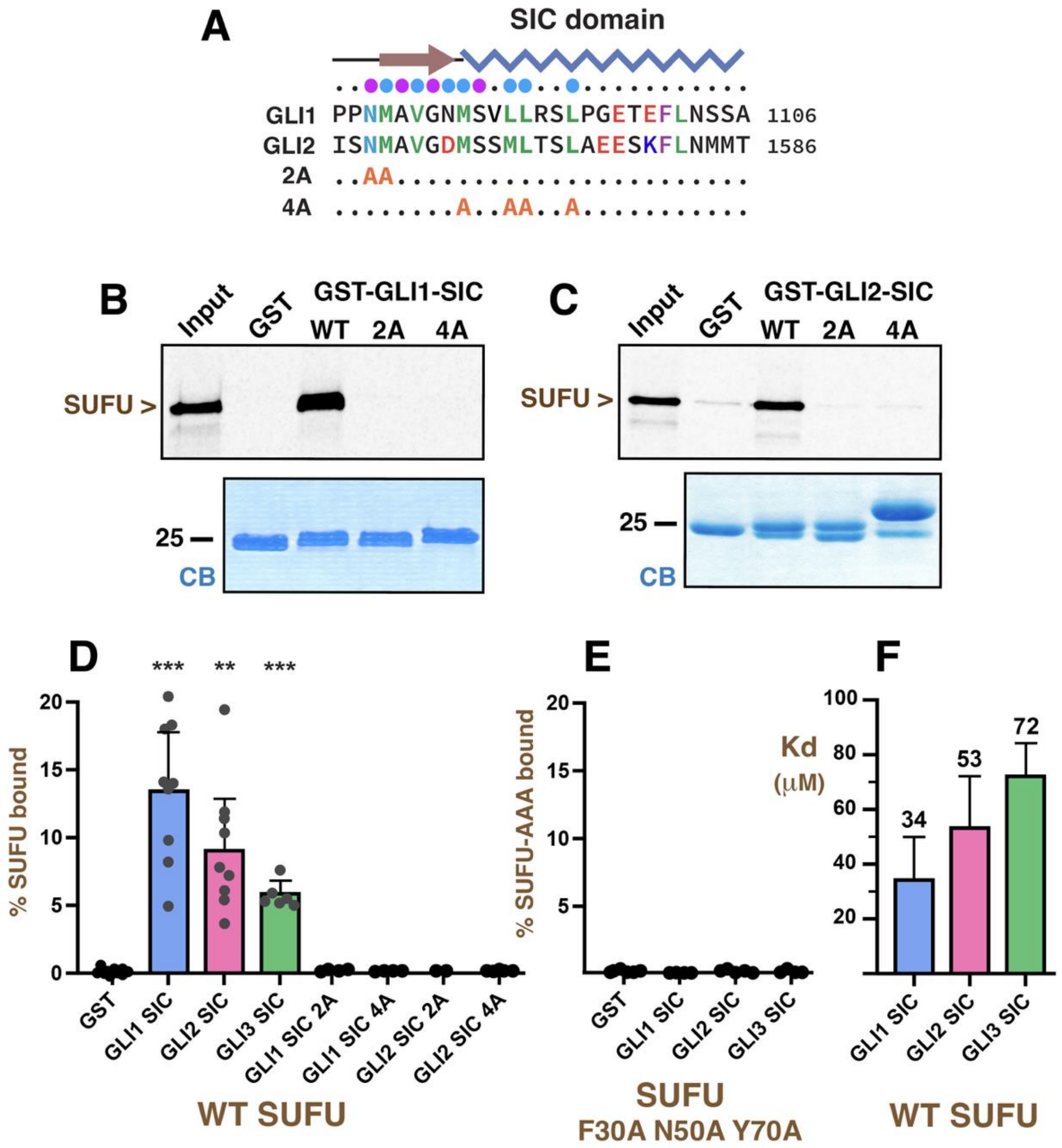


**Figure S17**. Substitutions of SIC domain contact residues with alanine abolish SIC-SUFU binding

**A** Sequence of the SIC domains of GLI1 and GLI2 with residues substituted to alanine in the 2A and 4A mutants shown.

**B** $^{35}$S-radiolabeled full-length wild-type human SUFU protein was tested for binding to purified GST-GLI1-SIC (WT) or the 2A or 4A variants thereof. Other details as in Figure 6E.

**C** Same as B, but using GST-GLI2-SIC and the 2A or 4A variants thereof.

**D, E** Quantification of binding assay results, as shown in Figure 6E, but with additional data added for the 2A and 4A mutants (**D**) and for the binding of the SUFU F30A N50A Y70A variant (**E**). 95% confidence intervals are shown; individual data points are displayed as dots. Binding of SUFU to all three SIC domains was significantly different from the minimal binding to GST alone: **, $p < 0.01$, ***, $p < 0.001$. Binding of SUFU to the SIC domain mutants was not significantly different than binding to GST alone. Binding of the SUFU F30A N50A Y70A to the GLI1, GLI2 and GLI3 SIC domains was also not significantly different than binding to GST alone.

**F** Data points in **D** were converted to a Kd as described in Materials and Methods, n = 6-9. Numbers above each bar are the mean Kd in micromolar units; error bars show the 95% confidence interval.

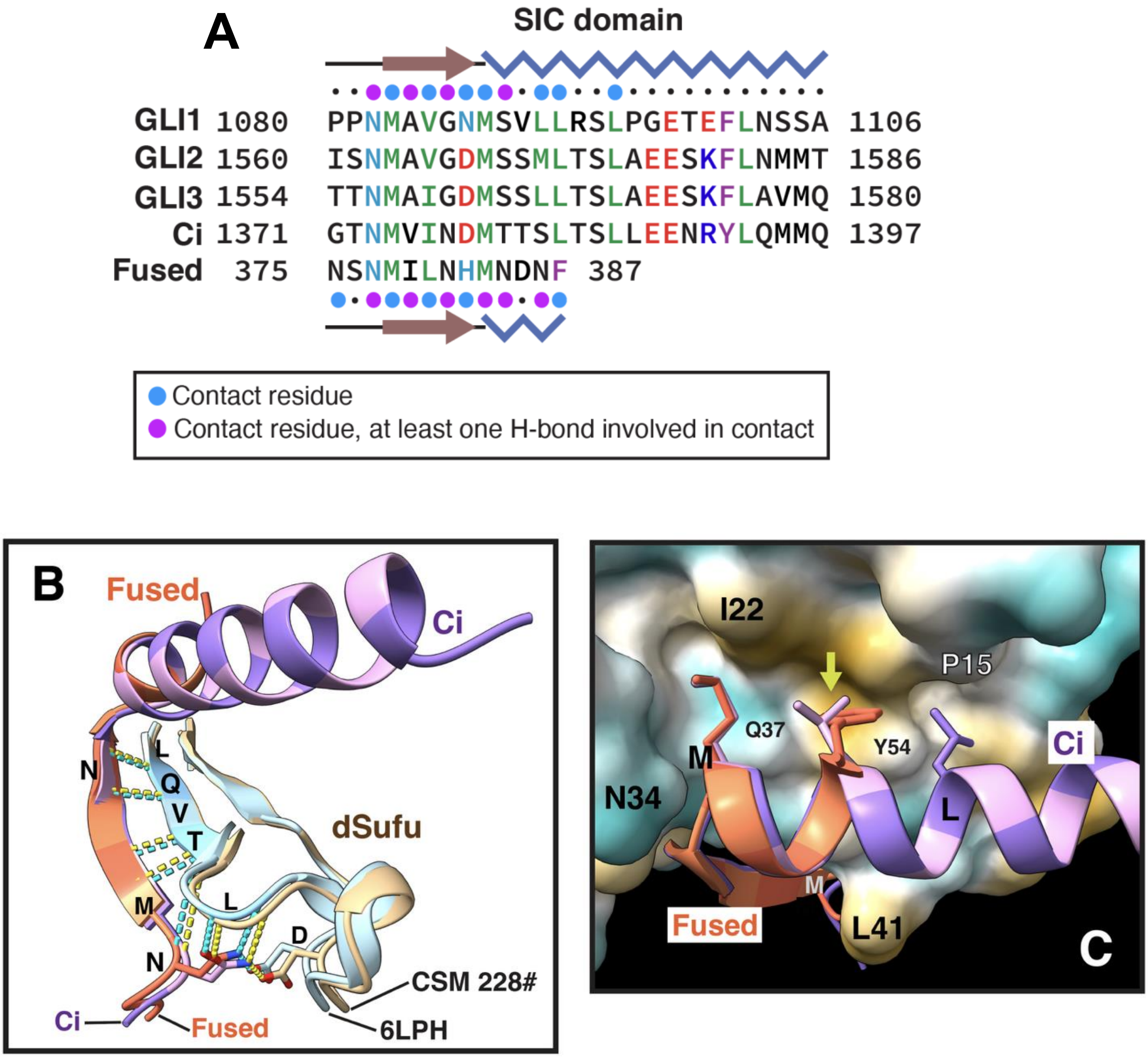


**Figure S18**. Ci and Fused bind to the same elements of dSufu.

**A** Alignment of SIC domains of human GLI1, GLI2 and GLI3 with D. melanogaster Ci and the Sufu-binding site (SBS) of *D. melanogaster* Fused. Residue coloring as in Figure 4A and S9A. Secondary structure and contact residues for interface regions is shown above (for GLI/Ci) and below (for Fused). Blue zig-zags depict helix and brown arrows depict β-strands. Cyan and magenta-colored circles denote residues that contact SUFU/dSufu, with magenta indicating the involvement of at least one H-bond in the contact (key shown).

**B** Superposition of Ci SIC domain (purple) bound to dSufu (tan) from CSM 228# with the Fused SBS (reddish orange) bound to the N-terminal domain of dSufu (light blue) from PDB structure 6LPH. SBS H-bonds are cyan, Ci H-bonds are yellow.

**C** Interaction of Ci and Fused SBS with hydrophobic groove on dSufu. dSufu hydrophobicity shown as in Figure 6C. Note the near-perfect superposition of Ci M1379 and Fused M383, and how Ci L1383 and Fused F387 fill the same shallow pocket on dSufu (arrow).

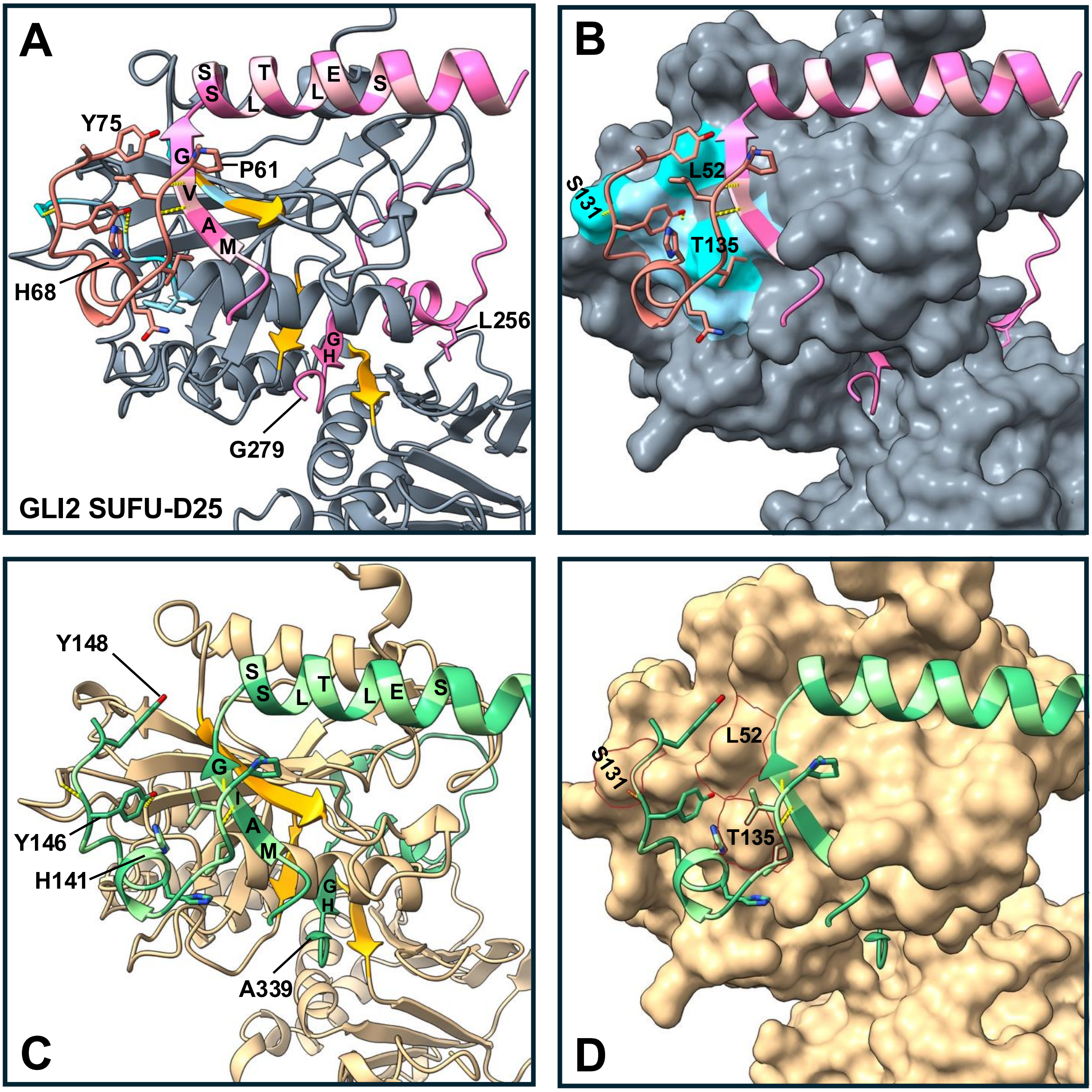


**Figure S19**. SR motif structure and interactions.

**A** GLI2's SR motif (residues 61-75) is colored rose; the C-terminal portion of the NR domain (245-279; includes both the RSSL and SYGHLS motifs) and SIC domain (1561-1586) of GLI2 are colored shades of pink. SUFU-D25 is colored slate gray with β-strands 1, 5 and 9 colored orange. SUFU residues that contact the SR motif are shaded sky blue or aqua. H-bonds made by GLI2 L62 and Y73 are shown as yellow dashes. The residue numbers of some important contact residues in GLI2 are shown. Model is CSM 294# GLI2 5z SUFU-D25.

**B** Same as **A** but with SUFU displayed in surface representation and orange coloring changed to slate gray. The numbers of some SUFU residues that contact the GLI SR motif are shown.

**C, D** Similar to **A** and **B**, but model is GLI3 5z SUFU-D25 CSM 190#. GLI3 is colored shades of green and SUFU-D25 is colored tan.

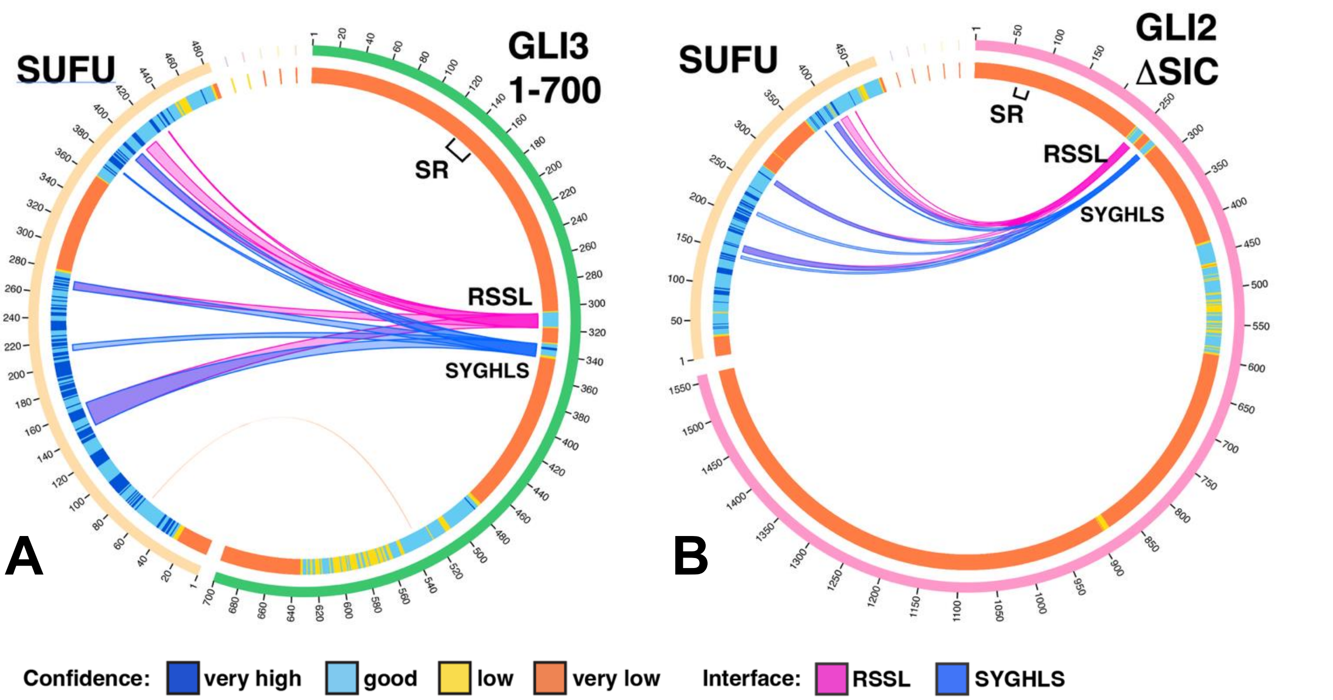


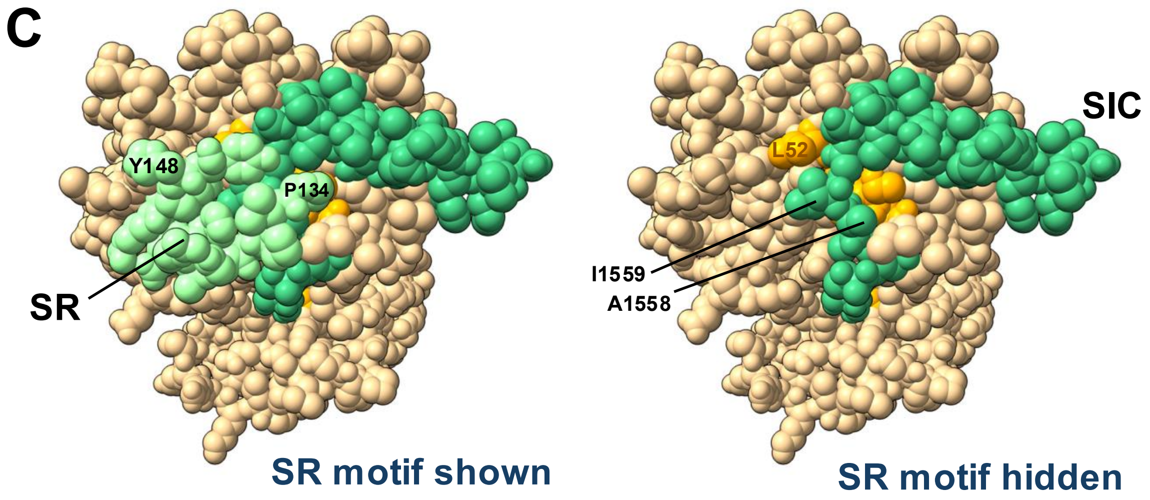


**Figure S20**. The SR motif is unstructured, and does not contact SUFU, in co-foldings lacking the SIC domain.

**A** Lack of structured SR motif in GLI3 repressor fragment (residues 1-700).
Model is CSM 280#, GLI3 1-700 5z SUFU

**B** Lack of structured SR motif when SIC domain (residues 1561-1586) is deleted from GLI2.
Model is CSM 374#, GLI2 1-1560 5z SUFU. Very similar results were obtained with GLI3 1-1554 5z SUFU (not shown).

**C** The SR motif contacts the SIC domain β-strand and shields it from solvent. Top down view of GLI3 CSM 248# with SR motif shown (left) or hidden (right). SUFU is tan with β1 orange; GLI3 SR motif is pale green and the SIC domain is darker green.

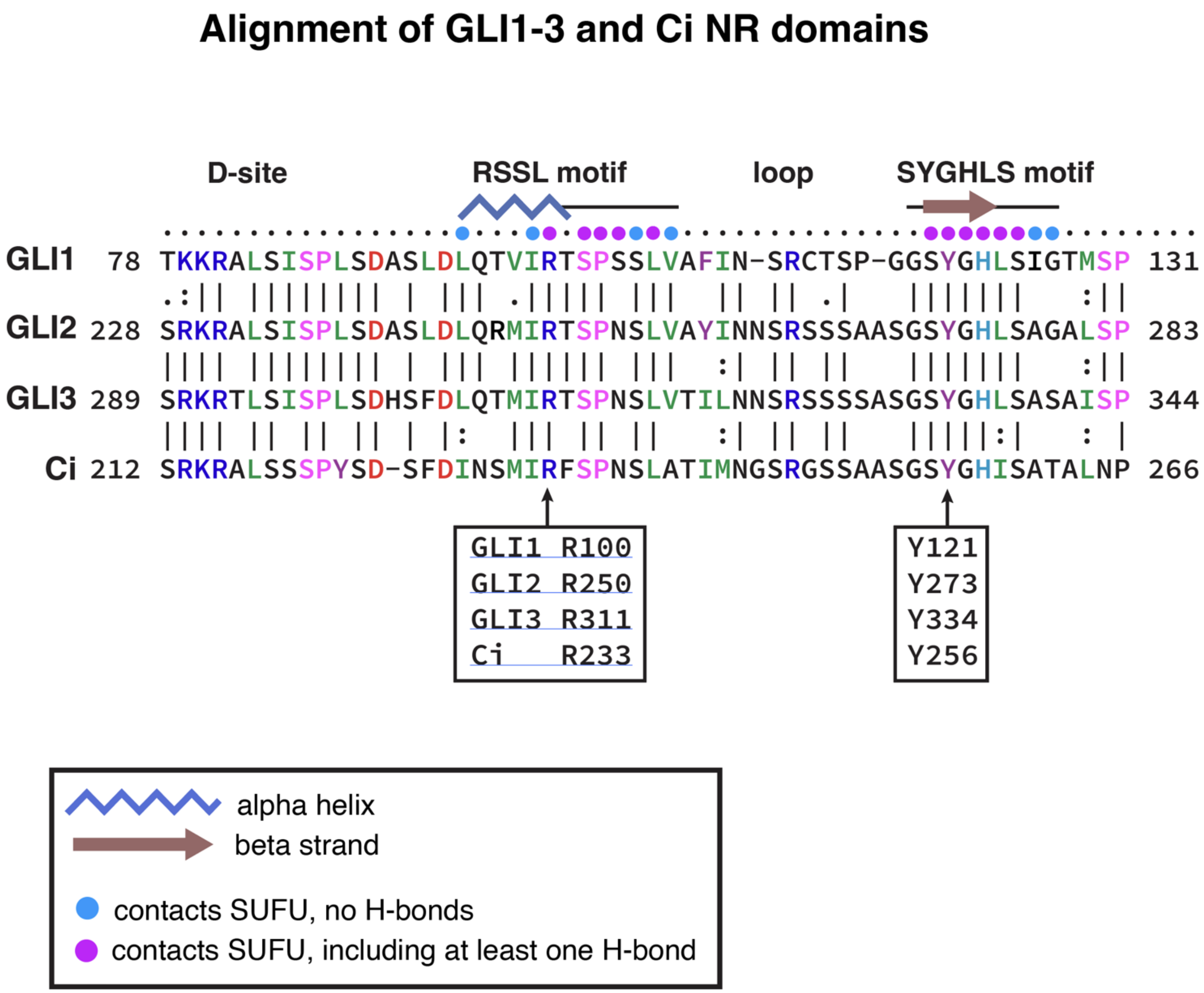


**Figure S21**. Alignment of the NR domains of GLI1, GLI2, GLI3 and Ci.

Vertical dashes indicate amino acid identity; dots indicate amino acid conservation (BLOSUM62). Only residues identical or conserved between all 3 human GLI proteins are so indicated; for these residues, conservation with Ci is also shown.
Residue coloring as in Figures 4 and S9A, with the addition that known phosphorylation sites for proline directed kinases are colored pink. The secondary structure for interface regions is shown above, with blue zig-zags for $\alpha$-helix and brown arrows for $\beta$-strands. 'D-site' indicates the MAP kinase docking site (Whisenant et al. 2010).

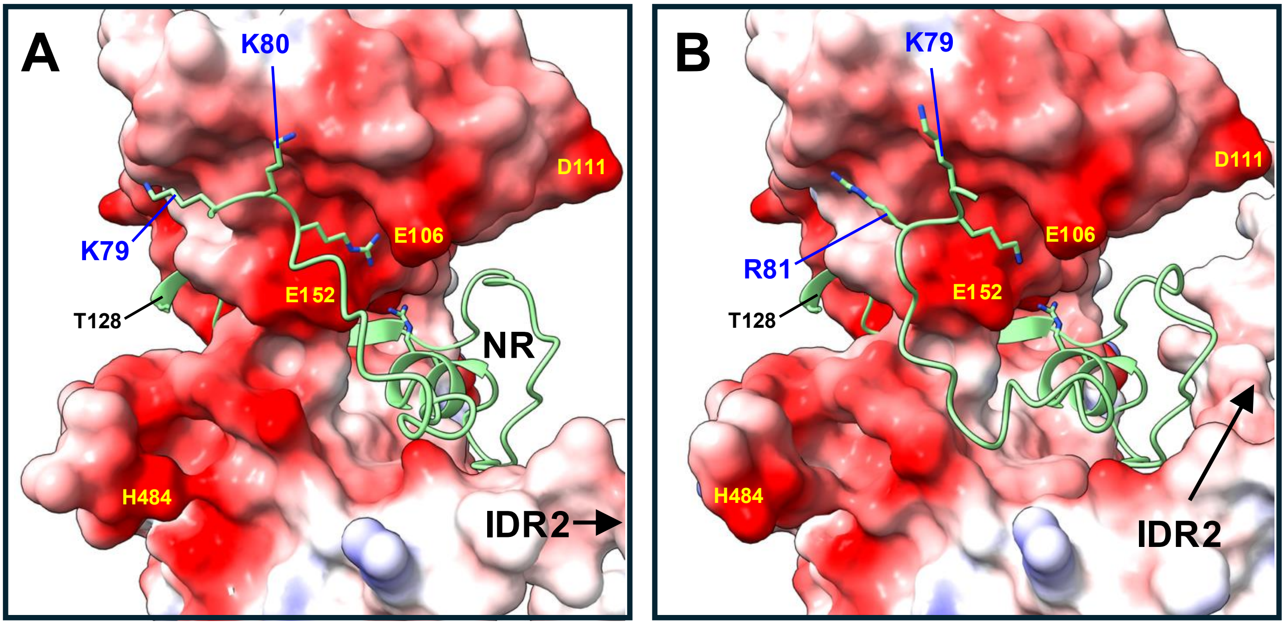


**Figure S22**. Possible fuzzy electrostatic interaction of GLI1 K79, K80, R81 with SUFU.

**A** Residues 79-128 of the GLI1 NR domain are shown in light green; SUFU is shown with electrostatic surfaces, red = negative, blue = positive, white = neutral. Model is CSM 25#, GLI1 1-232 SUFU.

**B** Same as **A**, but model is CSM 261#, GLI1 5z SUFU.

Note the different configurations of the positively-charged GLI1 K79, K80, R81 in the two models, even though collectively they contact similar negatively-charged regions of SUFU.

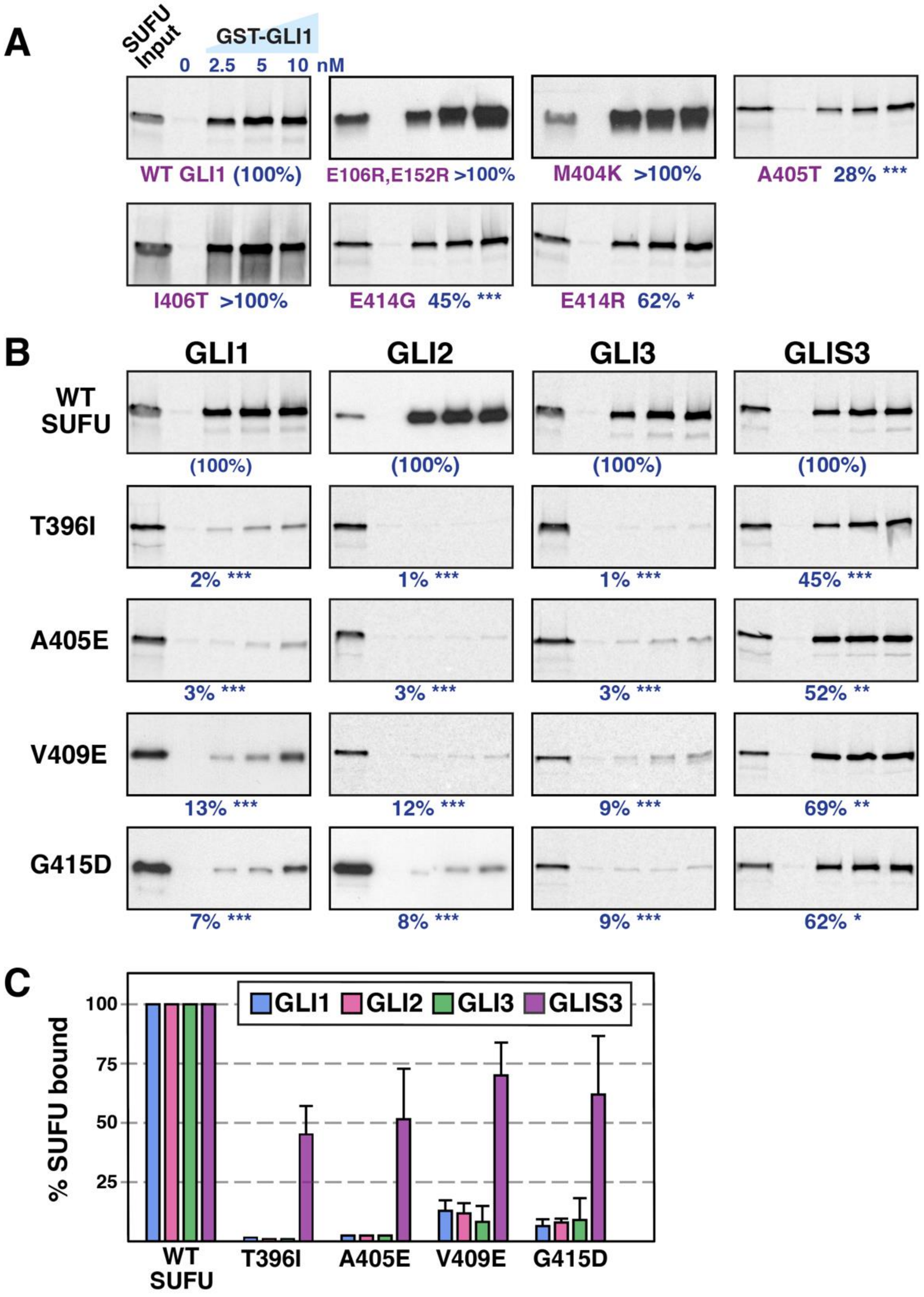


**Figure S23**. Additional SUFU mutations assessed for GLI and GLIS3 binding

**A** Radiolabeled SUFU variants were assessed for binding to GST-GLI1$_{68\text{-}232}$. Representative gel analysis results of 2-4 independent experiments are shown. Other details as in Figure 9.

**B** Radiolabeled SUFU variants were assessed for binding to GST-GLI proteins or GST-GLIS3. Representative gel analysis results of 2-4 independent experiments are shown. Other details as in Fig. 10.

**C** Data from the 2.5, 5 and 10 nM GST-GLI/GLIS points from all replicate experiments were averaged to determine percent binding and normalized to wild-type, n = 6-12. Error bars show the 95% confidence interval.

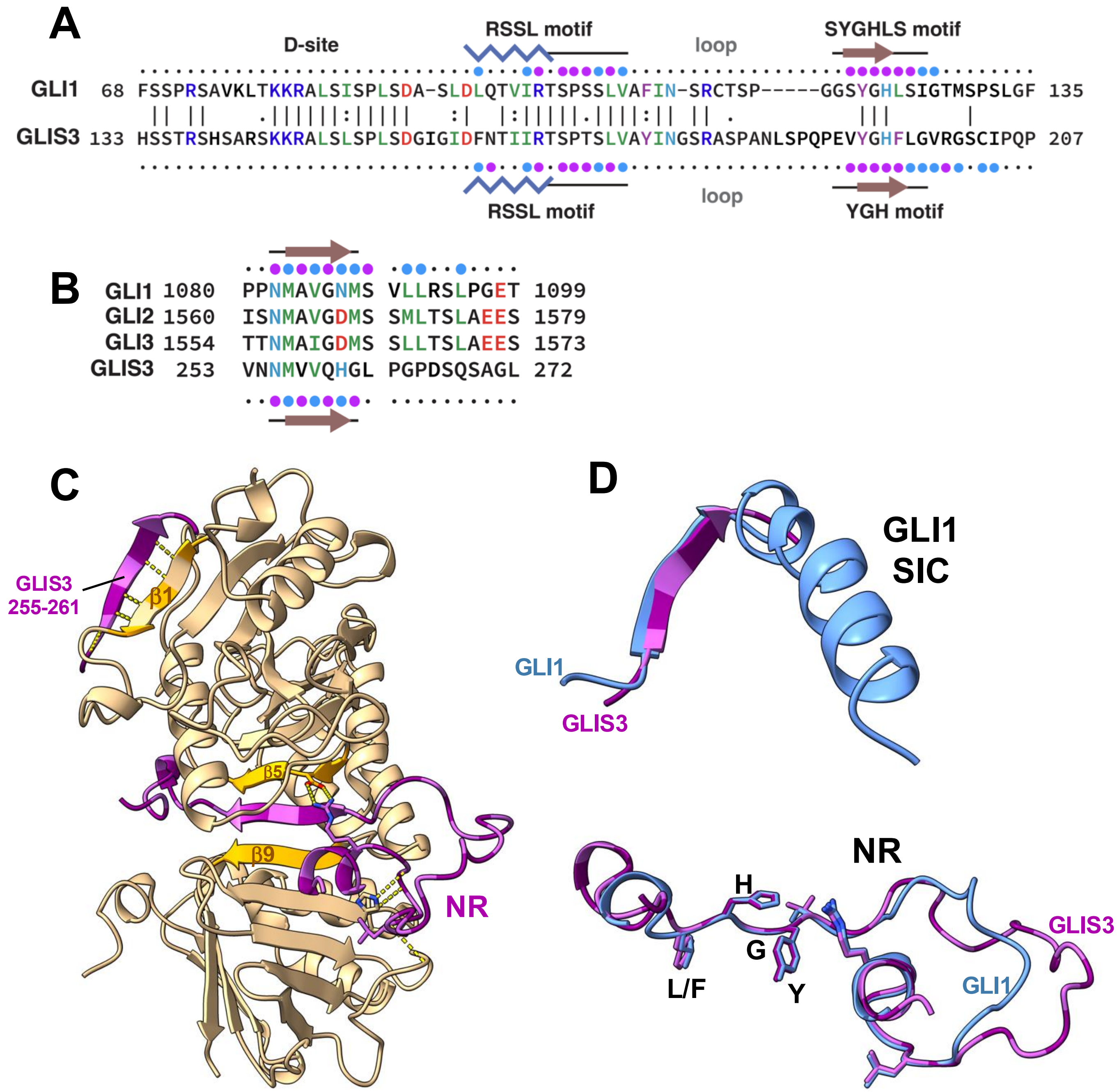


**Figure S24**. Similarity of GLIS3 and GLI1 interfaces with SUFU

The predicted GLIS3-SUFU structure displays RSSL- and YGH-mediated contacts with SUFU that are very similar to those with the GLI proteins, but reveals two additional contact regions that may explain the increased relative affinity of GLIS3 for SUFU mutants that otherwise disrupt GLI binding (Figure 10, Figure S23). First, GLIS3 makes additional contacts with SUFU using residues immediately C-terminal to its YGH motif; similar contacts are not seen in the predicted GLI-SUFU structures. Second, GLIS3 residues 255-261, which reside between the NR domain and the 5ZF domain, form an antiparallel β-strand that hydrogen bonds with SUFU β1 using a β-strand augmentation mechanism, echoing how the β-strand of the GLI SIC domain binds to SUFU β1 (and how the Fused SBS binds to dSufu β1). Unlike these other structures, however, the predicted GLIS3-SUFU structure does not show the presence of a subsequent helix that makes additional contacts with SUFU.

**Figure S24** (continued from previous page)

**A** Alignment of NR domains of human GLI1 and GLIS3. Contact residues and secondary structure indicated as in previous figures, with GLI contact residues shown on the top line and GLIS3 contact residues on the bottom line. Residue coloring as in previous figures.

**B** Alignment of SIC domains of human GLI1/2/3 with residues 253-272 of GLIS3, focusing on β-strand.

**C** CSM 327# GLIS3 105-284 SUFU. The structure shows full-length SUFU in tan with IDR1 and IDR2 hidden. SUFU β1, β5 and β9 are colored orange and the side chains of D159 and H394 are shown. GLIS3 160-205 and 255-261 are shown in shades of maroon with the side chains of R166 and L172 shown. Select hydrogen bonds are shown as yellow dashes.

**D** GLI1/GLIS3 comparison after superposition of SUFU chains from GLIS3 105-284 SUFU 327# and GLI1 5z SUFU 261#. SUFU chains are hidden, side chains of S/V-YGH-L/F motif residues are shown in addition to those of R100/R166 and L106/L172.

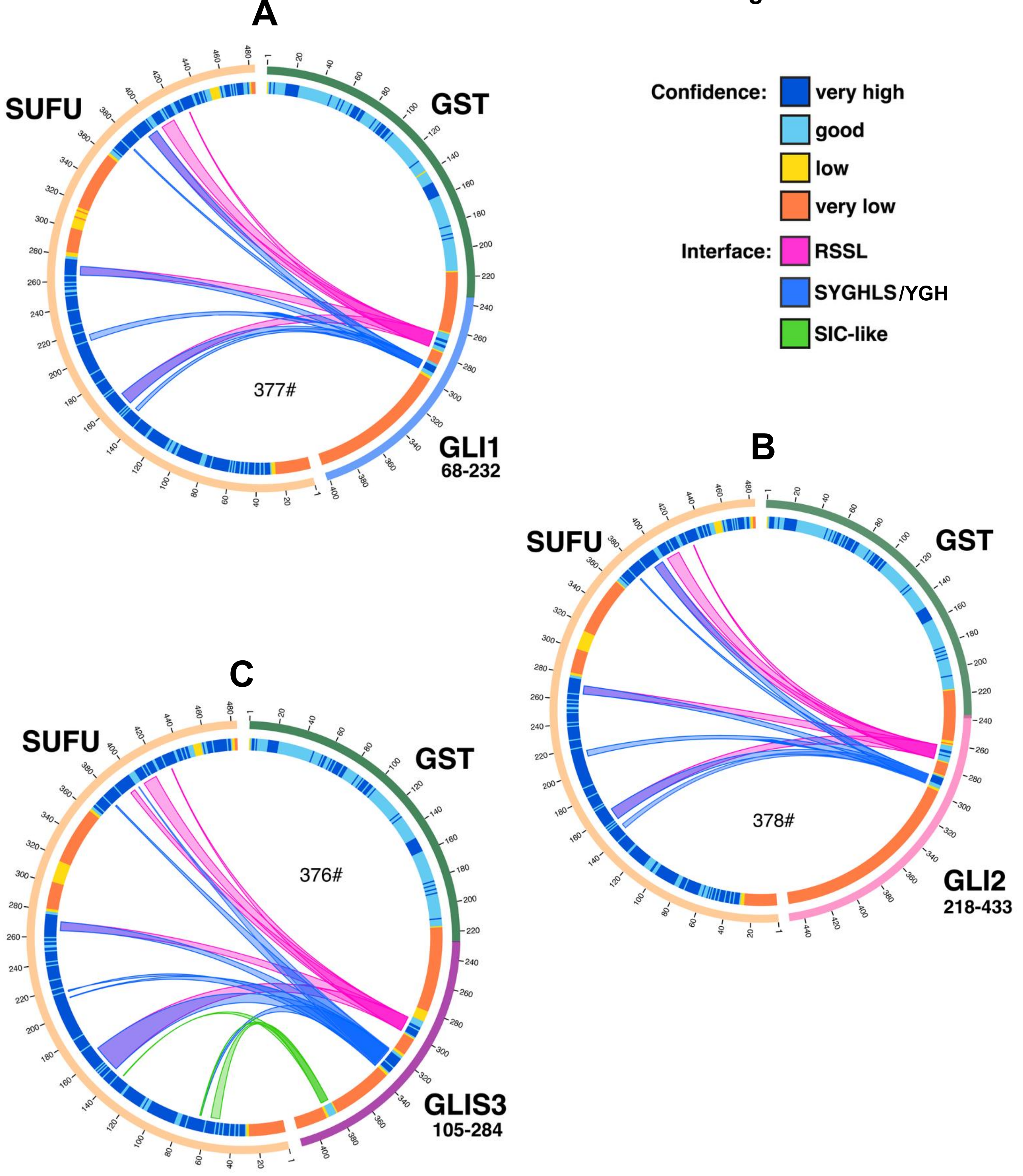


**Figure S25**. Chord diagrams of additional computed structure models:

A. CSM 377#, GST-GLI1$_{68-232}$ co-folded with full-length SUFU.

B. CSM 378#, GST-GLI2$_{218-433}$ co-folded with SUFU.

C. CSM 376#, GST-GLIS3$_{105-284}$ co-folded with SUFU.

Note the SIC $\beta$-strand-like interaction seen with GST-GLIS3$_{105-284}$ but not with the others, as well as the broadened YGH interface seen with GLIS3.
